# Conformational Variability of HIV-1 Env Trimer and Viral Vulnerability

**DOI:** 10.1101/2025.10.09.681376

**Authors:** Yiwei Cao, Wonpil Im

## Abstract

HIV-1 envelope glycoprotein (Env) is critical for viral fusion and entry into host cells and remains a primary target for vaccine and antiviral drug development. Advances in soluble gp140 trimer design have provided insight into the ectodomain structure and dynamics. While structural information is available for the membrane-proximal external region (MPER) and transmembrane domain (TMD), these regions remain comparatively understudied. Furthermore, high-resolution structural information for the cytoplasmic tail (CT), particularly within the context of the intact trimer, is limited and largely uncertain. Additionally, previous studies have typically treated the ectodomain and TMD as separate entities. To investigate the trimeric gp120–gp41 as a complete entity and its structural flexibility, we built a full-length model of the gp120–gp41 trimer that is fully glycosylated with N-linked glycans and embedded in a lipid bilayer, and performed all-atom molecular dynamics simulations. Our results show that the ectodomain maintains a rigid internal structure stable in the prefusion state, whereas the intrinsic flexibility of the MPER enables the ectodomain to adopt a range of tilted orientations, potentially enhancing spatial alignment for receptor engagement. The centrally positioned R696 residue in the TMD interacts with lipid headgroups, ions, and CT residues, resulting in conformational variability in the TMD and perturbations in the surrounding membrane that may facilitate the fusion process. Finally, we demonstrate how simulation trajectories can be leveraged to evaluate the accessibility of antibody epitopes across different regions of the protein.

## Introduction

Human immunodeficiency virus type 1 (HIV-1) is the most prevalent strain of HIV responsible for the development of acquired immunodeficiency syndrome (AIDS) (Sharp et al., 2011). The HIV-1 envelope (Env) consists of a host cell-derived lipid membrane and viral glycoproteins that play a crucial role in mediating viral entry into host cells. The Env glycoprotein is initially synthesized in the endoplasmic reticulum (ER) as a precursor gp160 and cleaved by furin into two subunits, gp120 and gp41. The non-covalently associated gp120–gp41 complex is transported to the cell surface in the form of a trimer, where it is subsequently incorporated into the envelope of nascent virions during viral assembly (Wyatt et al., 1998). Exposure of the Env protein is essential for binding to the primary receptor CD4 and co-receptors CCR5 or CXCR4, triggering membrane fusion (Dalgleish et al., 1984; Feng et al., 1996; Huang et al., 1996). However, this exposure also renders the virus susceptible to immune attack. In response to host immune pressure, Env is densely coated with N-linked glycans added during post-translational modification in the ER and Golgi apparatus, which effectively shield vulnerable epitopes from immune recognition (Wei et al., 2003).

Since HIV-1 was identified as the cause of AIDS in the early 1980s, extensive research has been conducted to elucidate the mechanisms of viral infection and immune response. A major advance was the development of soluble gp140 trimers, composing gp120 and the ectodomain portion of gp41, designed to stabilize the prefusion Env trimer for structural and immunological characterization. Native-like Env trimers, such as SOSIP (Sanders et al., 2013), native flexibly linked (NFL) (Sharma et al., 2015), and uncleaved prefusion-optimized (UFO) (Kong et al., 2016) constructs, mimic the structure of the viral spike and serve as valuable antigen targets for developing small molecule inhibitors and broadly neutralizing antibodies (bNAbs). With advances in structure determination techniques, such as X-ray crystallography and cryo-electron microscopy (cryo-EM), numerous high-resolution structures of gp140, both unliganded and antibody-bound, are now available in the Protein Data Bank (PDB). Despite this progress, the remaining portions of gp41, including the membrane-proximal external region (MPER), transmembrane domain (TMD), and cytoplasmic tail (CT), remain relatively understudied.

The MPER is a highly conserved region targeted by several bNAbs, including 10E8, 2F5, 4E10, and Z13e1 (Ofek et al., 2004; Cardoso et al., 2005; Pejchal et al., 2009; Huang et al., 2012). However, in most studies, the MPER is examined as a monomeric peptide bound to antibodies or as part of truncated Env constructs in membrane mimetics such as bicelles and nanodiscs (Rantalainen et al., 2020; Yang et al., 2022; Qi et al., 2025), which do not capture the structure and dynamics of the unliganded MPER in the context of an intact gp120–gp41 trimer embedded in a membrane bilayer. Despite the critical roles of the MPER and TMD in mediating the fusion of viral and host cell membranes (Salzwedel et al., 1999; Miyauchi et al., 2005), they are often excluded from structural studies due to difficulties in crystallization of hydrophobic TM segments while maintaining their native conformations. Therefore, nuclear magnetic resonance (NMR) spectroscopy remains one of the few viable methods for obtaining structural information of the MPER and TMD. However, NMR studies from different research groups have yielded conflicting conclusions regarding the oligomeric state of the TMD. Reported TMD structures include trimeric coiled coils (Dev et al., 2016; Kwon et al., 2018; Piai et al., 2020; Piai et al., 2021), monomeric helices (Apellaniz et al., 2015), and trimers that are not tightly bundled (Reardon et al., 2014).

Similarly, conclusions differ regarding the conformation and orientation of the MPER. Some studies suggest that gp41 bends at the MPER-TMD boundary (around residue 673), with the entire MPER adopting a helical conformation that lies nearly parallel to the membrane (Sun et al., 2008; Kwon et al., 2018), while others show that the C-terminal residues of the MPER form a continuous helix with the TMD (Dev et al., 2016; Chiliveri et al., 2018; Piai et al., 2020; Piai et al., 2021). In addition, the exceptionally long CT plays an important role in facilitating the incorporation of the Env glycoprotein into virions (Checkley et al., 2011). While the CT is known to contain three conserved amphipathic α-helical segments, referred to as lentiviral lytic peptides (LLPs), proposed models differ in the arrangement of LLPs (Murphy et al., 2017; Piai et al., 2021), and the complete structure of the CT and its location in the membrane remain inconclusive.

Molecular dynamics (MD) simulations have been employed to investigate the stability and conformational properties of both monomeric and trimeric TMD. An early study of the trimeric TMD established a foundational understanding of the domain’s stability, though it was limited by the computational resources available at the time (Kim et al., 2009). Subsequent work utilizing metadynamics found that the monomeric TMD is characterized by significant conformational plasticity and multiple metastable states, with the individual helix tilting in the bilayer and the midspan arginine (R696) interacting with lipid headgroups in either leaflet (Gangupomu et al., 2010; Baker et al., 2014). Baker et al. also simulated the monomeric TMD on Anton supercomputers, extended sampling to the multi-microsecond time scale, and demonstrated that TMD tilting and the interaction of R696 with lipids lead to local membrane thinning and water defects (Baker et al., 2014). Hollingsworth et al. modeled and simulated trimeric TMD in an asymmetric membrane and observed that TMD tilting and membrane thinning also occurred for the trimeric helical bundle, where water and ions permeated to stabilize the three positively charged R696 residues (Hollingsworth et al., 2018).

Piai et al. determined the NMR structure of a construct comprising the MPER, TMD, and CT, which currently serves as the only PDB structure to include the majority of the CT residues. They complemented this structural work with MD simulations to assess the structural stability of the trimeric MPER–TMD–CT complex (Piai et al., 2021). Recently, Majumder et al. simulated the same MPER–TMD–CT complex and applied a machine learning-based approach to classify the diverse conformational ensemble of the MPER-TMD-CT (Majumder et al., 2025). Maillie et al. combined conventional MD, steered MD, and coarse-grained simulations to demonstrate that interactions between MPER-targeting antibodies and membrane lipids are critical for effective epitope recognition (Maillie et al., 2025). In addition, MD simulations have been extensively applied to characterize the well-studied ectodomain.

Despite these advances, it remains challenging to investigate the gp120–gp41 trimer as an intact entity due to its structural complexity. In this work, we built a model of full-length gp120–gp41 trimer embedded in a lipid bilayer mimicking the lipid composition of the mammalian plasma membrane (van Meer et al., 2008; Sampaio et al., 2011; Ingolfsson et al., 2014; Pogozheva et al., 2022) (Figure 1). We chose this composition as a representative baseline, though we acknowledge that the native viral envelope may exhibit a distinct lipid profile that could influence protein-lipid interactions. Given the limited structural information and consequent uncertainty regarding the conformational organization of the CT, we also generated a CT-truncated model and used it as the primary system for analysis. While host furin cleavage of the gp160 precursor into gp120 and gp41 is a prerequisite for viral infectivity (McCune et al., 1988), native virions also incorporate a fraction of uncleaved gp160 (Zhang et al., 2021). Furthermore, many current immunogen designs, such as NFL and UFO constructs, utilize a covalent linker to stabilize the metastable prefusion conformation (Sharma et al., 2015; Kong et al., 2016). Therefore, we simulated both cleaved and uncleaved trimers to explore how the absence of proteolytic cleavage impacts the conformational landscape. We prepared simulation systems varying in the presence of the cleavage site and CT, as well as the initial position of protein in the membrane. Multiple microsecond-long all-atom MD simulations were performed for each system to explore the motions of individual protein domains and the membrane, and to examine how their conformational variability is affected by the difference in the initial configurations. Our results demonstrate that the ectodomain undergoes substantial tilting relative to the membrane plane while maintaining a rigid internal structure. In contrast, the MPER and TMD display highly diverse conformations, and their structural variations are influenced by the presence of the CT and the initial TMD position in the membrane. Moreover, by analyzing epitopes targeted by various bNAbs, we demonstrate that the simulation trajectories can be leveraged to assess the epitope accessibility.

**Figure 1.**
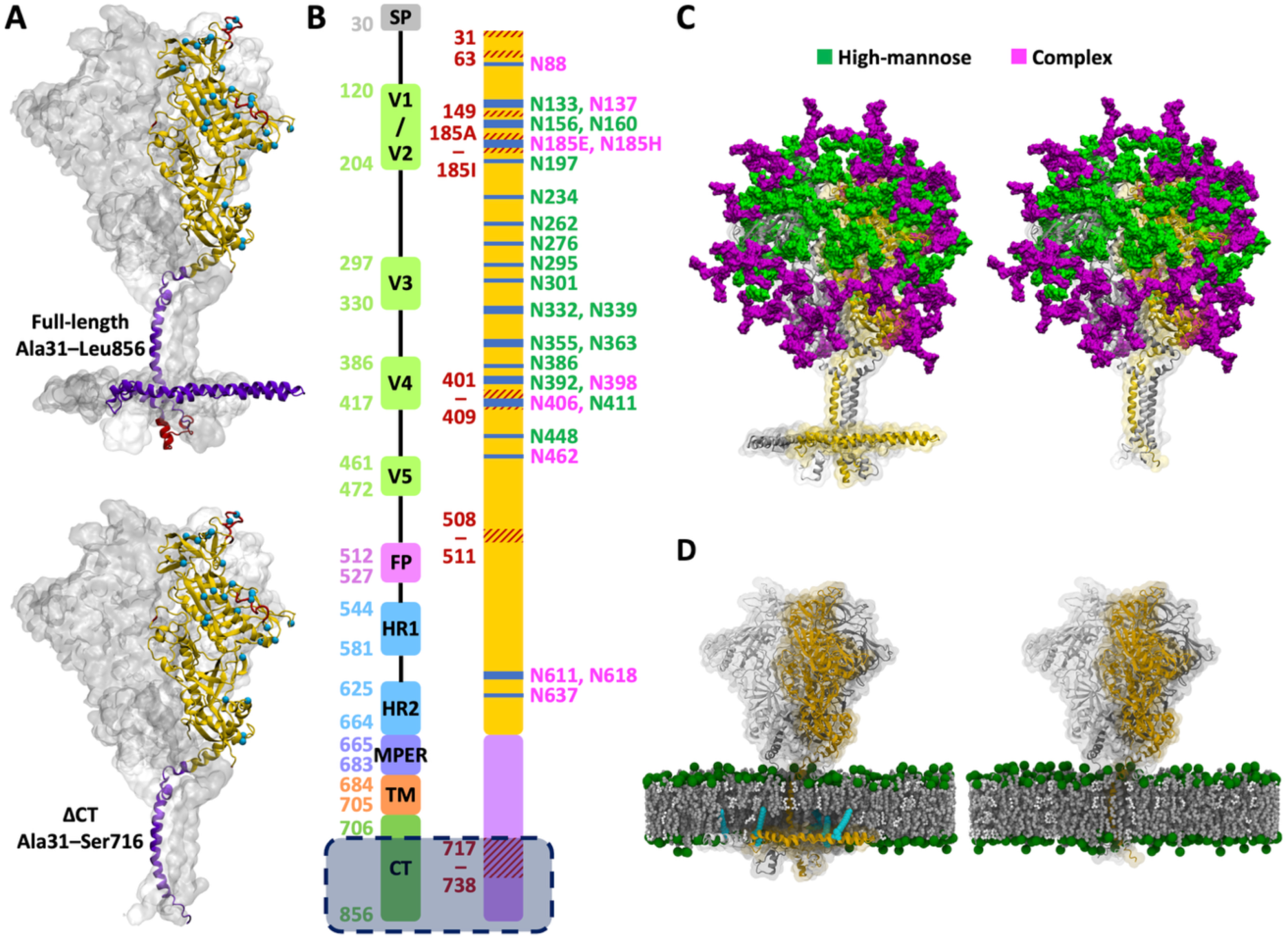
Model structure of a fully glycosylated full-length HIV Env trimer embedded in a membrane. (**A**) The model structure built by combining the cryo-EM structure of the ectodomain (yellow, PDB ID: 6B0N) with the NMR structure of the MPER, TMD, and CT (purple, PDB ID: 7LOI). The full-length model includes residues A31 to L856, while the CT-truncated (ΔCT) model includes residues A31 to S716. The missing loops in the PDB structures are highlighted in red, and the glycosylation sites are marked by cyan spheres. (**B**) Left: assignment of functional domains with boundary residue numbers, including signal peptide (SP), variable regions (V1-V5), fusion peptide (FP), heptad repeats (HR1 and HR2), membrane-proximal external region (MPER), transmembrane domain (TMD), and cytoplasmic tail (CT). Right: missing residues (red) and glycosylation sites (blue). The shaded region at the bottom marks CT residues excluded in the ΔCT model. (**C**) N-linked glycans shown as high-mannose (green) and complex (magenta) types. The full-length model is shown on the left and the ΔCT model on the right. (**D**) Env trimer embedded in a membrane. Lipid headgroups are highlighted by green spheres and glycans are omitted for visual clarity. The palmitoyl groups covalently attached to C764 and C837 are shown in cyan. Molecular illustrations were prepared using Visual Molecular Dynamics (VMD) (Humphrey et al., 1996).

## Results

### The ectodomain maintains a rigid internal structure and tilts independently of the TMD

The combination of cleavage state (cleaved vs. uncleaved), sequence length (full-length vs. CT-truncated), and initial TMD position in the membrane (high vs. low) resulted in eight distinct configurations, and we performed three independent 1-μs all-atom MD simulations for each configuration. Simulations trajectories are denoted as CH^CT^1, UL^CT^2, CL^ΔCT^3, UH^ΔCT^3, etc., where the first letter (C/U) indicates cleaved or uncleaved, the second letter (H/L) indicates the high or low initial TMD position, CT/ΔCT indicates the presence or absence of the CT, and the numeric suffix specifies the trajectory index among three replicas (Supplementary file 1–Supplementary Table 1).

In all simulations, both the ectodomain and TMD adopted variable orientations relative to the bilayer plane. To quantify the tilting of these two domains, we defined the tilt angles for the ectodomain (*θ*_EC_) and TMD (*θ*_TM_) (Figure 2A). *θ*_EC_ is measured between the bilayer normal and the vector from the center of mass (COM) of G594 (on all three protomers) to the COM of D167, while *θ*_TM_ is measured between the bilayer normal and the vector from the COM of I684 to the COM of V705. Across trajectories, *θ*_EC_ typically ranges from 0° to 40°, with only 0.7‰ exceeding 40°. Representative structures for different *θ*_EC_ values are shown in Figure 2A. In contrast to the wide range of *θ*_EC_, *θ*_TM_ generally remained within 0° to 20° with rare excursions to 30° observed in a small fraction of trajectories. Each 1-µs trajectory is divided into four consecutive 0.25-µs intervals, and data points from each interval are distinguished by four different colors (Figure 2—figure supplements 1–4). The variations of *θ*_EC_ and *θ*_TM_ over time show that large conformational changes predominantly occurred during the first 0.5 µs, followed by convergence of the *θ*_EC_ and *θ*_TM_ distributions during the second 0.5 µs in most trajectories. The temporal evolution of *θ*_EC_ and *θ*_TM_ is additionally shown in Figure 2—figure supplements 5–8.

**Figure 2.**
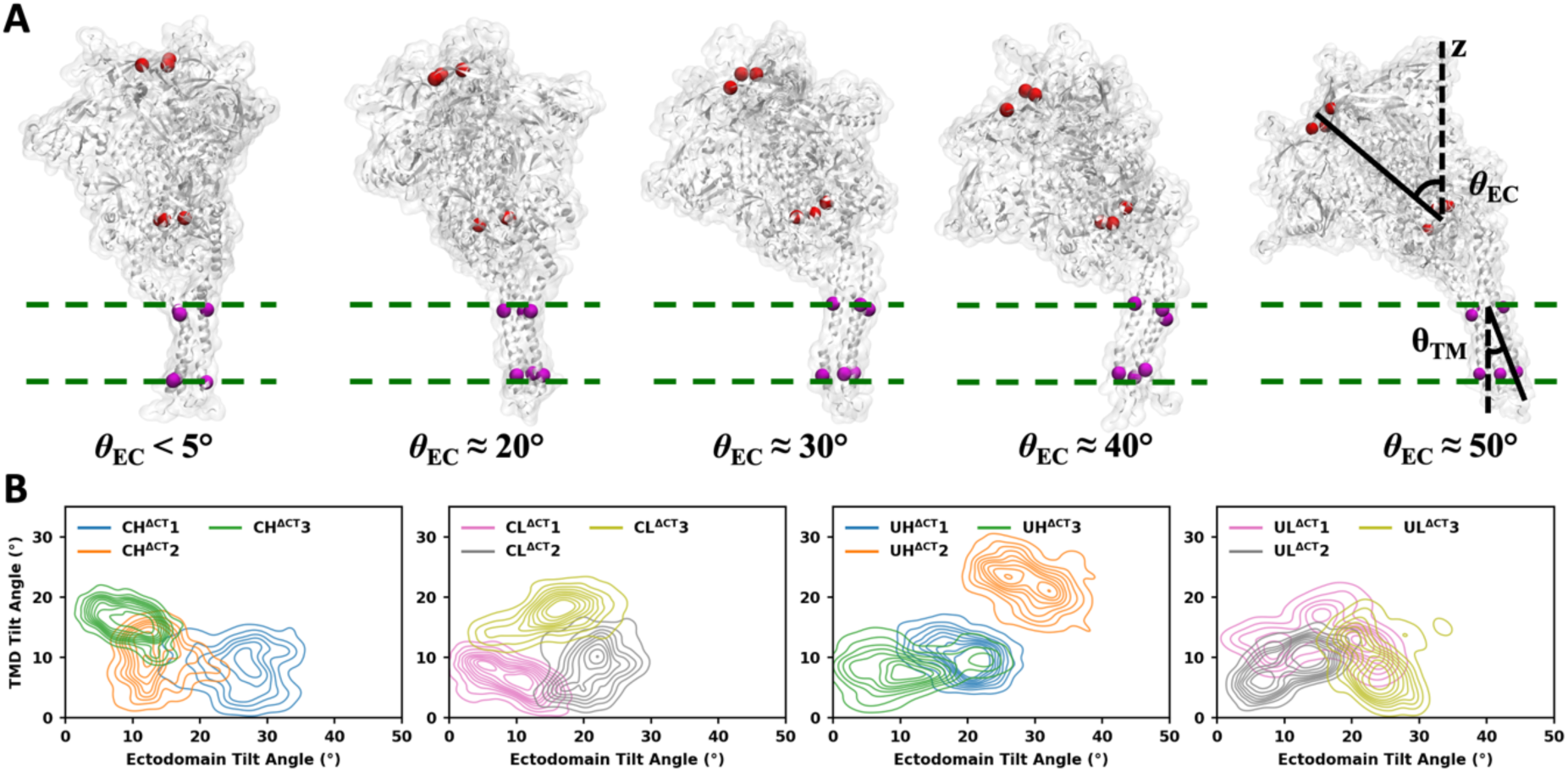
Tilting motions of the ectodomain and TMD are independent. (**A**) Representative structures illustrating different ectodomain tilt angles and the schematic showing how tilt angles are calculated. (**B**) Probability densities of ectodomain and TMD tilt angles, calculated from CT-truncated systems with various initial configurations.

For the CT-truncated systems, the joint probability densities of *θ*_EC_ and *θ*_TM_ calculated from the final 0.5 µs of each trajectory are shown in Figure 2B, while those for the full-length systems are shown in Figure 2—figure supplement 9. Although the combined dataset spans a broad range of *θ*_EC_ (0°–40°) and *θ*_TM_ (0°–30°), each individual trajectory explored only a portion of this space. For instance, CH^ΔCT^1 sampled large *θ*_EC_ with small *θ*_TM_, whereas CH^ΔCT^3 sampled small *θ*_EC_ with medium-to-large *θ*_TM_. Thus, multiple independent simulations are essential to capture a wide region of the conformational space. Notably, no consistent correlation was observed between *θ*_EC_ and *θ*_TM_, either within single trajectories or across all trajectories combined. Pearson correlation coefficients for *θ*_EC_ and *θ*_TM_ in single trajectories varied between -0.5 and 0.5, with examples such as CH^ΔCT^1 and CH^ΔCT^2 showing similar *θ*_TM_ but distinct θ_EC_ values, and CL^ΔCT^1 and CL^ΔCT^3 showing similar θ_EC_ but distinct *θ*_TM_ values. We also calculated the dynamical cross-correlation maps (Ichiye et al., 1991) of Cα atoms for all systems using CPPTRAJ (Roe et al., 2013). The results indicate only very weak correlations between the ectodomain and the TMD (Figure 2—figure supplements 10–13). Although Figure 2—figure supplements 12 and 13 also show low correlations between the ectodomain and the CT, we do not further interpret the coupling of the CT with the other domains due to its structural heterogeneity and the inherent uncertainty in its experimental structure.

Despite the considerable tilting of the ectodomain relative to the membrane, its internal structure remained rigid and well-preserved throughout the simulations. The root-mean-square fluctuation (RMSF) and root-mean-square deviation (RMSD) of the ectodomain were calculated after the snapshots from each trajectory were aligned to the initial structure by maximizing the overlap in the ectodomain. The majority of the ectodomain displayed low RMSF (< 2 Å), and the RMSD stabilized around 4 Å after an initial rise during the first 250 ns (Figure 3A, Figure 3—figure supplement 1). Higher RMSF values were observed only in the residues missing from the cryo-EM structure (PDB ID: 6B0N) (Sarkar et al., 2018), which was used as the template for the ectodomain in model building (these missing residues are highlighted in red in Figure 1A, B), and in part of HR1 (Q551-H564) that forms a flexible loop at the interface between two neighboring protomers.

**Figure 3.**
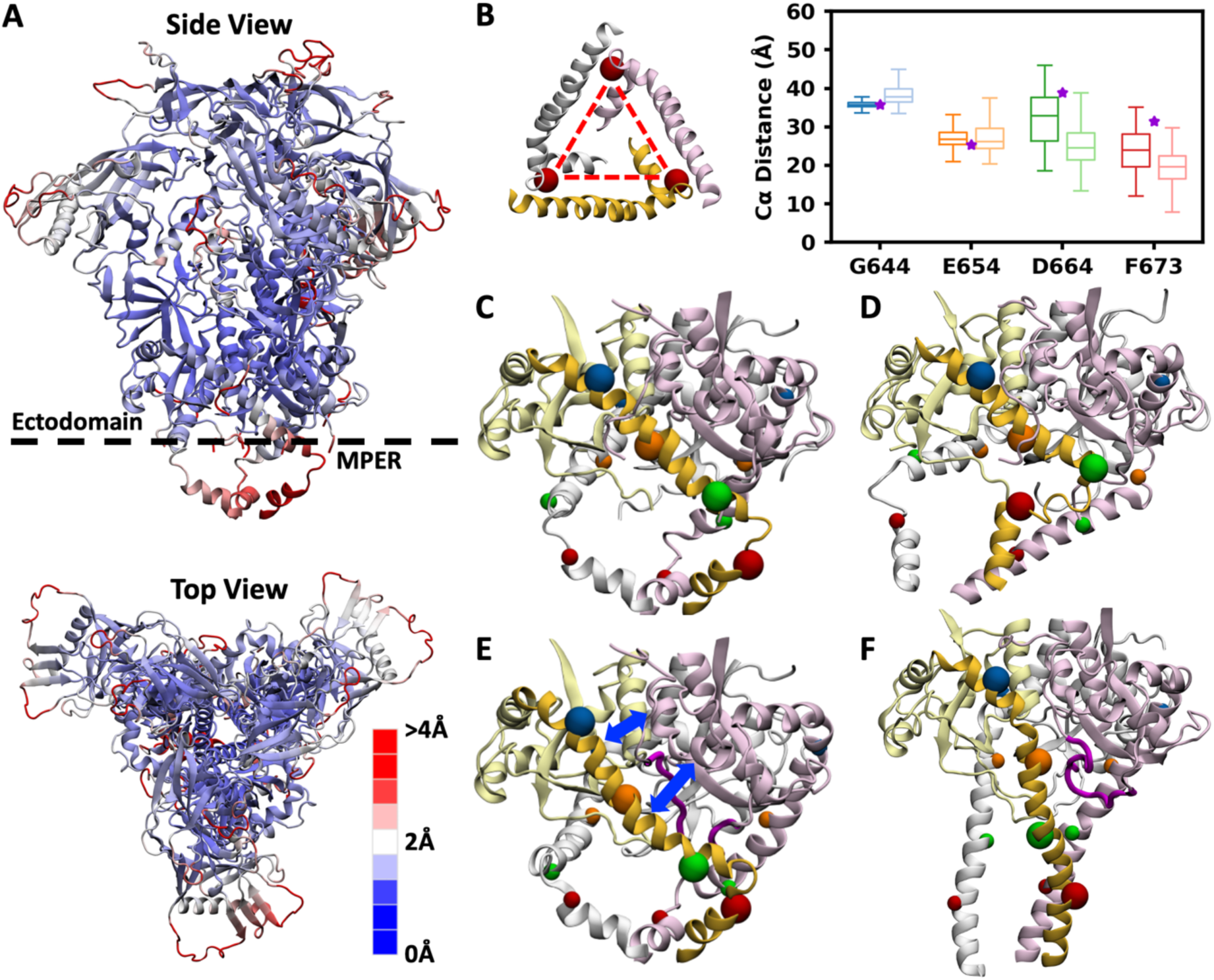
Ectodomain is rigid, whereas the MPER is highly flexible and adopts diverse conformations. (**A**) Top and side views of the ectodomain and MPER in the cleaved system, with RMSF indicated by color. (**B**) Schematic illustrating the calculation of interchain distances and their distributions at the Cα atoms of G644, E654, D664, and F673. For each residue, the distribution from cleaved systems is shown in dark color (left), and that from uncleaved systems is shown in light color (right). represented by solid and transparent colors, respectively. The initial values of interchain distances are marked by purple stars. (**C–F**) Local structures of the ectodomain C-terminus and MPER. The HR2 helix and MPER in one protomer are highlighted in dark yellow, with the Cα atoms of four selected residues marked by blue, orange, green and red spheres. (**C**) The initial conformation and (**D**) representative snapshot from simulations of the cleaved system. (**E**) The initial conformation and (**F**) representative snapshot from simulations of the uncleaved system.

In addition, the RMSF of the MPER was calculated with the trajectories aligned by the ectodomain. The entire MPER (K665–R683) and the adjacent heptad repeat 2 (HR2) segment (L660–D664) at the C-terminus of the ectodomain exhibited elevated fluctuations. The gp120–gp41 model was built based on the NMR structure in which the MPER adopts a bent conformation consisting of two helices joined by a sharp turn. The resulting trimer widens from the HR2 helix to the midpoint of the MPER (F673) and narrows from F673 to the TMD. However, this specific conformation was not maintained throughout the simulations. We measured the inter-chain distances between the Cα atoms of corresponding residues (G644, E654, D664, and F673) on neighboring protomers to characterize the structural variation along the HR2 helix and the MPER (Figure 3B). The inter-chain distances of G644 and E654 maintained narrow distributions centered on their initial values, while those of D664 and F673 exhibited broader distributions, reflecting inward shifts of three protomers. This effect was more pronounced in uncleaved systems. In cleaved systems, the HR2 helix interacted with residues M530–N543 and L619–N625 of neighboring protomers, which helped to stabilize the bent MPER conformation (Figure 3C, D). However, these interactions were not consistently observed across all protomers and all simulations; therefore considerable conformational variability remained in this region. In uncleaved systems, the closed loop at the cleavage site occupied the space between the HR2 helix and the neighboring protomer, disrupting their interactions and facilitating the inward shift of the HR2 helix (Figure 3E, F).

### The energetically unfavorable R696 in the hydrophobic core results in asymmetric, kinked TMD conformations and disrupts membrane integrity

Unlike the predominantly hydrophobic TMDs commonly found in many viral envelope proteins, the gp41 TMD contains multiple charged residues: R683 at the N-terminal boundary, R707 and R709 at the C-terminal boundary, and a central arginine, R696 (Figure 4A). In our simulations, R683 consistently interacted with lipid headgroups in the exoplasmic leaflet, while R707 and R709 interacted with those in the cytoplasmic leaflet, together acting as anchors that secured the TMD within the bilayer. In the NMR structure (PDB ID: 7LOI) (Piai et al., 2021), the side chain of R696 forms a cation-π interaction with the side chain of F699 and a hydrogen bond with the backbone carbonyl group of L692, but it is oriented outward from the helical bundle. When embedded in the bilayer, however, this configuration is energetically unfavorable, as the positively charged side chain directly contacts hydrophobic lipid tails (Cheng et al., 2012). Early in the equilibration stage, the TMD rapidly rearranged to allow R696 residues to interact with more favorable partners, including negatively charged lipid headgroups from either leaflet, ions and water molecules diffusing into the bilayer center, as well as polar and positively charged groups in the CT when present. Once the interactions between R696 residues and their binding partners (lipid headgroups, ions or CT residues) were established, they remained stable with minimal changes throughout the production stage. Because the limited space at the TMD core can only accommodate at most two inward-facing arginine residues, at least one R696 was forced outward to interact with lipid headgroups or CT residues. The differences in arginine orientation and interacting partners gave rise to asymmetric protomer conformations and distinct TMD tilts (Figure 4A–F). When R696 pointed outward, its interactions with lipid head groups or CT residues were strong enough to destabilize the local helix, introducing a kink into the TMD. Representative snapshots from different trajectories illustrate these asymmetric kinked conformations (Figure 4—figure supplements 1–8).

**Figure 4.**
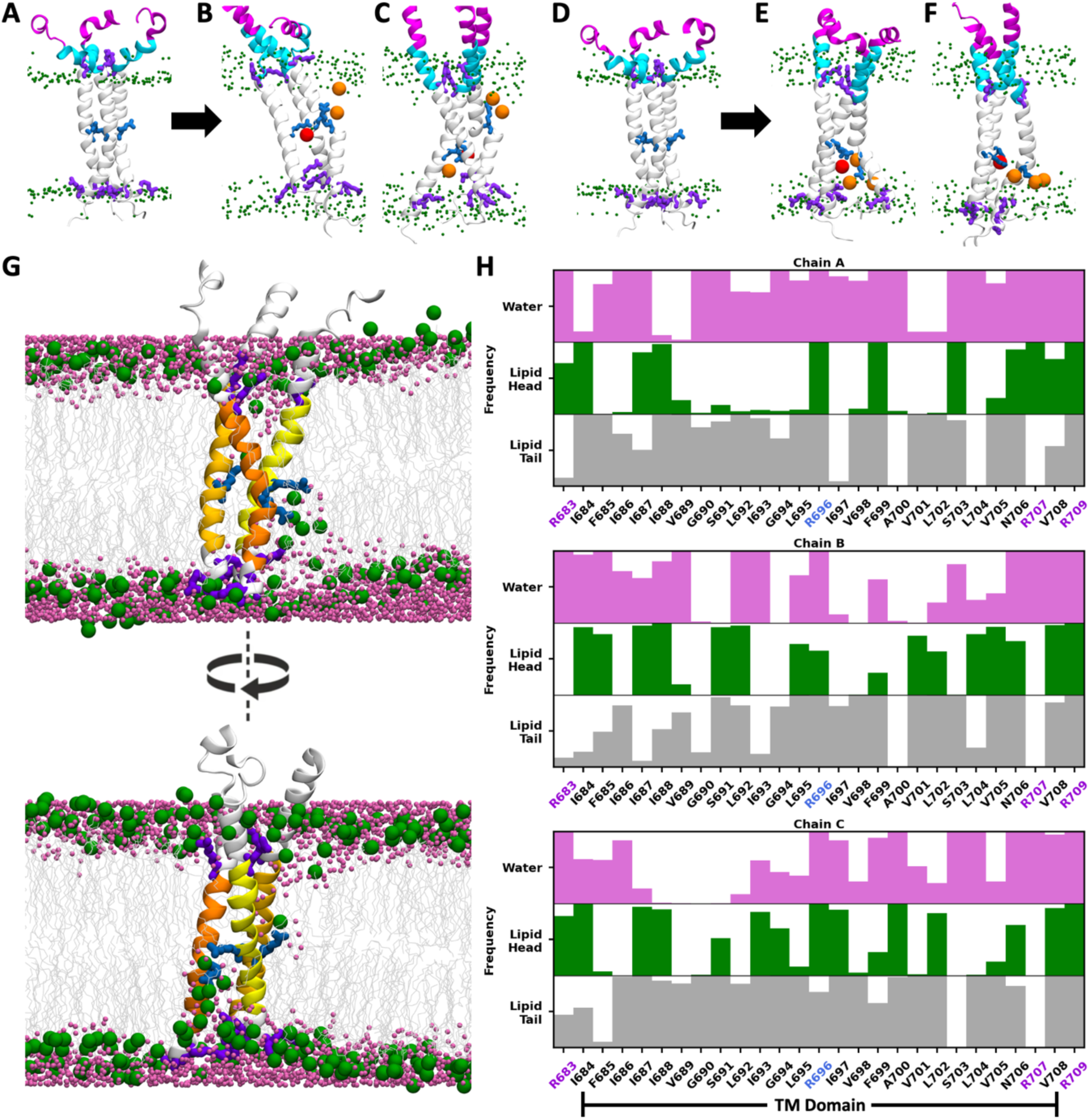
R696 interacts with lipid headgroups and disrupts membrane integrity. (**A–C**) MPER and TMD in the CT-truncated system with the “high” TMD configuration. MPER-N, MPER-C, and TMD are shown in magenta, cyan, and white, respectively. Lipid headgroups, R696, and the residues anchored in the lipid headgroups (R683, R707 and R709) are shown in green, blue, and purple, respectively. Lipid headgroups and ions interacting with R696 are highlighted in orange and red, respectively. (**A**) Initial conformation. (**B, C**) Representative snapshots from different trajectories. (**D–F**) MPER and TMD in the CT-truncated system with the “low” TMD configuration. (**G**) Two side views of the same snapshot where R696 of one protomer interacts with lipid headgroups in the exoplasmic leaflet and R696 of two protomers interact with lipid headgroups in the cytoplasmic leaflet. Lipid headgroups and tails are shown in green and gray, and water molecules in magenta. TMD of three protomers (i.e., chains A, B and C) are shown in light yellow, dark yellow and orange, respectively. (**H**) Frequency of TMD residues interacting with lipid headgroups, lipid tails, and water. For each TMD residue–interacting component pair, the frequency represents the fraction of snapshots in which the heavy atoms of the TMD residue and the corresponding component are within 5 Å. Bar shading reflects this fraction, with fully filled bars indicating 100% and empty bars indicating 0%.

Beyond local deformation of the TMD, R696–lipid interactions perturbed bilayer organization, inducing the translocation of lipid headgroups and water molecules toward the bilayer center (Figure 4G). We calculated the interaction frequencies of each TMD residue with lipid headgroups, lipid tails, and water molecules. The results show that the membrane disruption is persistent as many non-terminal TMD residues maintained frequent contacts with water and lipid headgroups throughout the simulations (Figure 4H). Previously, Kim et al. reported that the inter-chain interactions between protonated R696 gradually diminished over a short simulation time (23 ns), leading to increased crossing angles and reduced bundle length (Kim et al., 2009). Gangupomu et. al and Baker et. al observed that R696 snorkeled toward either exoplasmic or endoplasmic headgroups in simulations of the TMD monomer, resulting in TMD tilting and membrane thinning due to water penetration and lipid headgroups interacting with R696 (Gangupomu et al., 2010; Baker et al., 2014; Baker et al., 2014). These observations are consistent with our findings. Hollingsworth et. al also reported membrane thinning; however, they attributed this effect to interfacial interactions of R683 and R707 with both leaflets and proposed that R696 only interacted with water and ions permeating into the center of the TMD timer (Hollingsworth et al., 2018).

To explore whether R696 exhibits a preference for the exoplasmic versus cytoplasmic leaflet, or interacts with either randomly, we generated two initial structures (“high” and “low”) with the TMD positioned at two distinct positions, separated by 4 Å along the membrane normal (see Methods). In simulations initiated from the “high” TMD configuration, R696 residues in three protomers interacted with the lipid headgroups in either leaflet (Figure 4B, C, Figure 4—figure supplements 1, 3) In contrast, in simulations initiated from the “low” TMD configuration, R696 residues interacted exclusively with the headgroups in the cytoplasmic leaflet (Figure 4E and F, Figure 4—figure supplements 2, 4). In the full-length systems, the plate-shaped CT occupies a substantial space within the cytoplasmic leaflet, displacing lipids during model construction. Because the CT is not thick enough to fully span the cytoplasmic leaflet, an empty gap remained between the CT and the exoplasmic leaflet (Figure 1D). Over time, lipids in the exoplasmic leaflet shifted downward while the CT residues moved upward to fill this space, causing local bilayer thinning. Under these conditions, upward-oriented R696 can still interact with the headgroups in the exoplasmic leaflet, but downward-oriented R696 primarily contacted CT residues, with rare cases of lipids migrating upward from the cytoplasmic leaflet to approach the protein. A comparison of all full-length systems (Figure 4—figure supplements 5–8) shows that R696 preferentially adopted downward orientations in the simulations initiated from the “low” TMD configuration (Figure 4—figure supplements 6, 8). In these cases, upward-shifted CT residues interacted with the C-terminal half of the TMD, rather than solely with R696, resulting in deeper burial of the TMD in the membrane. Taken together, these observations suggest that interactions of R696 with lipid headgroups and CT residues may modulate TMD tilt and kink formation during viral entry. However, whether the orientation of R696 dynamically switches between the two leaflets over longer timescales, and whether a preference exists for either leaflet, remain to be examined in future experimental and/or enhanced sampling simulation studies.

### MPER adopts diverse conformations, and its exposure depends on both MPER and TMD conformations

Starting from the initial helix-turn-helix conformation consisting of two separate helical segments, the N-terminal half (MPER-N) and the C-terminal half (MPER-C), MPER underwent rapid rearrangements, and a wide variety of conformations were sampled across all trajectories from all systems. In the initial structure, the trimeric MPER was positioned perpendicular to the membrane, with the helical MPER-N tilted inward and MPER-C tilted outward (Figure 4A). Such conformation and orientation were maintained in some trajectories such as CL^ΔCT^3 (the third trajectory of the cleaved, CT-truncated system with the low TMD position, Figure 4—figure supplement 2C). In other trajectories, such as CL^CT^1, the helix-turn-helix MPER in one protomer shifted into a horizontal orientation parallel to the membrane surface (Figure 4—figure supplement 6A). In UL^ΔCT^1, the entire MPER adopted a more vertical arrangement, with both MPER-N and MPER-C tilted outward (Figure 4E, Figure 4—figure supplement 4A). We also observed in UH^ΔCT^3 and UL^ΔCT^3 that the HR2 helix in the ectodomain, MPER, and TMD merged into a continuous long helix (Figure 4C, F, Figure 4—figure supplements 3C, 4C). In addition, loss of helical structure within the MPER was common, particularly in the MPER-C region, which often transitioned to a random coil. Therefore, the distinct MPER conformations reported in various experimental studies can be all valid as each captures a possible state within the highly flexible conformational landscape of the MPER. It is noteworthy that the MPER in three protomers can adopt different conformations and orientations, resulting in asymmetric local structures.

Because the MPER is a target of multiple bNAbs, we next examined how its exposure is affected by TMD conformation. As described above, interactions between R696 and lipid headgroups affect the burial depth of the TMD, thereby influencing the positioning of the adjacent MPER. To quantify MPER exposure, we measured the vertical distance from the Cɑ of F673, approximately the midpoint of the MPER, to the highest point of the neighboring lipid headgroups, denoted by *d*_F673_. Positive values indicate that F673 lies above the bilayer surface, and negative values indicate its membrane burial. In the initial “low” and “high” TMD configurations, *d*_F673_ was 6.1 Å and 9.1 Å, respectively, but it spanned a wide range from -15 Å to 20 Å across simulations (Figure 5A, B, Figure 5—figure supplement 1). Two examples illustrate this variability. In the first example, all three R696 residues interacted with the cytoplasmic leaflet, drawing the TMD deeper into the membrane. Consequently, the entire MPER-C and most of MPER-N were buried in the membrane, with one F673 positioned 11.3 Å below the membrane surface, thus bringing the ectodomain in close proximity to the membrane (Figure 5C, E). In the second example, two R696 residues interacted with the exoplasmic leaflet, while one interacted with the cytoplasmic leaflet. In the protomer with upward-oriented R696, the MPER-N, MPER-C, and TMD formed a continuous helix. As a result, most of the MPER extended outside the bilayer, with *d*_F673_ reaching 18.0 Å and the ectodomain displaced farther from the membrane (Figure 5D, F).

**Figure 5.**
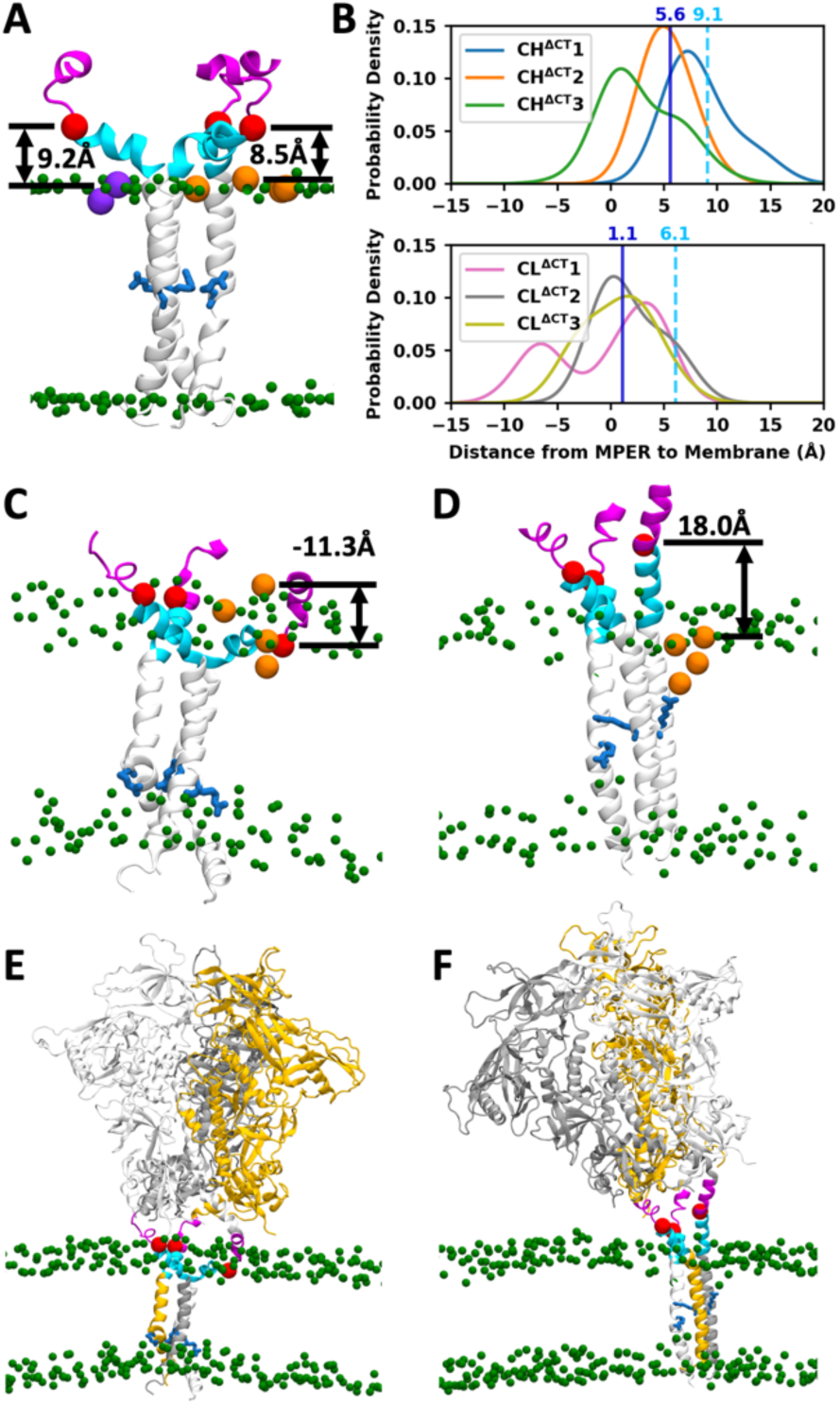
MPER exhibits diverse conformations, and its exposure depends on both MPER and TMD. (**A**) The initial structure of the CH^ΔCT^ system, where *d*_F673_ of two promoters equals 8.5 Å and 9.2 Å. Lipid headgroups are shown in green and R696 in blue. *d*_F673_ is defined as the distance from the Cɑ of F673 (red) to the highest among the adjacent lipid headgroups (orange and purple). (**B**) Distribution of *d*_F673_ in the CL^ΔCT^ and CH^ΔCT^ systems. The cyan dashed line indicates the mean *d*_F673_ of three protomers in the initial structure, and the blue solid line indicates the mean across all data sampled from simulations. (**C, D**) Representative snapshots illustrating the buried (**C**) and exposed (**D**) MPER. (**E, F**) The entire trimer structures corresponding to (**C**) and (**D**), respectively.

The *d*_F673_ distributions across all cleaved CT-truncated systems suggest that *d*_F673_ tends to be smaller when the simulations start from the “low” TMD configuration (Figure 5B). In both “high” and “low” configurations, the mean *d*_F673_ calculated from the simulation trajectories is smaller than its value in the initial structure, due to both protein conformational changes and lipid diffusion. In the “low” configuration, *d*_F673_ decreased by 5.0 Å (from 6.1 Å to 1.1 Å), a slightly larger reduction than in the “high” configuration where *d*_F673_ dropped by 3.5 Å (from 9.1 Å to 5.6 Å). This indicates that deeper MPER burial arises from a larger-scale protein motion rather than merely from a lower initial placement.

### Ectodomain epitopes are conditionally accessible, whereas MPER epitopes are virtually inaccessible in the closed prefusion state

To access the accessibility of epitopes on different regions of the trimeric Env protein in the prefusion state, we quantified how frequent their epitopes were exposed without steric clashes from neighboring protein residues, glycans or membrane lipids. We examined six selected antibodies: PGT128 targeting the V3 loop and the N332 glycan (PDB ID: 5JSA) (Kong et al., 2016); PG9 targeting the V1/V2 loop (PDB ID: 3U2S) (McLellan et al., 2011); VRC01 targeting the CD4 binding site (PDB ID: 4LST) (Zhou et al., 2013); 35O22 targeting the gp120–gp41 interface (PDB ID: 4TVP) (Pancera et al., 2014); and 10E8 (PDB ID: 6VPX) (Rantalainen et al., 2020) and 4E10 (PDB ID: 1TZG) (Cardoso et al., 2005) targeting the MPER (Figure 6A, Supplementary file 1–Supplementary Tables 2–5).

**Figure 6.**
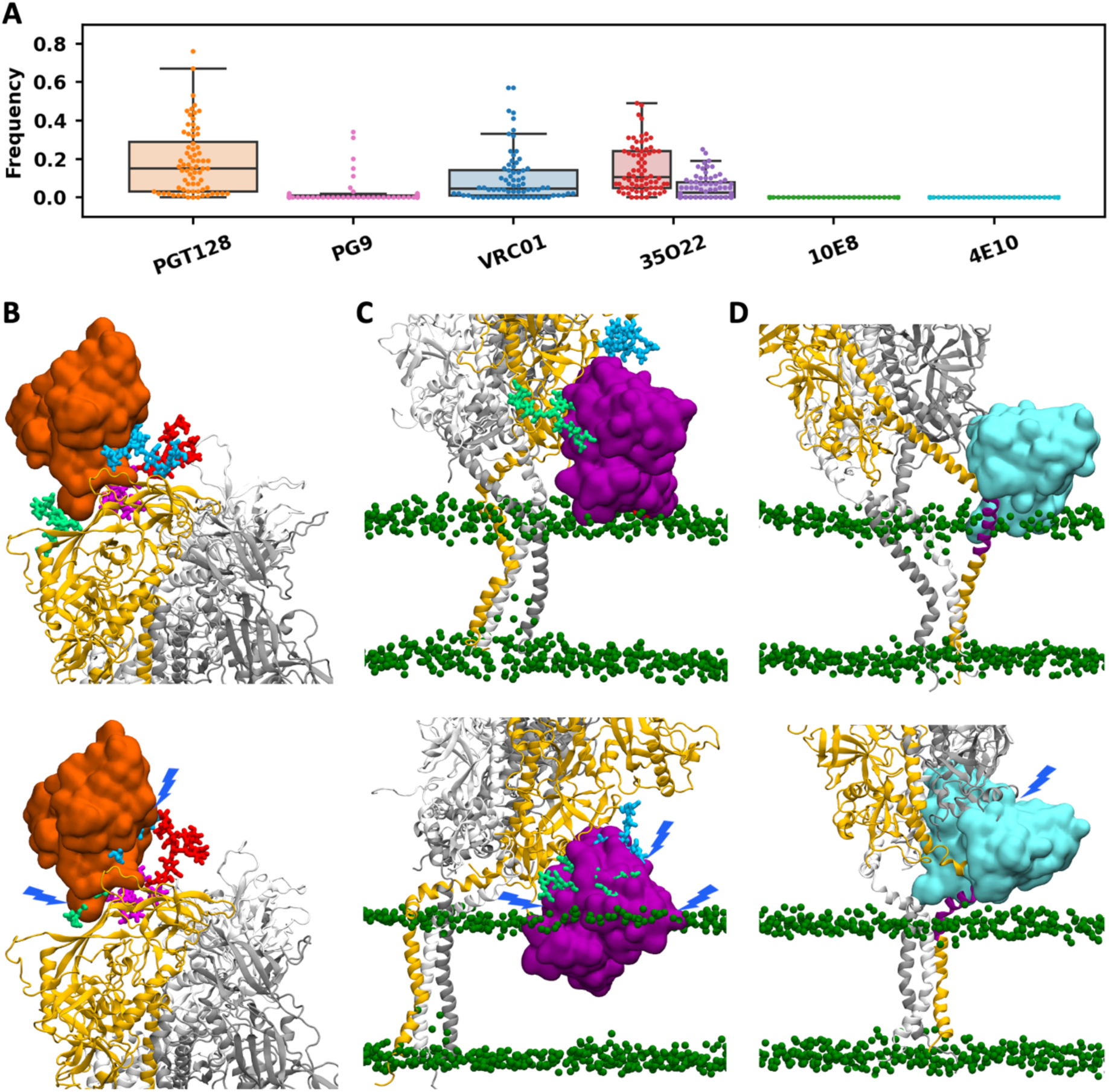
Antibody epitope accessibility. (**A**) The frequency of accessibility. Each marker represents the epitope on one of the three protomers across all trajectories. For 35O22, red indicates the accessibility frequency without considering steric clashes with the membrane, while purple indicates the frequency accounting for clashes with the membrane. (**B–D**) Representative snapshots showing conformations with the epitope exposed (upper) and shielded (lower) for antibodies PGT128, 35O22, and 4E10, respectively. The antibody V_H_ and V_L_ domains are shown in surface representation, with lipid head groups in green spheres and glycans that may interfere with the antibody in distinct colors.

The epitope of PGT128 comprises the V3 loop and the N332 glycan. It is widely recognized that PGT128 binding is mediated by the protein-protein interaction with the V3 loop and the protein-glycan interaction with the N332 glycan, facilitated by its extra-long HCDR3 loop penetrating the glycan shield (Figure 6—figure supplement 1). Due to the absence of defined secondary structures, glycans exhibit greater conformational flexibility than the protein, although the crowded surface of gp120–gp41 may partially restrict the motion of glycans. The N332 glycan must adopt specific conformations to enable interactions with PGT128, whereas most conformations block antibody approach. Additional glycans on N137, N156, and N301 can further occlude the site (Figure 6B). Epitope accessibility for PGT128 varied widely across protomers and trajectories, with many cases exceeding 35%. When considering all trajectories from all systems collectively, approximately half of them exhibited at least one protomer with >35% accessibility (Supplementary file 1–Supplementary Table 2). For instance, in trajectory CH^ΔCT^1, two protomers showed accessibility >40%, whereas in CL^CT^2, all three protomers are <5%. The high accessibility frequency observed for the PGT128 epitope aligns with its exceptional potency. As demonstrated by Walker et al., PGT128 is capable of neutralizing approximately 72% of global isolates with a median *IC*_50_ of ∼0.02 µg/mL. This potency is approximately 10-fold greater than that of PG9 and VRC01, though its breadth is lower than the 93% reported for VRC01 (Walker et al., 2011). This comparatively lower breadth may be attributed to strict sequence dependency. Because PGT128 recognition depends on the N332-centered glycan epitope, loss, truncation, or shifting of the N332 glycan to N334 prevents productive engagement regardless of local steric accessibility.

The second antibody, PG9, targets the V1/V2 apex, where binding can be hindered by six glycans, N156, N160, and N185E on the same protomer, and N160, N185E, and N185H on the neighboring protomer (Figure 6—figure supplement 2). In most trajectories, the epitopes on all three protomers were occluded in >95% of the snapshots, with a few exceptions (CH^ΔCT^1, CH^ΔCT^2, and CL^ΔCT^3) where accessibility was non-negligible (Supplementary file 1–Supplementary Table 3). This is consistent with the lower neutralization potency and moderate breadth of PG9, which exhibits a median *IC*_50_ of ∼0.22 µg/mL and a breadth of ∼79% (Walker et al., 2009).

The third antibody, VRC01, targets the CD4 binding site where six glycans, N185H, N197, N276, N363, and N462 on the same protomer and N301 on the neighboring protomer, can interfere with antibody binding (Figure 6—figure supplement 3). The VRC01 epitope is moderately to highly accessible on at least one protomer in many trajectories, whereas in some trajectories, the epitopes on all three protomers are nearly completely shielded (Supplementary file 1–Supplementary Table 4). This intermediate accessibility is consistent with the biological requirement for the CD4 binding site to remain periodically available for receptor engagement while maintaining a certain degree of glycan shielding to evade neutralization. The potency of VRC01 is even lower than that of PG9, with a reported median *IC*_50_ of ∼0.32 µg/mL, but it possesses an exceptionally high breadth of ∼93% (Wu et al., 2010; Walker et al., 2011).

35O22 targets the gp120–gp41 interface that is not intrinsically membrane-proximal. However, its binding orientation combined with ectodomain tilting can lead to steric clashes between the antibody and the membrane (Figure 6—figure supplement 4). Comparing accessibility with and without including the steric effects of membrane lipids shows that the membrane can substantially hinder 35O22 binding, particularly when the antibody approaches from the direction of ectodomain tilt (Figure 6C). For example, in system UH^ΔCT^2, the accessibility frequencies of the three protomers were 49%, 5%, and 25%, when only the shielding of three glycans (N88, N234, and N618) was considered. When the steric effects of both glycans and membrane lipids were included, the first protomer’s frequency dropped from 49% to 1%, while the other two remained essentially unchanged (Supplementary file 1–Supplementary Table 5). Altogether, these results demonstrate that epitope accessibility for this antibody is highly sensitivity to the membrane environment, glycan orientation and ectodomain tilting. This complex dependency provides a structural context for the experimental profile of 35O22, which exhibits high potency with a median *IC*_50_ of ∼0.03 µg/mL, but a relatively limited breadth of ∼62% (Huang et al., 2014).

The remaining two antibodies, 10E8 and 4E10, target the MPER. In the PDB structures of 10E8, the epitope is a 17-residue peptide corresponding to residues 671-687 in this work. In the PDB structure of 4E10, the epitope is a 12-residue peptide corresponding to residues 669-680. Superposition of these antibodies onto the initial simulation structure revealed extensive clashes with the protein and glycans linked to N611, N618, and N637 on the neighboring protomer, as well as slight steric clashes with the membrane (Figure 6—figure supplements 5, 6). Using the strict criteria (≤10 heavy-atom clashes with protein/glycan and ≤20 with lipids), no snapshot was identified in which the epitope of either antibody was accessible. Even with relaxed criteria (≤20 heavy-atom clashes with protein/glycan or ≤40 with lipids), accessible cases remained rare, which require either large ectodomain tilts to create space on the opposite side or the MPER transitioning into continuous helices to displace the ectodomain from the membrane (Figure 6D, Figure 6—figure supplements 7–10). Given the extremely low frequency of such specific conformations, MPER epitopes are effectively inaccessible in the prefusion trimer. Though differing in potency — with 10E8 exhibiting a median *IC*_50_ of ∼0.35 µg/mL compared to ∼1.93 µg/mL for 4E10 — both antibodies demonstrate extremely high breadth of ∼98% (Huang et al., 2012). This extensive breadth is primarily attributed to the high sequence conservation of the MPER across global isolates. The negligible epitope accessibility observed in the prefusion trimer supports the conclusion that these antibodies require the transition of the Env trimer into intermediate states to fully engage their epitopes (Frey et al., 2008).

## Discussion

The hydrophobic and flexible nature of the MPER has made it difficult to crystallize, and most structural insights have come from NMR studies. However, structural information for the MPER and TMD has been inconsistent in the literature. For example, one study of the MPER in DPC micelles suggested that it adopted a distorted helical structure lying parallel to the membrane surface (Sun et al., 2008), while NMR studies using gp41 fragments containing part of the MPER and TMD embedded in bicelles or bilayers reached conflicting conclusions. Some proposed a kink (Apellaniz et al., 2015) or a turn (Kwon et al., 2018) between the MPER and TMD, whereas others suggested that the MPER and TMD formed a continuous helix aligned perpendicular to the membrane (Dev et al., 2016; Chiliveri et al., 2018). In the NMR structure (PDB ID: 7LOI) used to build our model structure, a kink appears between the MPER and TMD, along with a turn in the middle of the MPER (Fu et al., 2018; Piai et al., 2021). Crystal structures of MPER peptides bound to various antibodies show that in addition to the helix-turn-helix conformation, the MPER can also adopt alternative structures in which the MPER-C remains helical while part of the MPER-N becomes unstructured (Williams et al., 2017). Such variability underscores the intrinsic structural flexibility of the MPER, with each experimental method likely capturing only a subset of possible states under specific conditions. In this study, we employed all-atom MD simulations to sample a broader conformational landscape and gain a more comprehensive view of the structural heterogeneity of the MPER. Our results reveal that the MPER can adopt virtually all of the conformations reported experimentally. This is consistent with the all-atom simulations of MPER–TMD–CT and MPER–TMD in an asymmetric membrane conducted by Majumder et al., which likewise demonstrated multiple conformational states, including helix-turn-helix and extended helical conformations (Majumder et al., 2025). While MPER plasticity has been linked to its role in virus-host membrane fusion because it enables the ectodomain and TMD to adopt distinct orientations during large-scale structural rearrangements (Salzwedel et al., 1999), our results show that this flexibility is already inherently present in the prefusion state. At this stage, the MPER functions as a flexible hinge that facilitates ectodomain tilting, which can be critical for the spatial alignment of the CD4-binding site with the host receptor for efficient receptor engagement.

Comparison of the full-length and CT-truncated systems shows that the primary difference arises from changes in the lipid bilayer, particularly in the exoplasmic leaflet, whereas differences in protein conformation and dynamics are less evident. Previous experimental studies have reported that mutations of TMD residues and CT truncation can substantially affect the antigenicity of the ectodomain (Edwards et al., 2002; Chen et al., 2015; Dev et al., 2016). However, the ectodomain remains relatively rigid in our simulations for both full-length and CT-truncated systems. It is unclear whether this behavior reflects insufficient conformational sampling or potential artifacts associated with the model structures. Structural information for the CT is very limited, and the NMR structure (PDB ID: 7LOI) was the only available CT structure at the time the simulation systems were constructed. As a result, the extent to which this structure represents the native CT conformation remains uncertain. Additional experimental structural characterization of the CT will be important for achieving a more complete understanding of its functional role. Similar limitations apply to other modeled regions where structural information is incomplete, including missing loops in the ectodomain, the cleavage site and the HR2 region where two PDB structures (IDs: 6B0N and 7LOI) were merged. These regions introduce additional uncertainty, and the extent to which they influence the interpretation of our results remains an open question.

Our analysis of the ectodomain and TMD coupling indicates that the motions of these two domains are largely independent. This observation does not contradict experimental studies demonstrating functional coupling between the TMD, CT, and the antigenic profiles of Env (Chen et al., 2015; Dev et al., 2016). Munro et al. proposed that unliganded Env is intrinsically dynamic, transitioning among three distinct prefusion conformations: a closed ground state (predominant), a transient state, and a CD4-/co-receptor-stabilized state. Both laboratory-adapted and clinically isolated strains can spontaneously transition among these three states, although their relative occupancies differ (Munro et al., 2014). It is therefore possible that TMD mutations or CT truncation alter the equilibrium distribution among these states, thereby affecting the epitope exposure, particularly for epitopes that are occluded in the closed ground state while exposed in the CD4-/co-receptor-stabilized state. However, transition among these three states occur on millisecond-to-second timescales (Munro et al., 2014). Our simulations on microsecond timescales primarily capture conformational variations within the closed ground state and suggest that the MPER acts as a hinge, providing substantial flexibility that enables the ectodomain and TMD to move independently while Env remains in the closed ground state. Enhanced sampling methods could be applied to more thoroughly explore the conformational landscape, including not only variations within the closed ground state but also transitions among the closed ground, transient and CD4-/co-receptor-stabilized states.

In addition to the limitations inherent to protein structure modeling, the choice of lipid composition remains an open question. In this work, we selected an asymmetric mammalian plasma membrane because it is one of the 18 complex biomembrane systems we previously studied (Pogozheva et al., 2022), and among them, it provides the closest available approximation to the HIV membrane. Nevertheless, experimental studies have reported differences in lipid composition between HIV virions and the host plasma membrane (Aloia et al., 1993; Brugger et al., 2006; Huarte et al., 2016; Mucksch et al., 2019; Tomishige et al., 2023). Although we do not anticipate that our main conclusions regarding Env domain motions and MPER flexibility would change substantially, evaluating the influence of lipid composition represents an important direction for future work.

To probe antibody recognition, we assessed epitope accessibility using snapshots extracted from simulation trajectories. Unlike static experimental structures, this approach incorporates protein dynamics and the influence of glycans and membrane lipids, yielding a frequency-based measure of how often each epitope is accessible rather than a binary classification. It is worth noting that viral glycans vary highly in both site occupancy and the specific glycoforms present at each glycosylation site, as indicated by mass spectrometry data (Behrens et al., 2016; Cao et al., 2017). For structure modeling, however, we assumed full occupancy at all glycosylation sites and selected a single representative glycoform per site. As a result, the estimated glycan shield may not fully reflect the *in vivo* situation. Furthermore, this analysis is based on geometric and steric criteria without accounting for potential conformational adaptations of gp120–gp41, antibodies, or the membrane. The resulting accessibility frequencies should therefore be interpreted as an approximation rather than a definitive indicator of binding competence. Our results indicate that ectodomain epitopes, though heavily shielded by glycans, can still become transiently accessible in the closed prefusion state. The frequency provides a quantitative measurement of how vulnerable each epitope is to antibody binding. When estimating steric clashes between the membrane and antibodies, we adopted relaxed cutoffs to account for lipid fluidity, but the membrane due to fast lipid dynamics may in reality accommodate even greater steric overlap. Although the MPER can transiently extend out of the membrane when gp120–gp41 remains in the prefusion state, MPER-targeting antibodies encounter substantial steric hindrance from the bulky gp120 subunit, surrounding glycans, and the membrane throughout nearly all simulations. Therefore, these antibodies are unlikely to adopt the orientations required to bind their epitopes, which is consistent with experiment studies suggesting that MPER-targeting antibodies bind effectively only after the gp120–gp41 trimer undergoes major conformational rearrangements toward a fusion-intermediate or post-fusion state (Frey et al., 2008; Alam et al., 2009; Chen et al., 2014; Lee et al., 2016). Shehata et al. also built a model of the full-length gp120–gp41 trimer embedded in a lipid bilayer and performed all-atom simulations, in which a tilting motion of the ectodomain was observed. Based on the analysis of accessible surface area using different probe radii, they reported that antibody epitopes on the ectodomain are largely shielded by glycans, while the MPER epitope is mainly occluded by the membrane, with tilt angles above 30° required to achieve greater MPER exposure (Shehata et al., 2025).

Overall, the data presented here demonstrate that structural modeling integrated with all-atom MD simulations can be applied to complex biomolecular systems like the gp120–gp41 trimer embedded in a membrane bilayer, which enables detailed characterization of protein dynamics at the molecular level and facilitates the investigation of antigen-antibody interactions, thereby offering potential guidance for rational vaccine design.

## Materials and methods

### System building

#### Modeling of full-length gp120–gp41 trimer

The full-length gp120–gp41 trimeric model was generated by combining the crystal structure of the ectodomain with the NMR structure containing the MPER, TMD, and CT (Figure 1A, Figure 1—figure supplement 1A). Considering structure resolution and the number of missing residues, we selected the crystal structure of a soluble Env trimer exhibiting a closed-form, native-like prefusion conformation (PDB ID: 6B0N) (Sarkar et al., 2018). This structure contains the entire ectodomain, including the V1-V5 loops, fusion peptide (FP), heptad repeat 1 (HR1), and heptad repeat 2 (HR2) (Figure 1B). In contrast to the ectodomain well represented in the PDB with numerous crystal and cryo-EM structures, structural data for the MPER, TMD and CT are scarce. For these regions, we chose the NMR structure (PDB ID: 7LOI) (Piai et al., 2021) containing the MPER, TMD, and CT. Both 6B0N and 7LOI are trimeric structures and the C-terminus of 6B0N includes five residues (L660–D664) overlapping with the N-terminus of 7LOI. Measurement of the Cα–Cα distances for D664 in the three protomers revealed that the inter-protomer spacing in 7LOI (16 Å) was significantly shorter than in 6B0N (39 Å) (Supplementary file 1–Supplementary Figure 1A). To reconcile this difference, we performed short MD simulations on 6B0N with residues E32–Q640 fixed and a distance restraint applied to D664 to reduce the Cα–Cα distance to 33 Å. Similarly, for 7LOI, residues I675–L856 were fixed, and a distance restraint was applied to D664 to increase the Cα–Cα distance to 33 Å (Supplementary file 1–Supplementary Figure 1B). These two adjusted structures were then combined by aligning the overlapping residues (Supplementary file 1–Supplementary Figure 1C). The missing loops in 6B0N (T63, D149, E185A–N185I, S401–G409) and the unresolved region between the TMD and CT in 7LOI (F717–G738) were grafted from a modeled structure generated by I-TASSER (Yang et al., 2015).

6B0N is a native flexibly linked (NFL) trimer in which the furin cleavage site ^508^REKR^511^ is substituted with a 10-residue linker ^508^GG<u>GGSGGGG</u>S^511^, but the underlined residues are unresolved in the structure. To construct the cleaved gp120–gp41 model, we removed the initial “GG” and final “S” from 6B0N. For the uncleaved gp160 model, we closed the cleaved loop by adding ^508^REKR^511^ back to the structure. If we simply connect G507 and A512 without adjusting the conformation of flanking loops on both sides of the cleavage site, two neighboring protomers become entangled, yielding a knot-like fold (Supplementary file 1–Supplementary Figure 2A, B). In 6B0N, the helical HR2 region lies at the C-terminus of each protomer, allowing it to thread through a loop formed by the neighboring protomer (Supplementary file 1–Supplementary Figure 2C). However, in the context of the full-length gp120–gp41, it is implausible that the three protomers are entangled as it does not conform to a realistic folding pathway. Therefore, we performed short MD simulations with restraints to adjust the conformations of the flanking loops and the HR2 helix before closing the loop with REKR fragment (Supplementary file 1–Supplementary Figure 2D, E).

Finally, two mutations (S764C and S837C) were introduced in the CT to restore the palmitoylation sites, and lipid tails oriented towards the hydrophobic core of the bilayer were then attached to the palmitoylation sites using the PDB Manipulation module in CHARMM-GUI (Jo et al., 2008; Jo et al., 2014; Park et al., 2023) (Figure 1D). In addition to the full-length model, we built the CT-truncated model by removing modeled residues F717–G738 that are originally unresolved in the PDB structure 7LOI, and residues E739–L856 that form the large CT plate.

#### Glycosylation

N-linked glycans were modeled using *Glycan Reader & Modeler* (Jo et al., 2011; Park et al., 2017; Park et al., 2019) in CHARMM-GUI. The reported mass spectrometry (MS) experiments have revealed the probability of different types of N-linked glycans at each glycosylation site (Behrens et al., 2016; Cao et al., 2017). For each site, we selected the one with the highest probability. As the MS data provide only the glycan type (high-mannose, hybrid, and complex) and composition (number of each monosaccharide unit), a representative isomer was chosen whenever multiple isomers corresponded to the same composition. For example, HexNAc(2)Hex(9), i.e., the Man_9_ glycan, has a single isomer, whereas HexNAc(2)Hex(8), i.e., the Man_8_ glycan, has multiple isomers depending on which non-reducing terminal mannose is trimmed from Man_9_. For complex N-linked glycans, a single composition can correspond to multiple isomers differing in the number of branches on the α1-3 and α1-6 arms and in the placement of non-reducing terminal neuraminic acid (Neu5Ac). In the absence of linkage-specific information, one isomer was arbitrarily selected from available possibilities. The selected glycan sequences for the 27 glycosylation sites in each promoter are summarized in Supplementary file 1–Supplementary Table 6. The selected glycan sequences were represented in the Glycan Reader Sequence format (Jo et al., 2011; Park et al., 2017) and added to the corresponding glycosylation sites using the *Glycan Reader & Modeler* graphical interface.

#### Membrane

The full-length and CT-truncated models were embedded into an asymmetric lipid bilayer with the lipid composition corresponding to a mammalian plasma membrane (van Meer et al., 2008; Sampaio et al., 2011; Ingolfsson et al., 2014; Pogozheva et al., 2022), which consists of phosphatidylcholine (PC), phosphatidylethanolamine (PE), phosphatidylinositol (PI), phosphatidylserine (PS), phosphatidic acid (PA), sphingomyelin (SM), cholesterol (CHOL), and glucosylceramide (GlcCer). In the CT-truncated system, the exoplasmic and cytoplasmic leaflets contain similar numbers of lipids, despite differences in composition. In the full-length system, the cytoplasmic leaflet contains approximately 100 fewer lipids than the exoplasmic leaflet due to the space occupied by the CT. The lipid composition is summarized in Supplementary file 1–Supplementary Table 7. Membrane systems were constructed using CHARMM-GUI Membrane Builder, which provides a user-friendly graphical interface for selecting lipid types and defining their numbers in each leaflet (Jo et al., 2007; Jo et al., 2009; Wu et al., 2014; Lee et al., 2016; Lee et al., 2019). Although it is recognized that the charged residues R683, R707, and R709, flanking the N- and C-termini of the TMD, are anchored within the lipid headgroups, there remains flexibility to shift the TMD slightly up or down within the membrane. Assuming the lipid bilayer is aligned parallel to the xy-plane, the protein was positioned at two distinct depths along the z-axis, differing by approximately 4 Å, referred to as the “high” and “low” TMD configurations. To examine whether this subtle difference in the initial configurations leads to distinct conformational changes during simulations, we initiated independent simulations from each configuration. The glycoprotein-membrane system was solvated in a box of approximately 210 × 210 × 260 Å^3^ using the TIP3P water model (Jorgensen et al., 1983), and KCl was added at a concentration of 0.15 M to neutralize the system. The final simulation input files were automatically generated by CHARMM-GUI.

#### Simulation details

The CHARMM36(m) force field (Guvench et al., 2009; Klauda et al., 2010; Raman et al., 2010; Guvench et al., 2011; Huang et al., 2017) was used for proteins, carbohydrates, and lipids. The total number of atoms is approximately 1,100,000 (including ∼300,000 water molecules and ∼1,700 ions), with small variations between systems. van der Waals interactions were smoothly switched off over 10–12 Å using a force-based switching function (Steinbach et al., 2004), and long-range electrostatic interactions were calculated using the particle-mesh Ewald method (Essmann et al., 1995) with a mesh size of ∼1 Å. All simulations were performed using GROMACS (Van Der Spoel et al., 2005). Bond lengths and angles involving hydrogens were constrained using the LINCS algorithm (Hess et al., 1997). The system was equilibrated in the canonical (NVT) ensemble at 310.15 K for 2 × 1.25 ns with a 1-fs time step using the Berendsen thermostat (Berendsen et al., 1984) (coupling constant *τ*_t_ = 1 ps), and then switched to the isothermal–isobaric (NPT) ensemble for 1.25 ns with a 1-fs time step, followed by 3 × 5 ns with a 2-fs time step using the Berendsen thermostat (*τ*_t_ = 1 ps) and barostat (reference pressure = 1 bar, coupling constant *τ*_p_ = 5 ps, compressibility = 4.5×10^−5^ bar^−1^). Positional and dihedral restraints were applied to proteins, glycans, and lipids, with force constants progressively reduced over successive intervals (Supplementary file 1–Supplementary Table 8). In the production runs, temperature was maintained using the Nosé–Hoover thermostat (Hoover, 1985; Nosé, 2006) and the pressure coupling was applied using the semi-isotropic Parrinello–Rahman barostat (Parrinello et al., 1981; Nosé et al., 2006). A 4-fs time step was used with the hydrogen mass repartitioning technique (Hopkins et al., 2015) and all restraint potentials were removed. The Python library MDTraj (McGibbon et al., 2015) was used to analyze the simulation trajectories.

#### Assessment of antibody epitope accessibility

We selected six antibodies based on epitope locations and the availability of PDB structures containing both antibody and epitope: PGT128 targeting the V3 loop and the N332 glycan (PDB ID: 5JSA); PG9 targeting the V1/V2 loop (PDB ID: 3U2S); VRC01 targeting the CD4 binding site (PDB ID: 4LST); 35O22 targeting the gp120–gp41 interface (PDB ID: 4TVP); 10E8 (PDB ID: 6VPX) and 4E10 (PDB ID: 1TZG) targeting the MPER. For each antibody–epitope complex, TM-align (Zhang et al., 2005) was used to identify the optimal alignment between the epitope from the PDB structure and each protomer of the trimeric protein. The resulting rotation matrix was applied to position the antibody relative to each protomer. Steric clashes were quantified by counting any heavy atoms of the protein, glycans, or lipids within 2 Å of a heavy atom of the antibody. An epitope was classified as occluded by the protein and glycans if more than 10 heavy-atom clashes occurred with these components. To account for the inherent flexibility and fluidity of the membrane, a more permissive threshold was adopted for lipids: an epitope was classified as occluded by the membrane only if more than 20 lipid heavy-atom clashes were detected. To calculate the frequency of epitope accessibility, snapshots were extracted from simulation trajectories, and then structural alignment and assessment of steric clashes were performed on each snapshot.

## Data availability

The modeled structures, force field parameters, and GROMACS input files are available on Zenodo [https://doi.org/10.5281/zenodo.17259909]. The code used in this article can be found at https://doi.org/10.5281/zenodo.17259951. The simulation trajectories can be found at https://doi.org/10.5281/zenodo.18853902, https://doi.org/10.5281/zenodo.18854615, and https://doi.org/10.5281/zenodo.18854639.

## Article and author information

### Author details

**Yiwei Cao**

Department of Biological Sciences, Lehigh University

**Contribution:** Conceptualization, Formal analysis, Investigation, Methodology, Writing – original draft, Writing – review and editing

**Competing interests:** No competing interests declared

### Wonpil Im

Department of Biological Sciences, Lehigh University

**Contribution:** Conceptualization, Funding acquisition, Methodology, Supervision, Writing – review and editing

**Competing interests:** No competing interests declared

---

**Figure 2—figure supplement 1.**
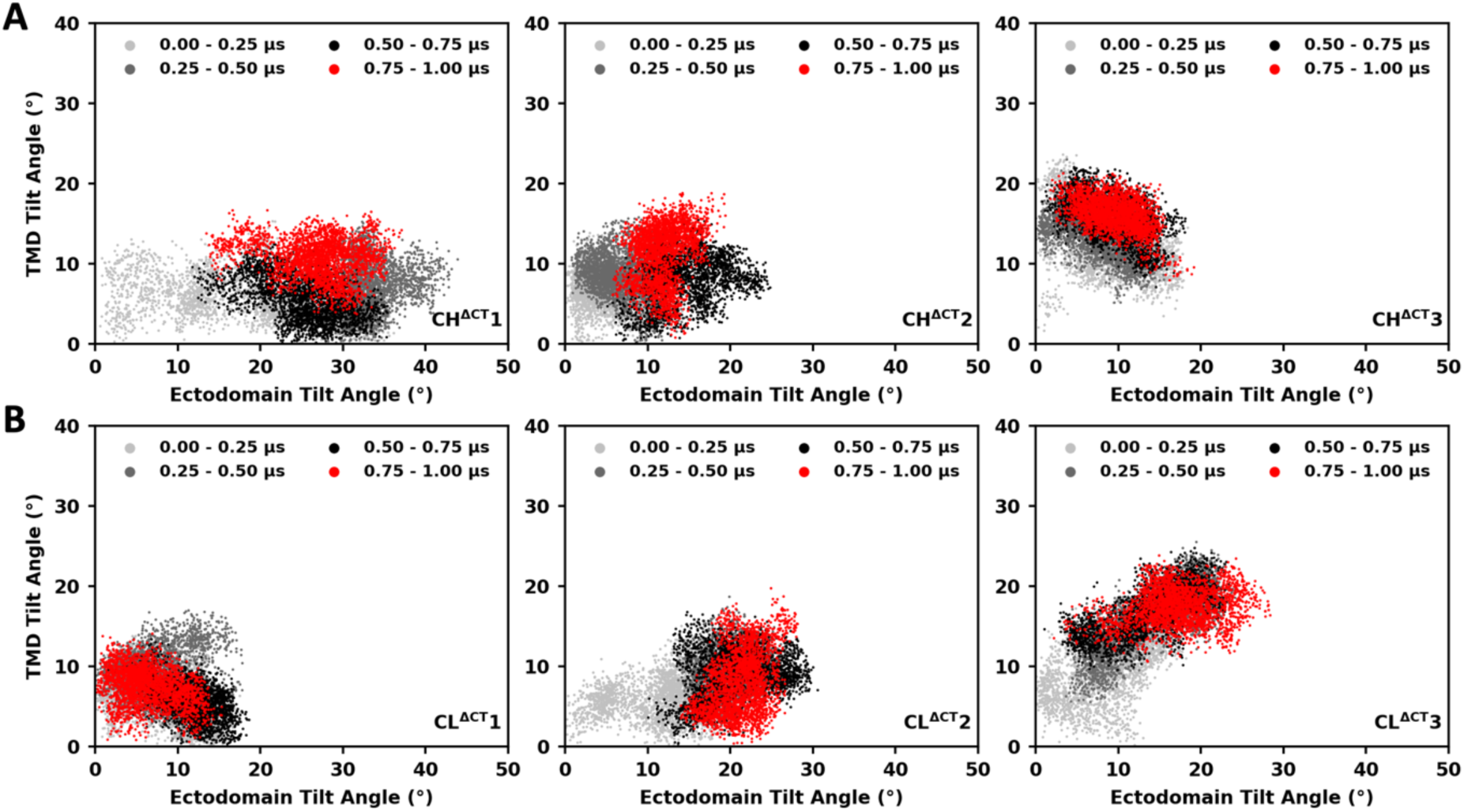
Ectodomain tilt versus TMD tilt, grouped by time intervals (cleaved CT-truncated systems). (**A**) Three trajectories starting from the “high” TMD configuration. (**B**) Three trajectories starting from the “low” TMD configuration. The 1-μs trajectory was divided into four intervals, with values from each interval shown in light gray, dark gray, black, and red, respectively.

**Figure 2—figure supplement 2.**
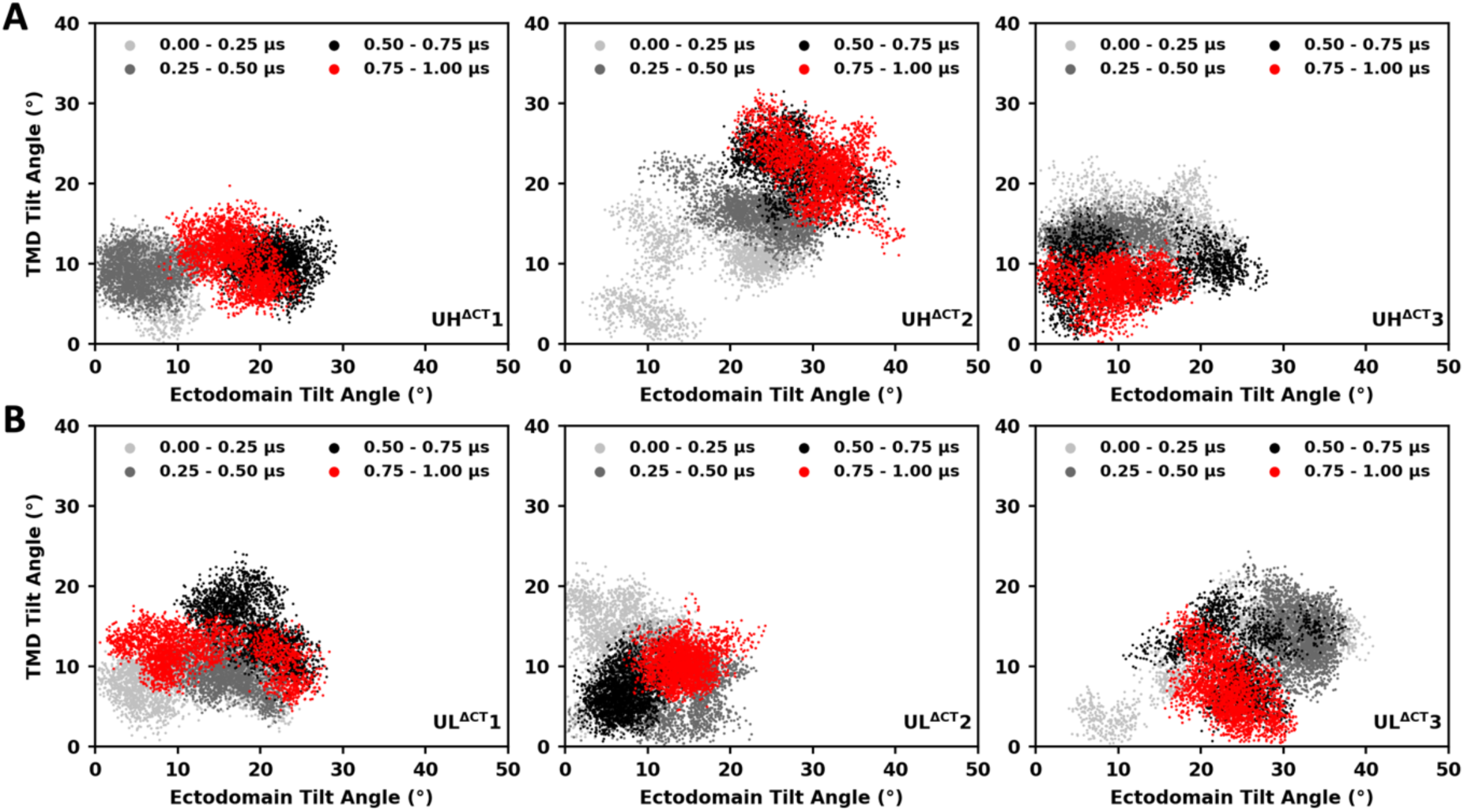
Ectodomain tilt versus TMD tilt, grouped by time intervals (uncleaved CT-truncated systems). Labeling and color coding are the same as in Figure 2—figure supplement 1.

**Figure 2—figure supplement 3.**
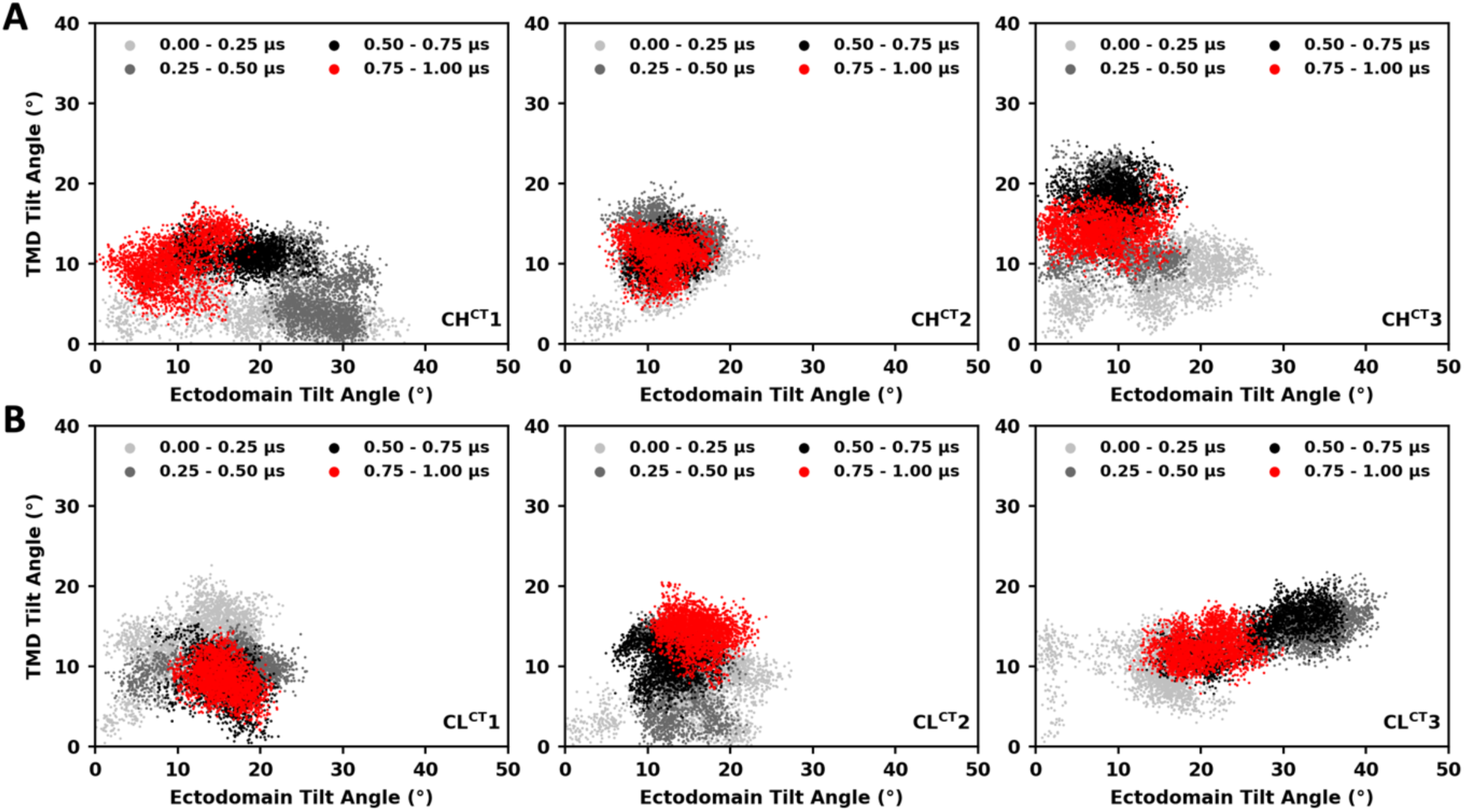
Ectodomain tilt versus TMD tilt, grouped by time intervals (cleaved full-length systems). Labeling and color coding are the same as in Figure 2—figure supplement 1.

**Figure 2—figure supplement 4.**
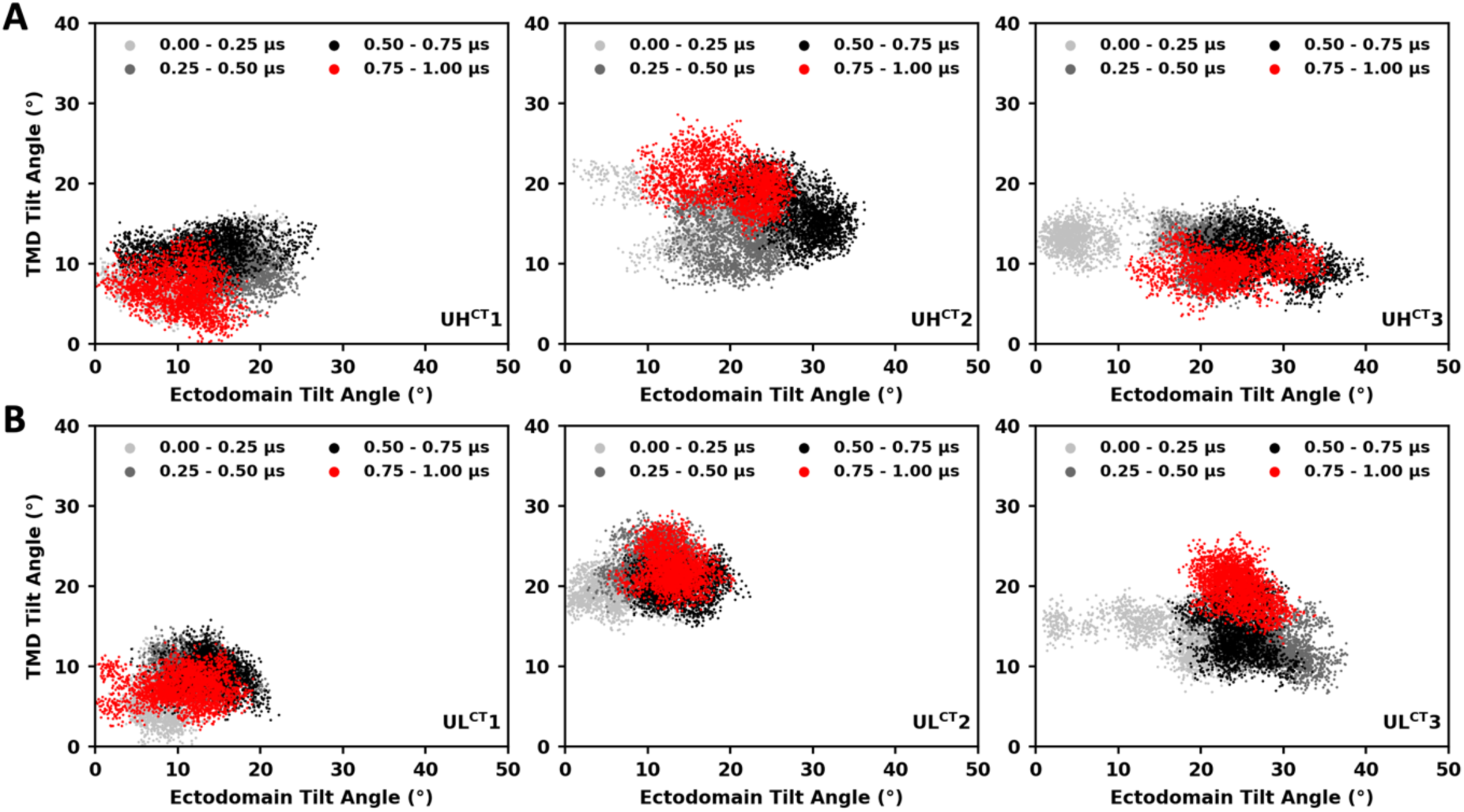
Ectodomain tilt versus TMD tilt, grouped by time intervals (uncleaved full-length systems). Labeling and color coding are the same as in Figure 2—figure supplement 1.

**Figure 2—figure supplement 5.**
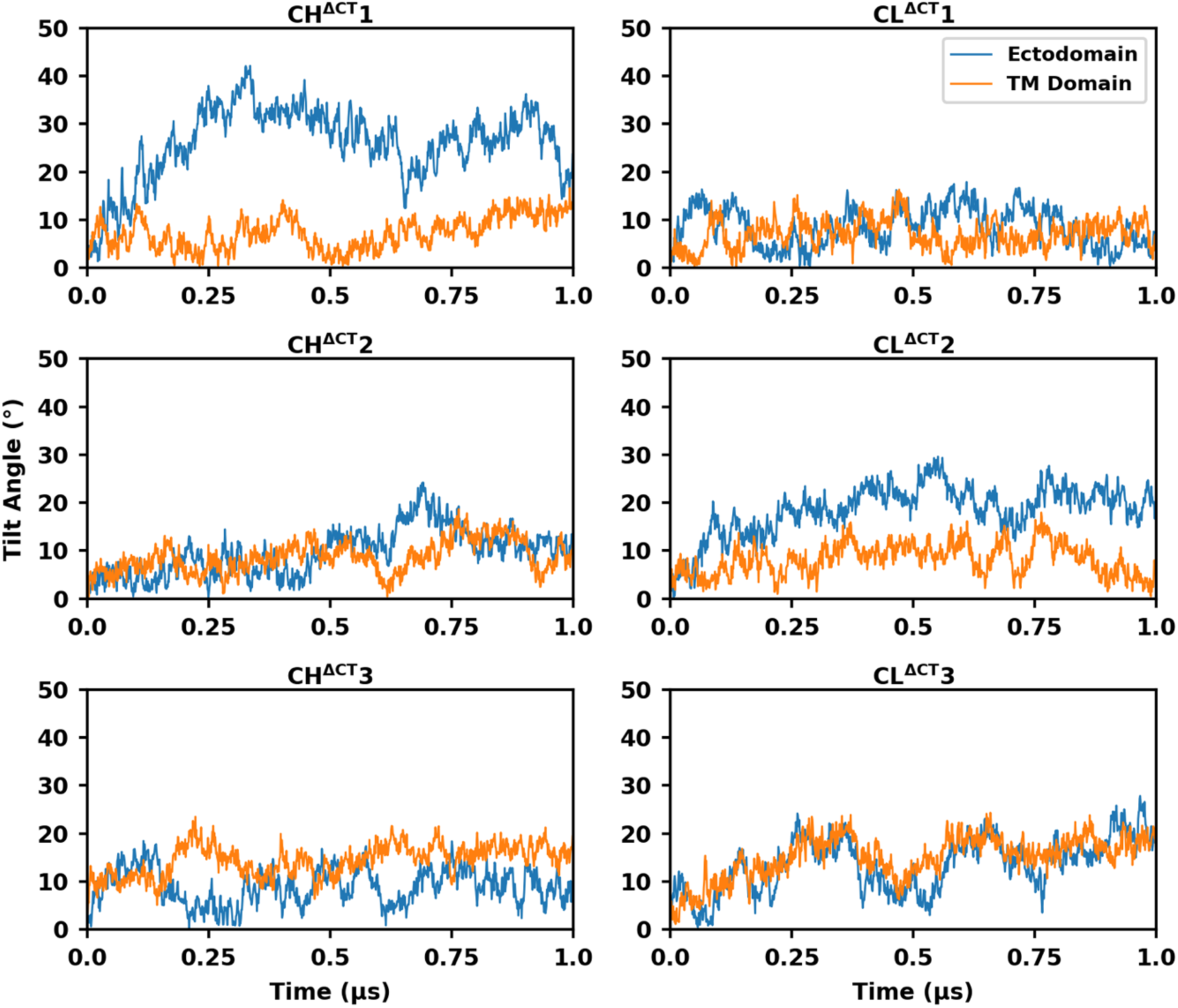
Temporal evolution of Ectodomain and TMD tilt angles (cleaved CT-truncated systems).

**Figure 2—figure supplement 6.**
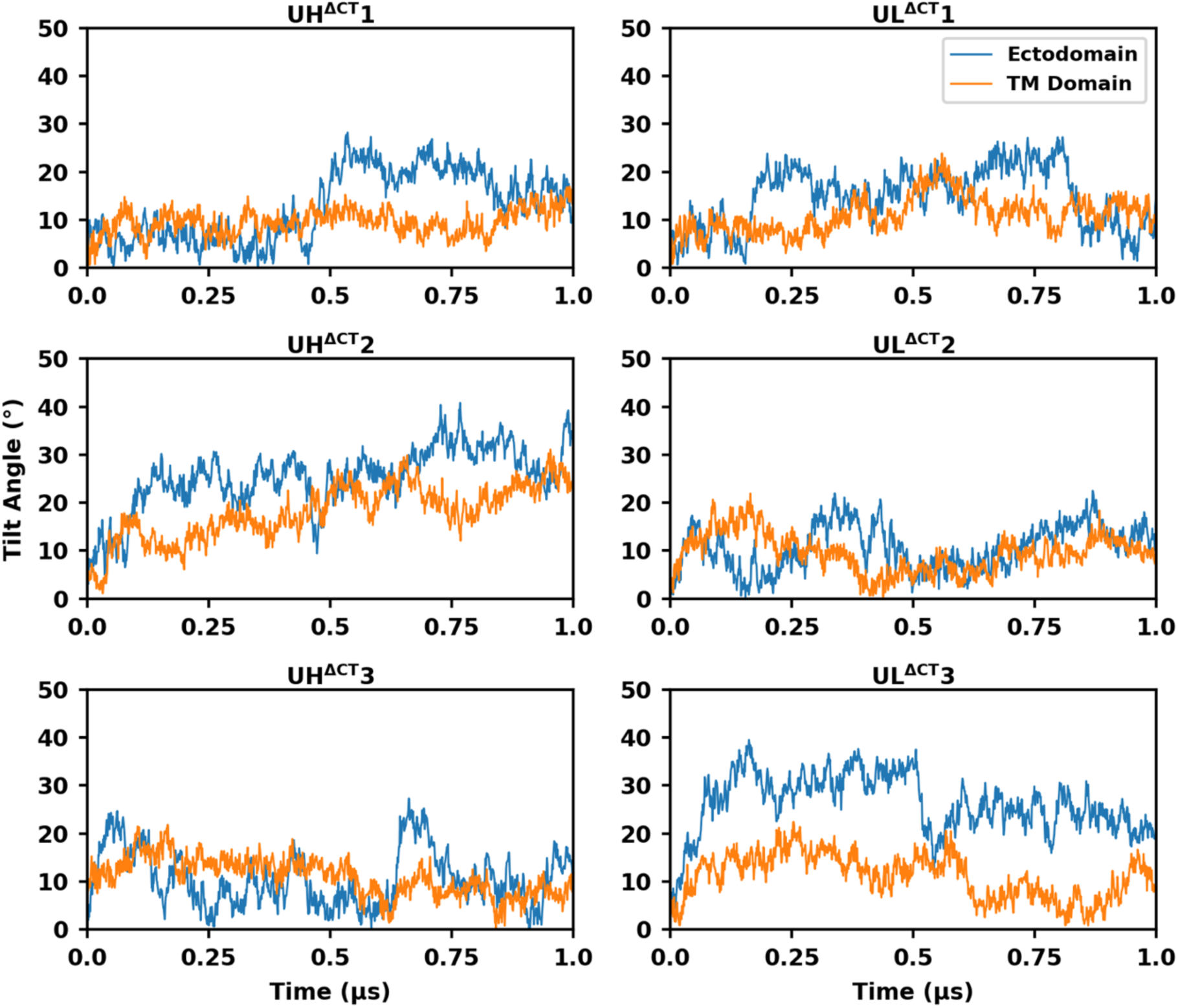
Temporal evolution of Ectodomain and TMD tilt angles (uncleaved CT-truncated systems).

**Figure 2—figure supplement 7.**
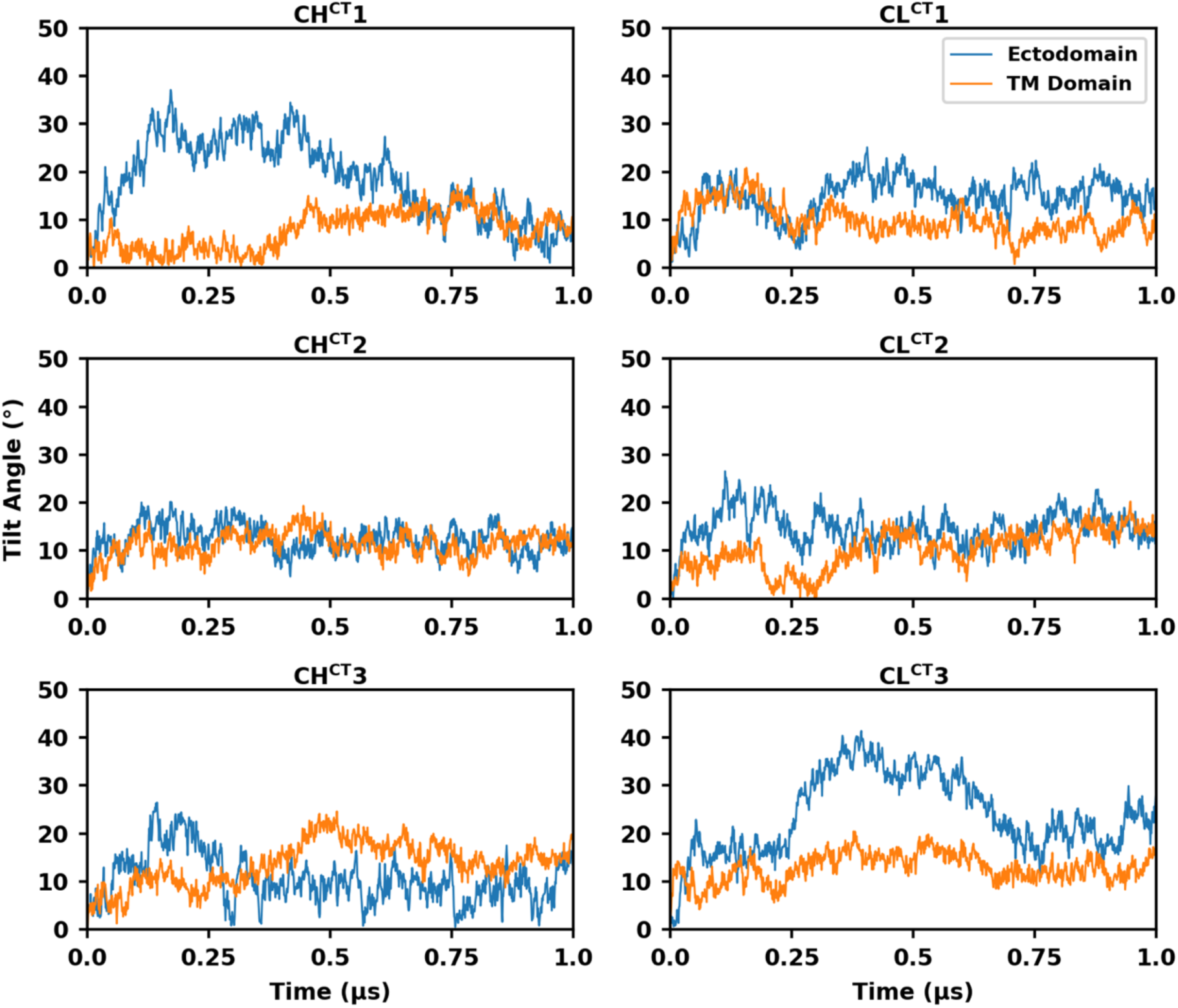
Temporal evolution of Ectodomain and TMD tilt angles (cleaved full-length systems).

**Figure 2—figure supplement 8.**
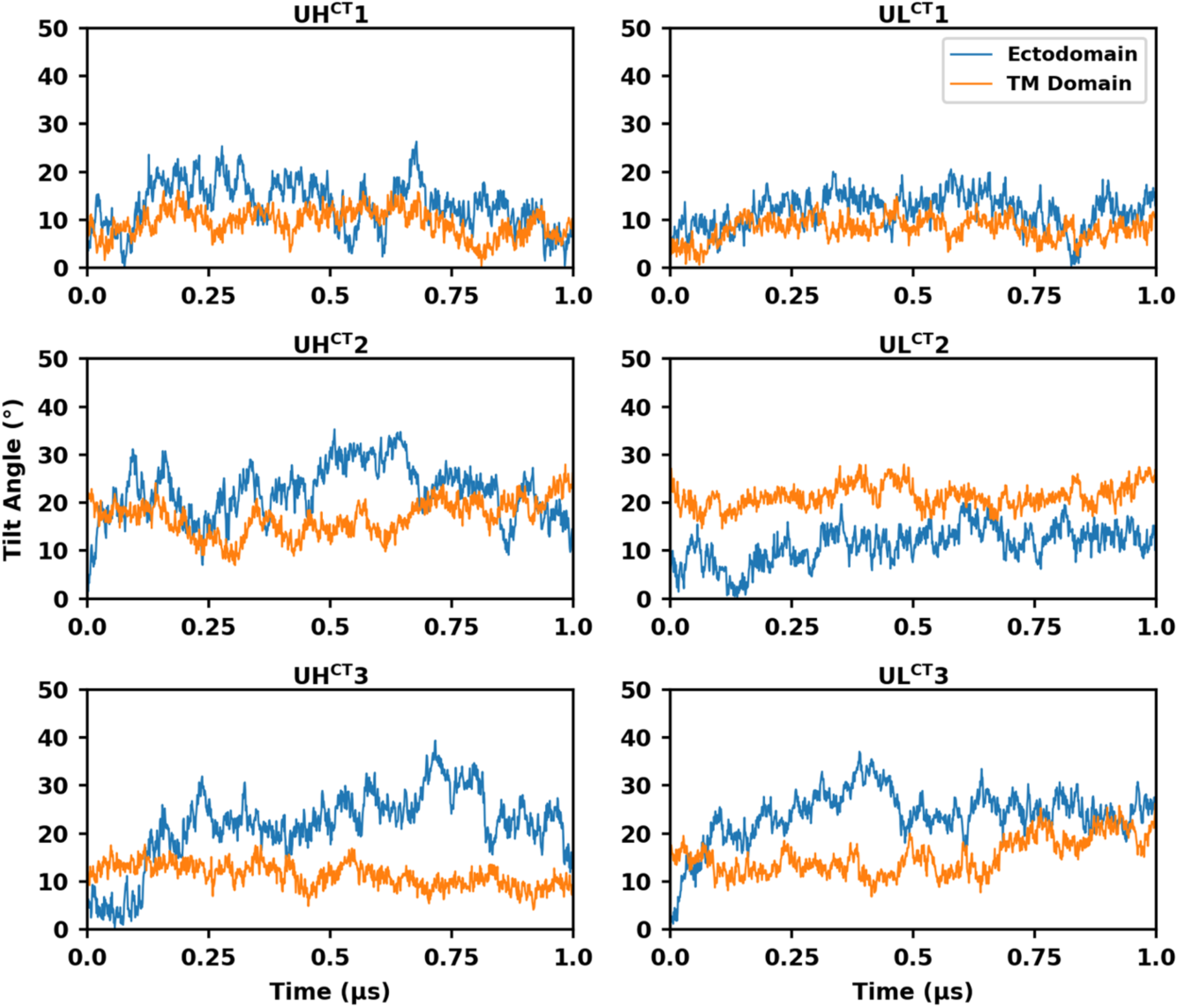
Temporal evolution of Ectodomain and TMD tilt angles (uncleaved full-length systems).

**Figure 2—figure supplement 9.**
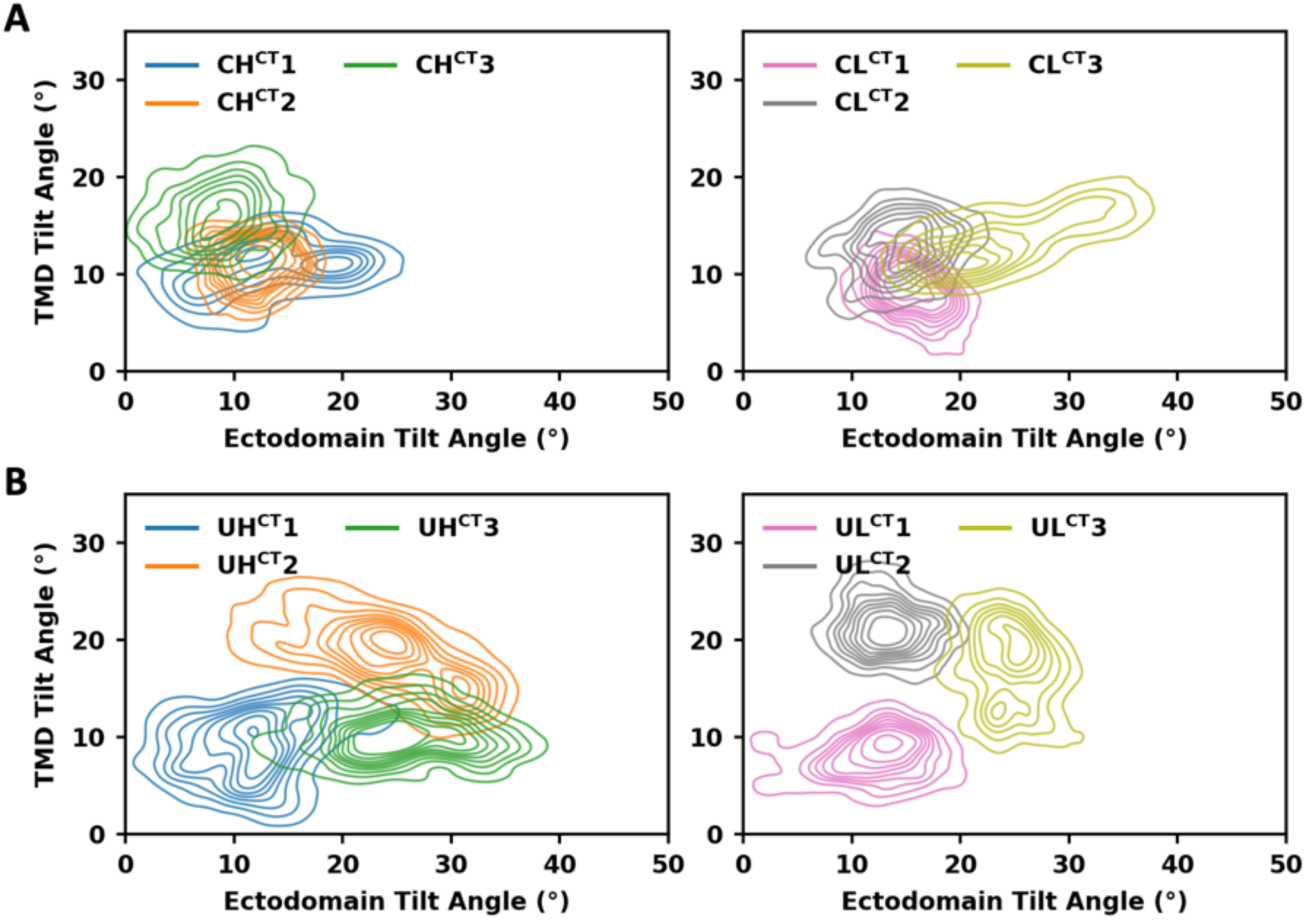
Probability densities of ectodomain and TMD tilt angles (full-length systems).

**Figure 2—figure supplement 10.**
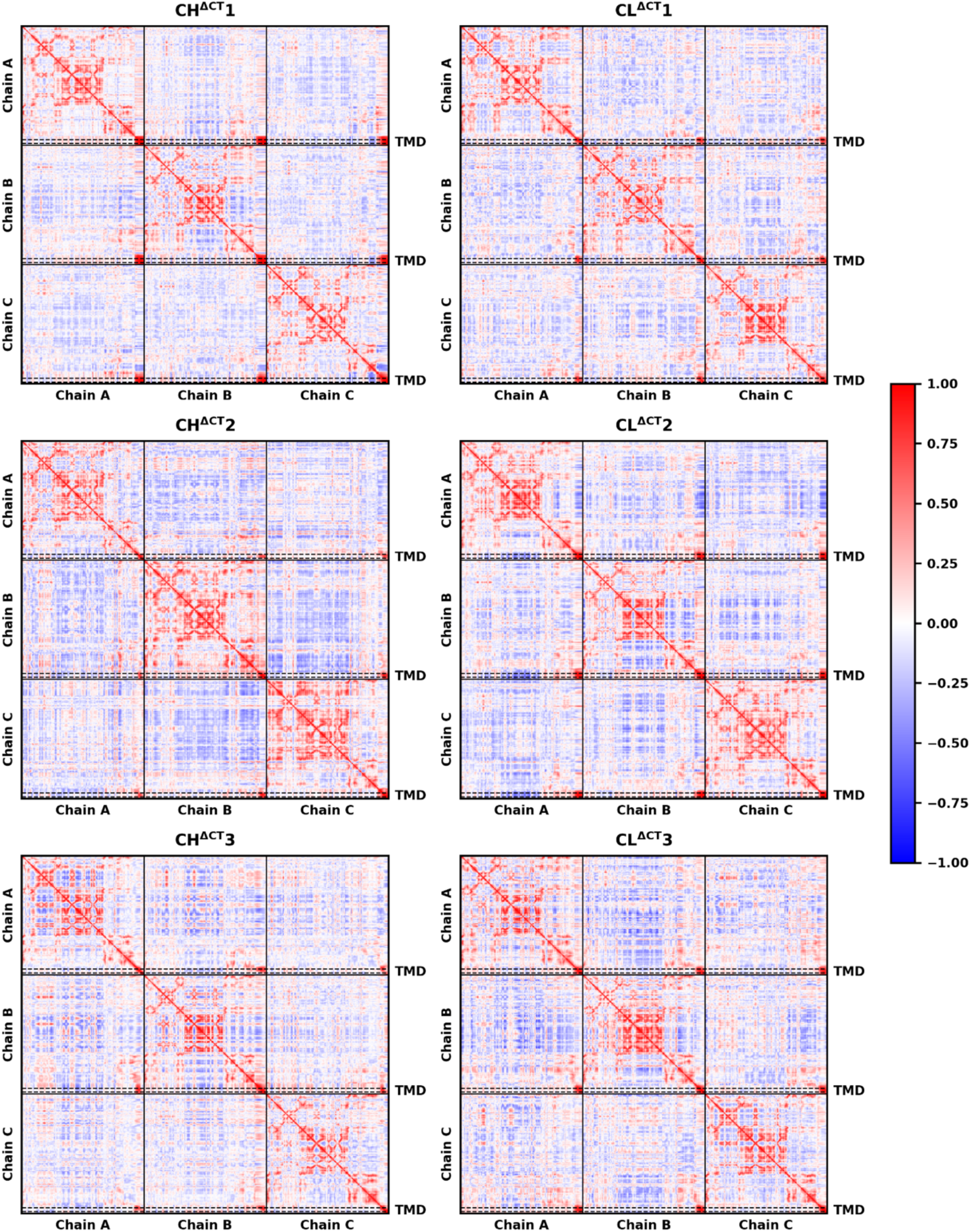
Dynamic cross-correlation matrix of Cα atoms (cleaved CT-truncated systems). The x-axis (left to right) and the y-axis (top to bottom) correspond to residue indices ranging from 31 to 716 for each of the three protomers, where residues 31-664 compose the ectodomain. The TMD (residues 684-705) is indicated by dashed boxes. Corelated and anti-correlated motions are color-coded from red to blue, respectively.

**Figure 2—figure supplement 11.**
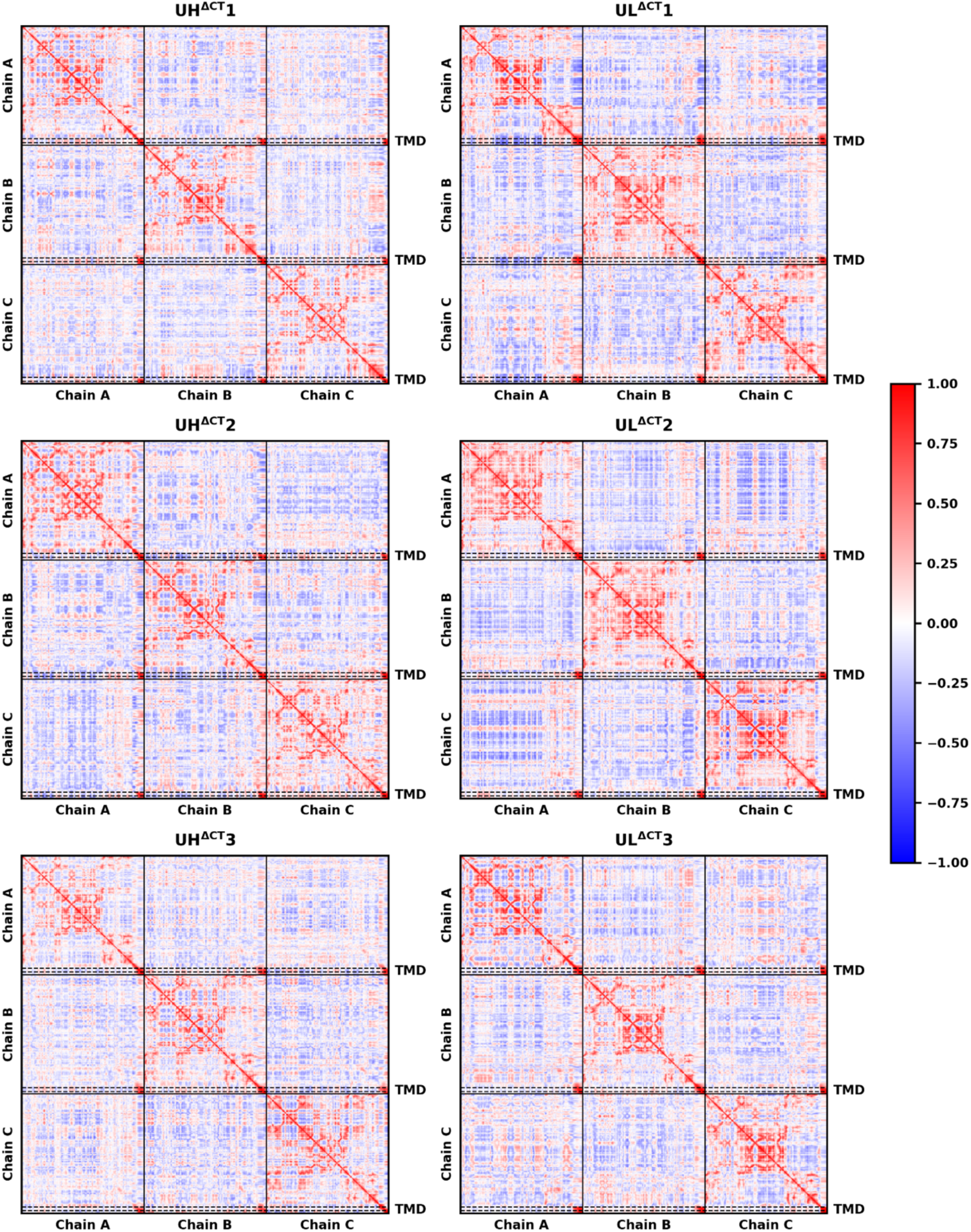
Dynamic cross-correlation matrix of Cα atoms (uncleaved CT-truncated systems). The x-axis (left to right) and the y-axis (top to bottom) correspond to residue indices ranging from 31 to 716 for each of the three protomers, where residues 31-664 compose the ectodomain. The TMD (residues 684-705) is indicated by dashed boxes. Corelated and anti-correlated motions are color-coded from red to blue, respectively.

**Figure 2—figure supplement 12.**
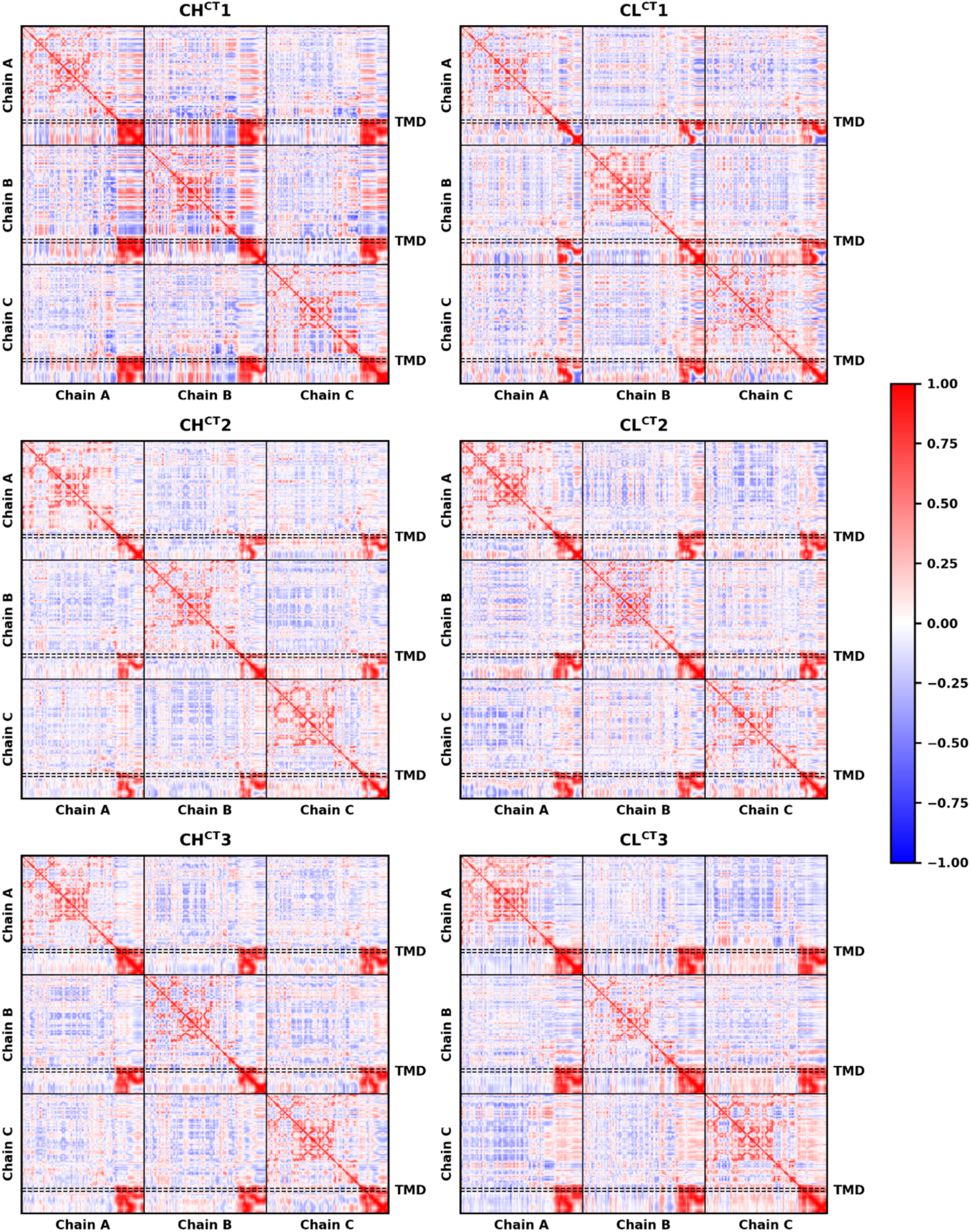
Dynamic cross-correlation matrix of Cα atoms (cleaved full-length systems). The x-axis (left to right) and the y-axis (top to bottom) correspond to residue indices ranging from 31 to 716 for each of the three protomers. Residues 31-856 compose the ectodomain, and residues 706-856 compose the CT. The TMD (residues 684-705) is indicated by dashed boxes. Corelated and anti-correlated motions are color-coded from red to blue, respectively.

**Figure 2—figure supplement 13.**
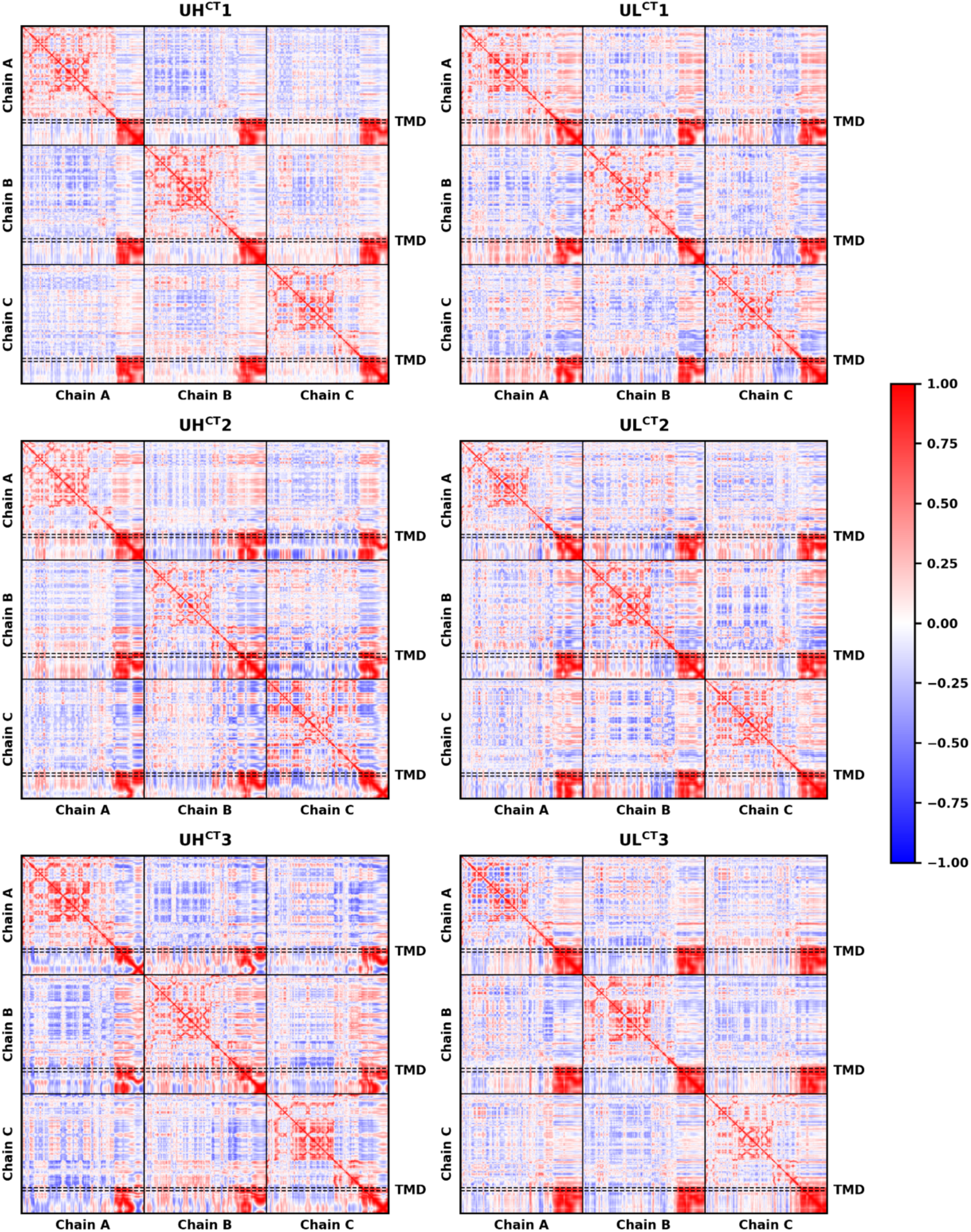
Dynamic cross-correlation matrix of Cα atoms (uncleaved full-length systems). The x-axis (left to right) and the y-axis (top to bottom) correspond to residue indices ranging from 31 to 716 for each of the three protomers. Residues 31-856 compose the ectodomain, and residues 706-856 compose the CT. The TMD (residues 684-705) is indicated by dashed boxes. Corelated and anti-correlated motions are color-coded from red to blue, respectively.

**Figure 3—figure supplement 1.**
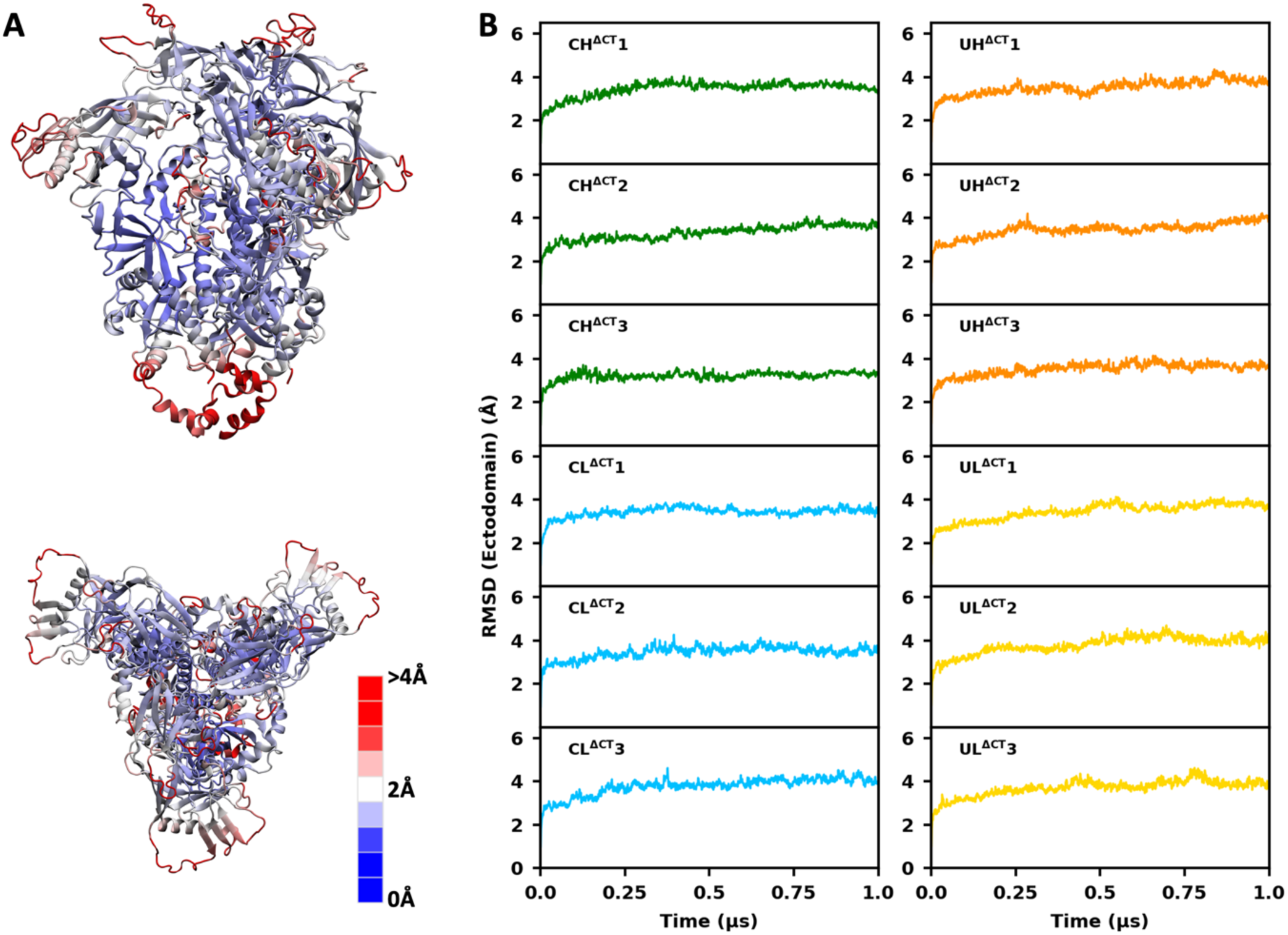
RMSF and RMSD of the ectodomain. (**A**) Top and side views of the ectodomain and MPER in the uncleaved system, with RMSF indicated by color. (**B**) RMSD relative to the initial model as a function of time, calculated from the trajectories of CT-truncated systems.

**Figure 4—figure supplement 1.**
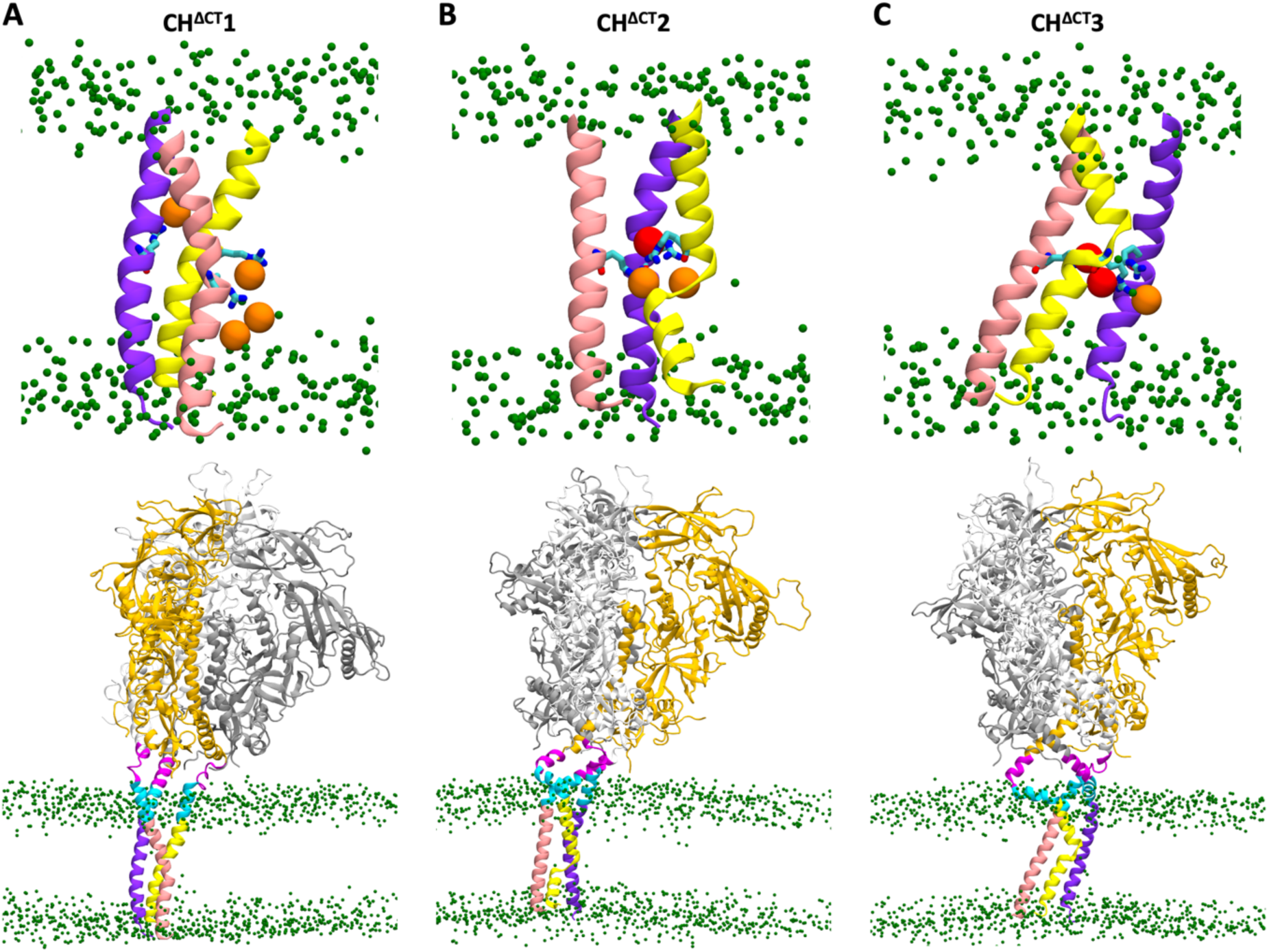
Local conformations of the TMD and global conformations of protein and membrane (CH^ΔCT^ systems). Three protomers of the TMD are shown in light yellow, purple, and pink; three protomers of the ectodomain in dark yellow, gray, and white; MPER-N and MPER-C in magenta and cyan, respectively; and lipid headgroups in green. Lipid headgroups interacting with R696 are highlighted in orange, and the ions interacting with R696 in red.

**Figure 4—figure supplement 2.**
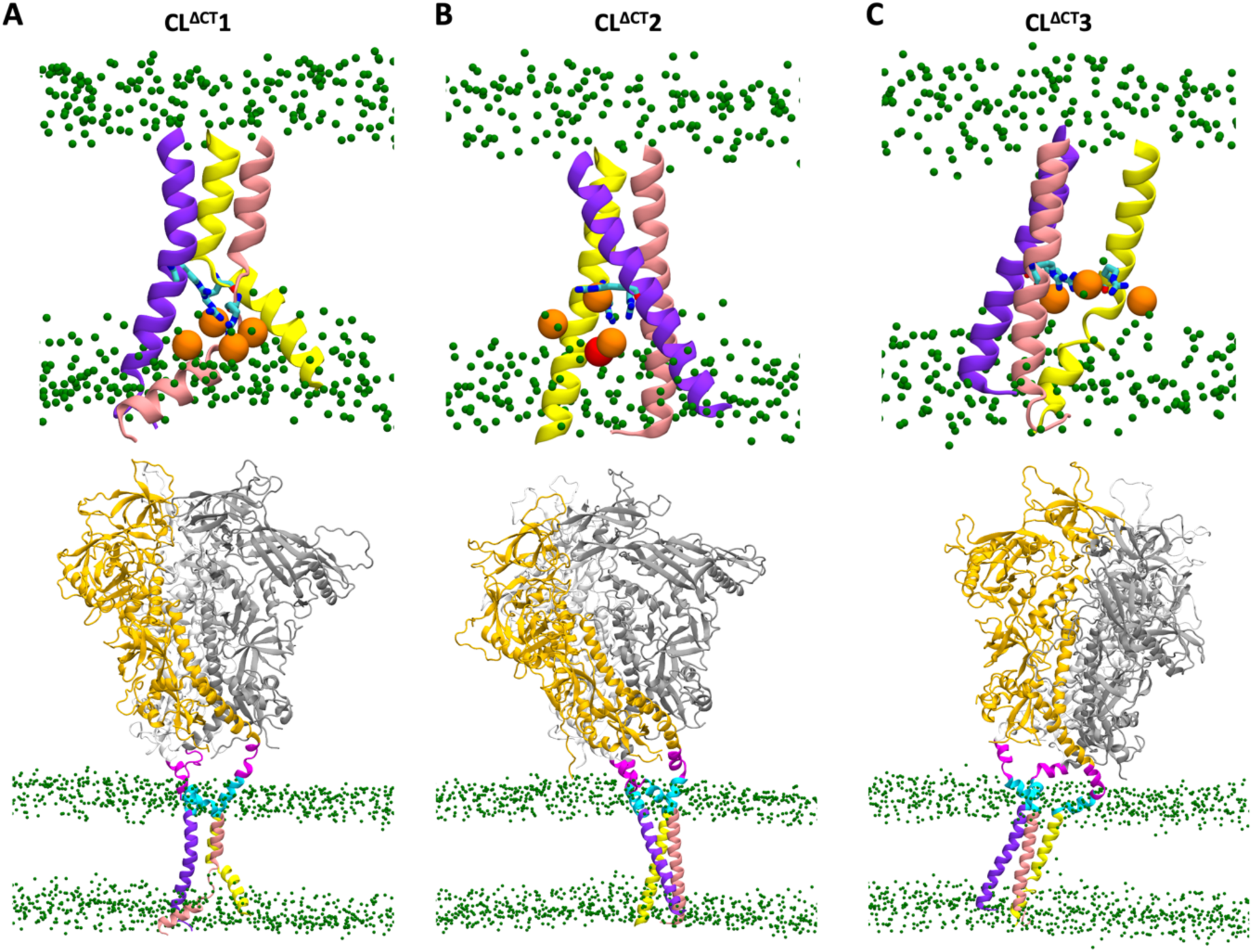
Local conformations of the TMD and global conformations of protein and membrane (CL^ΔCT^ systems). Labeling and color coding are the same as in Figure 4—figure supplement 1.

**Figure 4—figure supplement 3.**
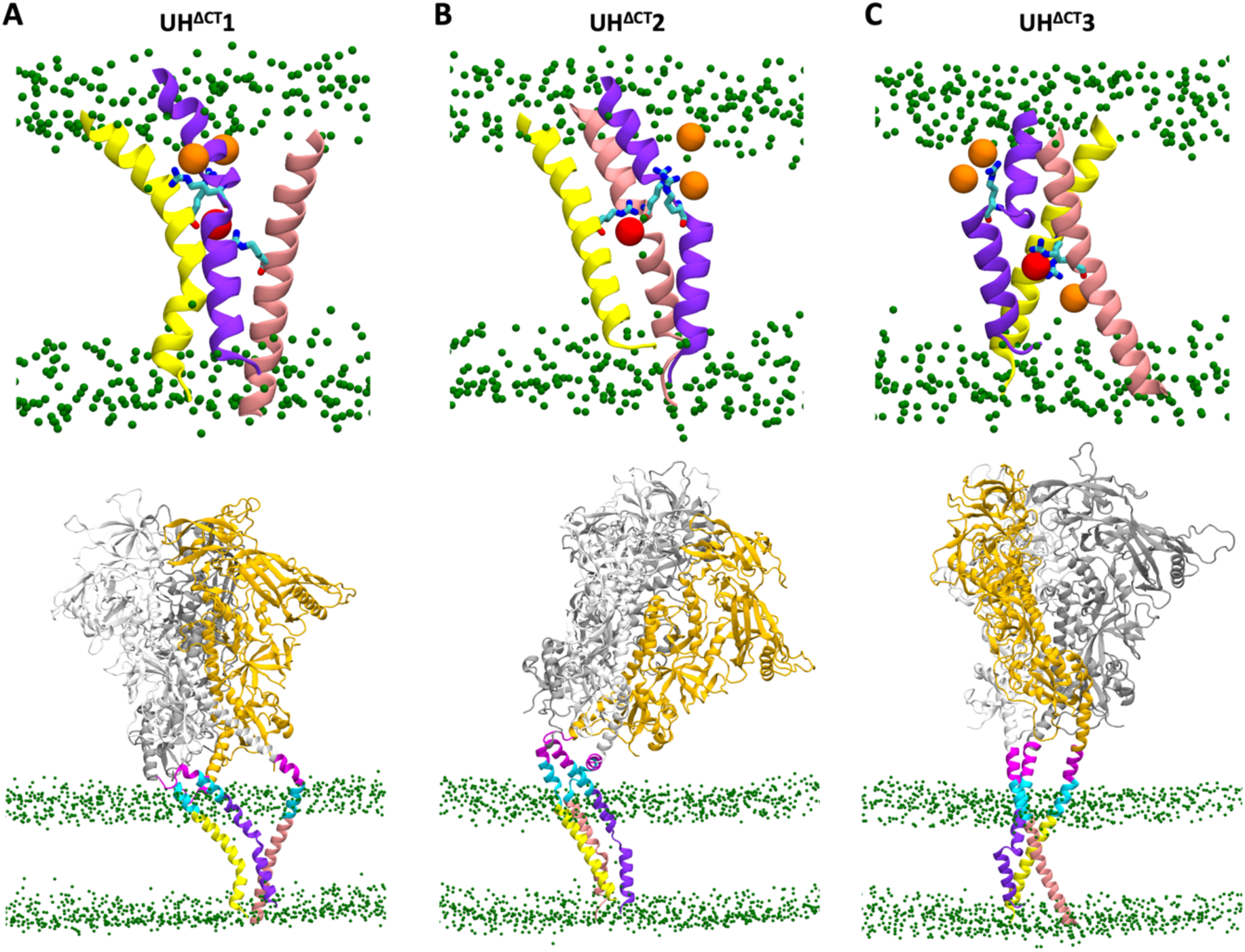
Local conformations of the TMD and global conformations of protein and membrane (UH^ΔCT^ systems). Labeling and color coding are the same as in Figure 4—figure supplement 1.

**Figure 4—figure supplement 4.**
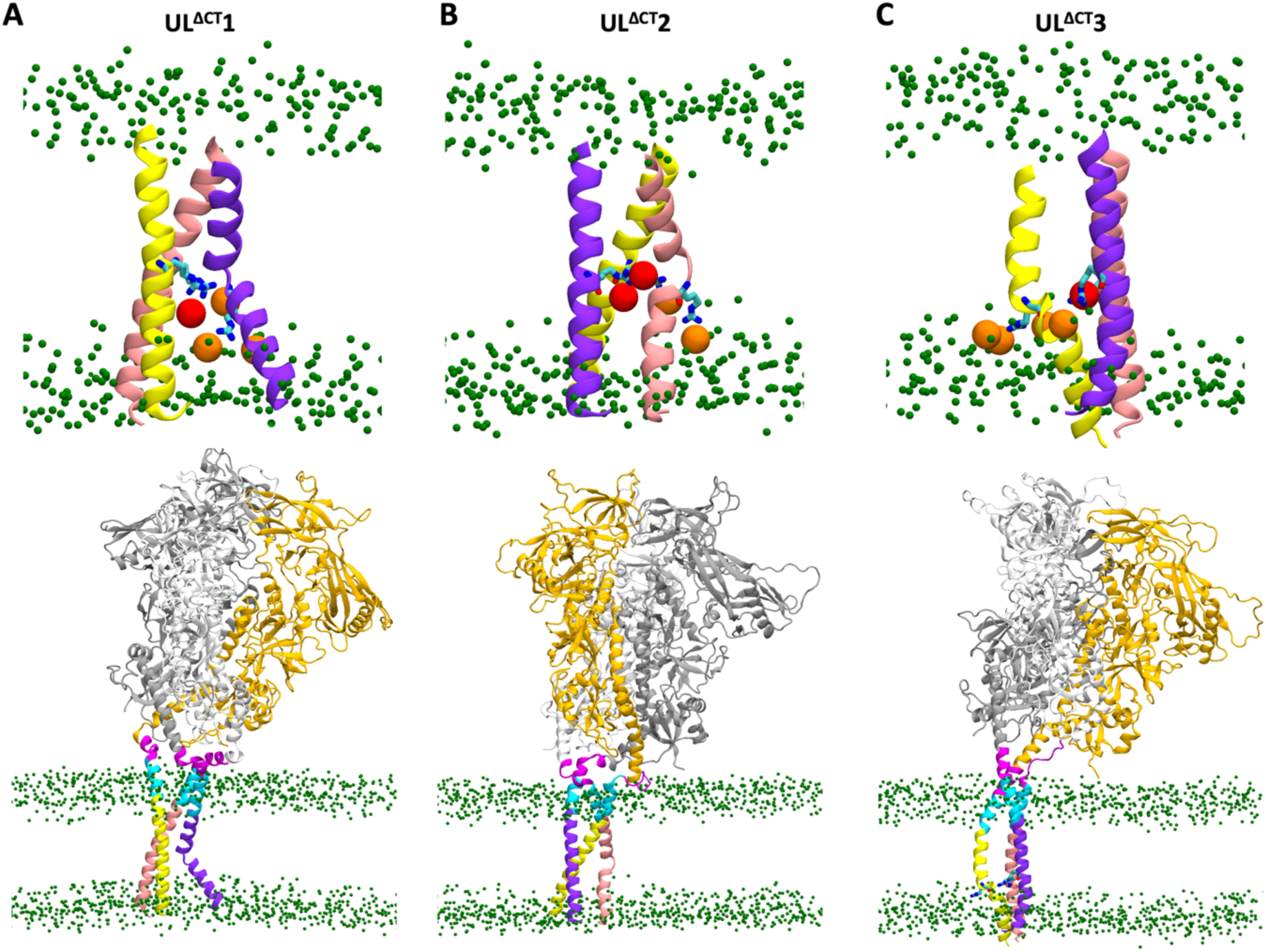
Local conformations of the TMD and global conformations of protein and membrane (UL^ΔCT^ systems). Labeling and color coding are the same as in Figure 4—figure supplement 1.

**Figure 4—figure supplement 5.**
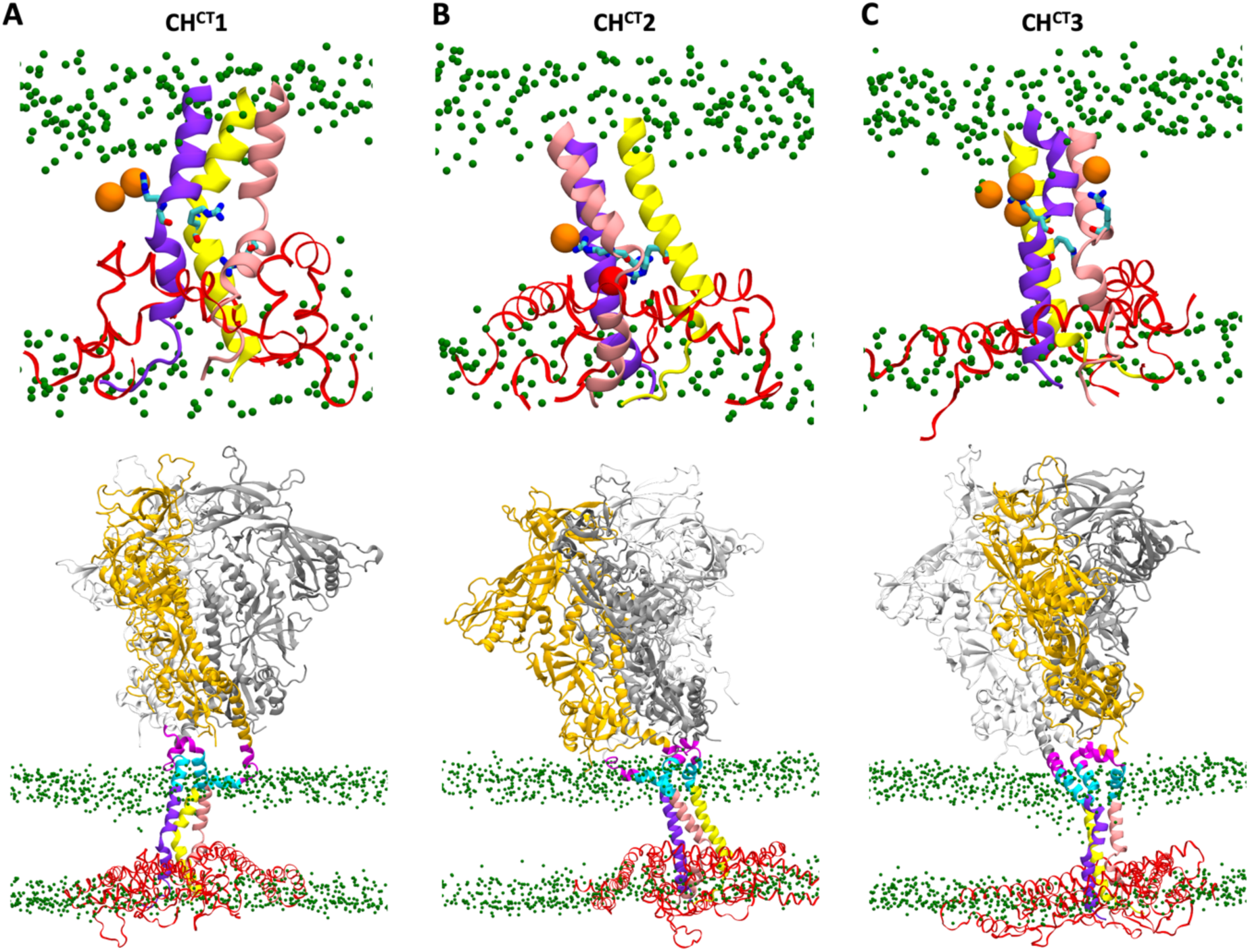
Local conformations of the TMD and global conformations of protein and membrane (CH^CT^ systems). Labeling and color coding are the same as in Figure 4—figure supplement 1, with the CT additionally shown in red.

**Figure 4—figure supplement 6.**
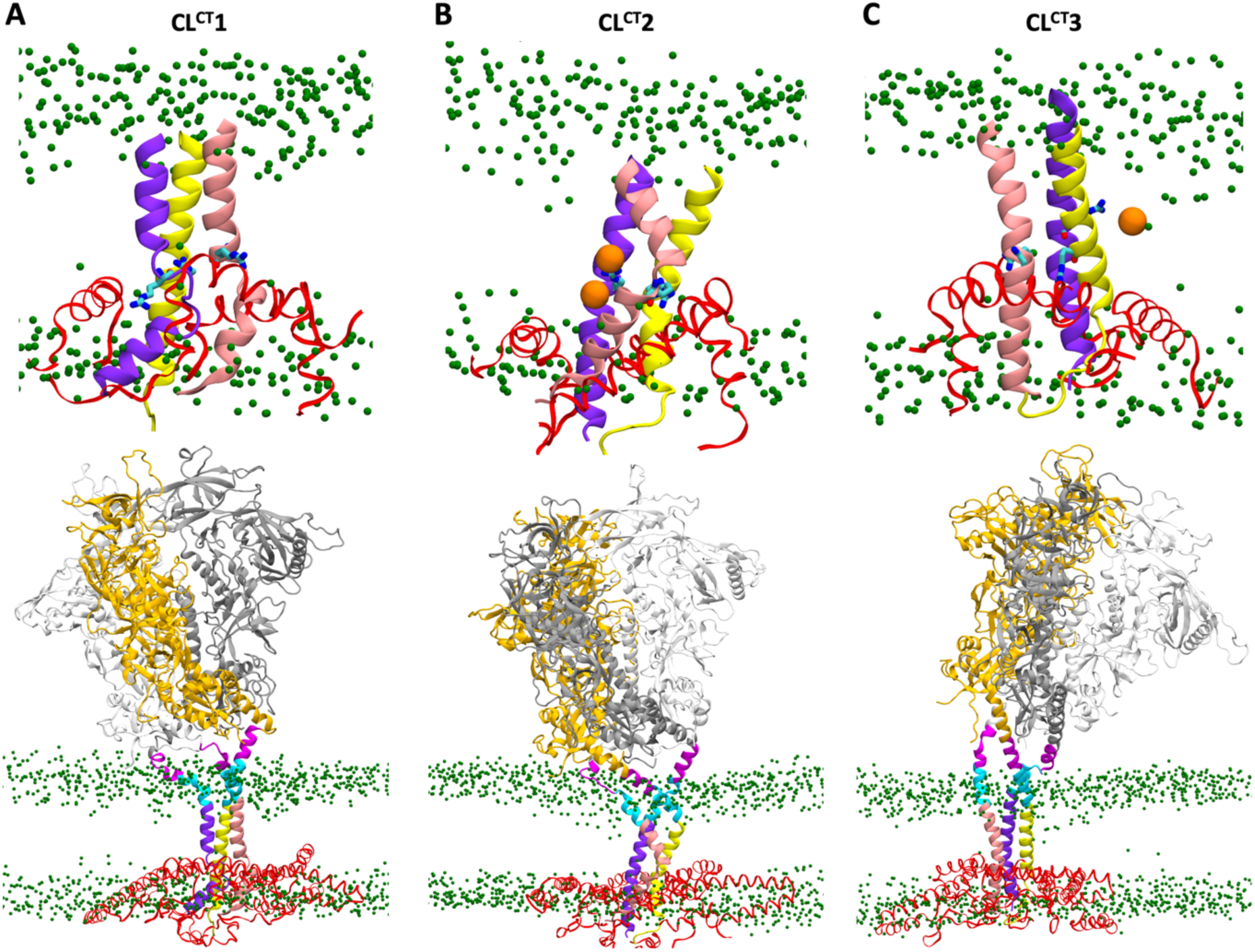
Local conformation of the TMD and global conformation of protein and membrane (CL^CT^ systems). Labeling and color coding are the same as in Figure 4—figure supplement 5.

**Figure 4—figure supplement 7.**
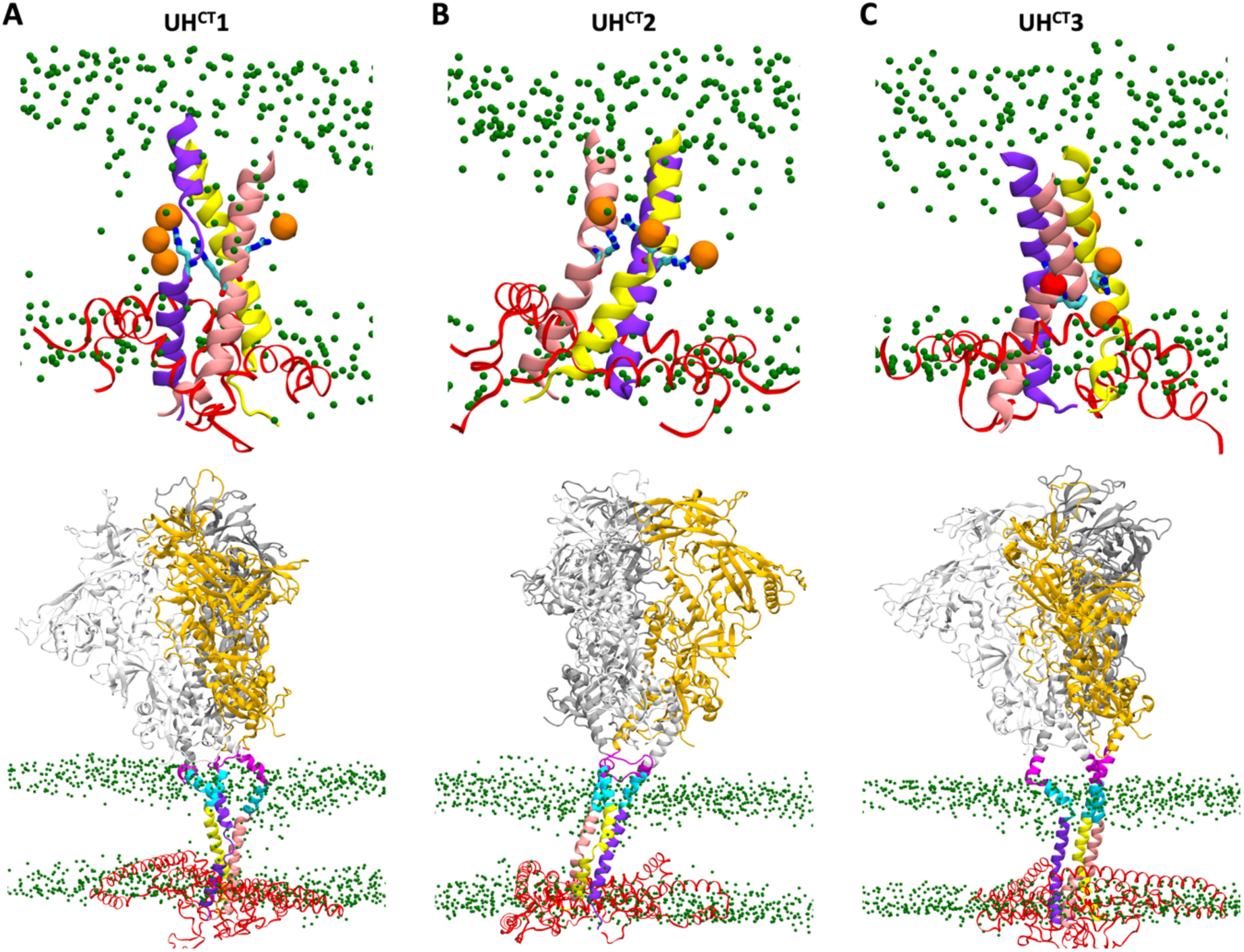
Local conformations of the TMD and global conformations of protein and membrane (UH^CT^ systems). Labeling and color coding are the same as in Figure 4—figure supplement 5.

**Figure 4—figure supplement 8.**
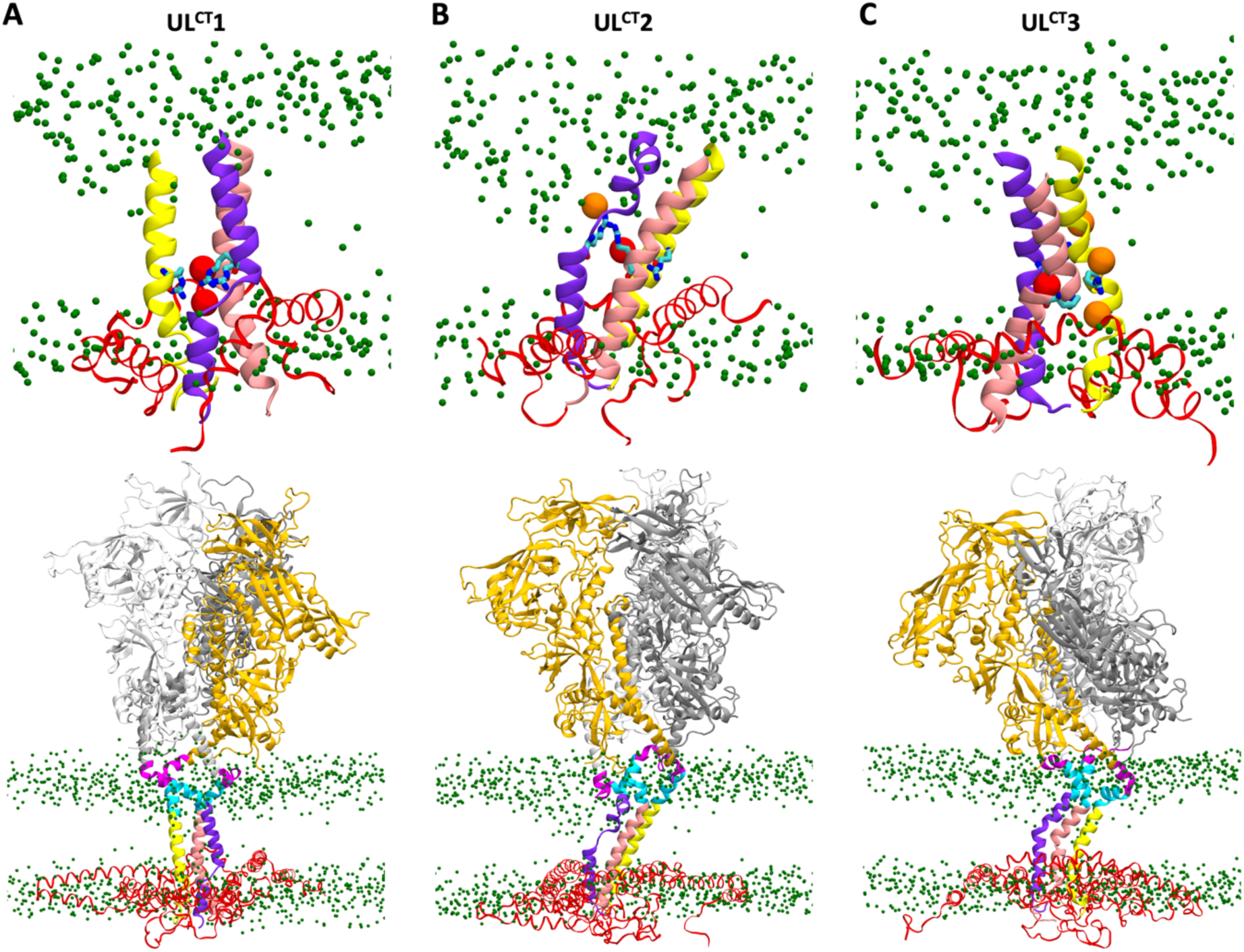
Local conformations of the TMD and global conformations of protein and membrane (UL^CT^ systems). Labeling and color coding are the same as in Figure 4—figure supplement 5.

**Figure 5—figure supplement 1.**
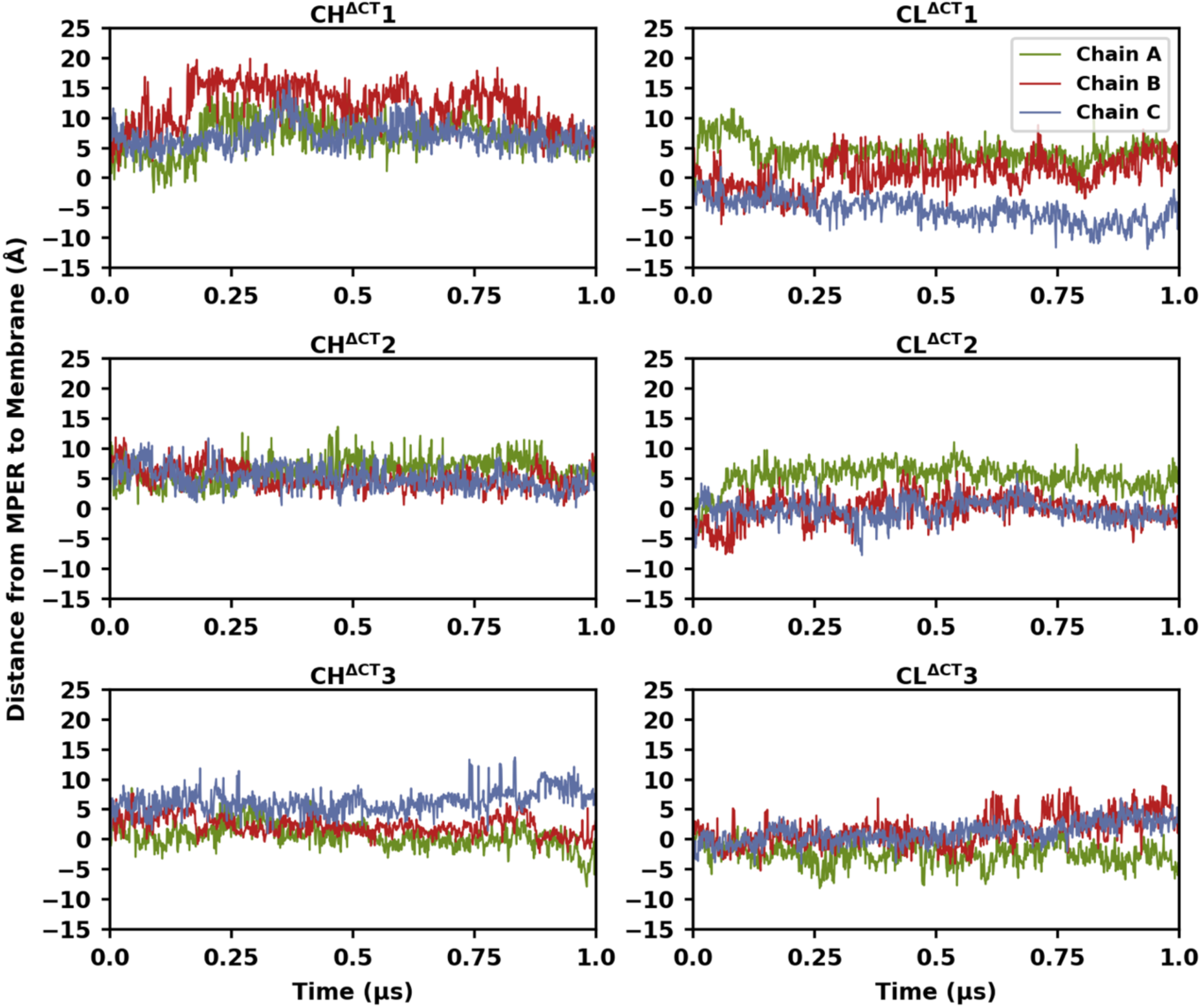
Temporal evolution of the distance from the MPER midpoint to the membrane surface. The distance *d*_F673_, defined in Figure 5A, is shown as a function of simulation time for the CH^ΔCT^ and CL^ΔCT^ systems.

**Figure 6—figure supplement 1.**
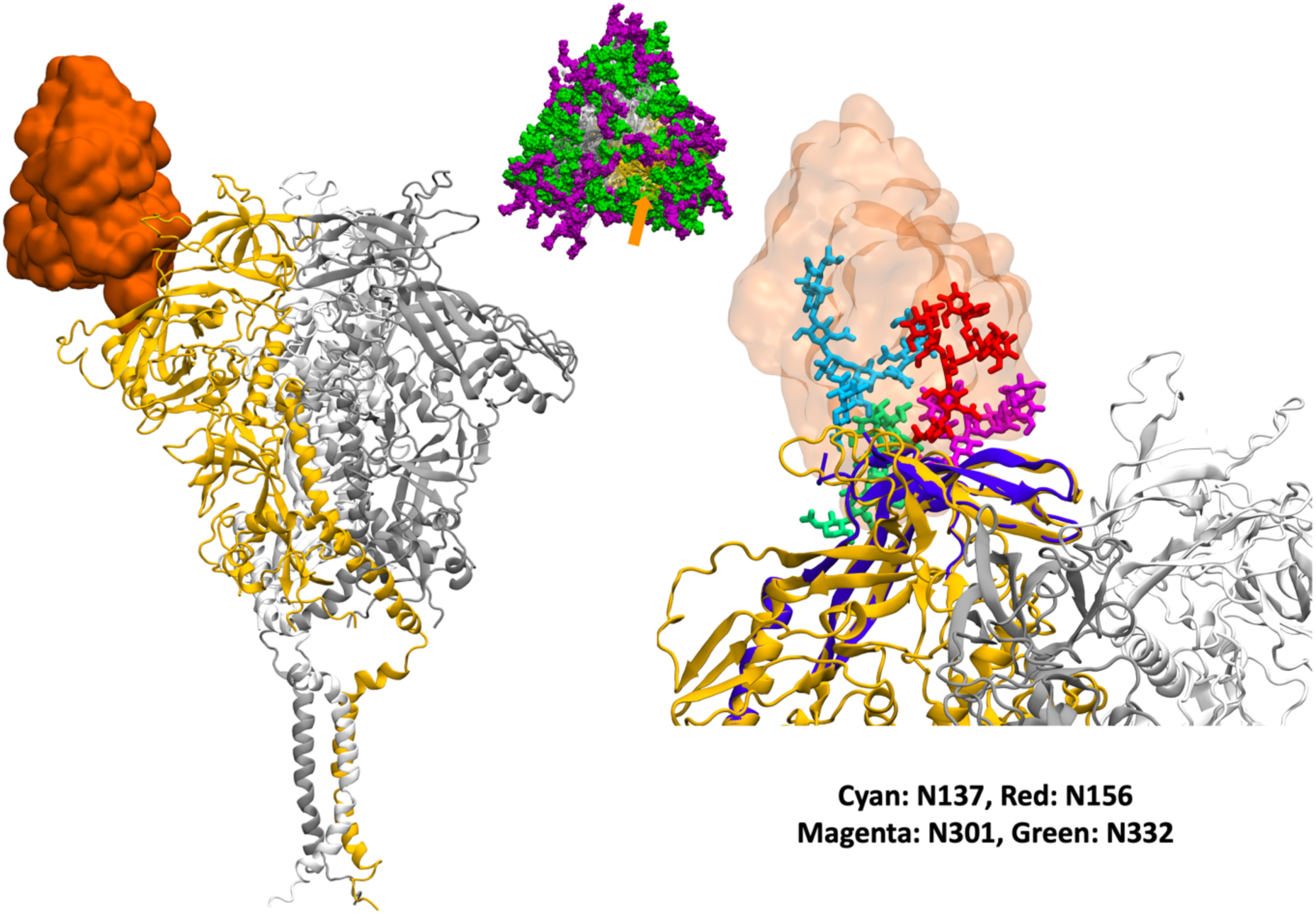
Shielding of antibody PGT128 epitope. (**Left**) Variable domains of the heavy and light chains aligned onto our modeled structure. (**Middle**) Top view of the glycosylated trimeric protein, with the orange arrow indicating the epitope location. (**Right**) Part of the epitope in the antibody-epitope complex PDB structure was used for structural alignment and is highlighted in purple. Glycans capable of hindering antibody binding are shown in various colors.

**Figure 6—figure supplement 2.**
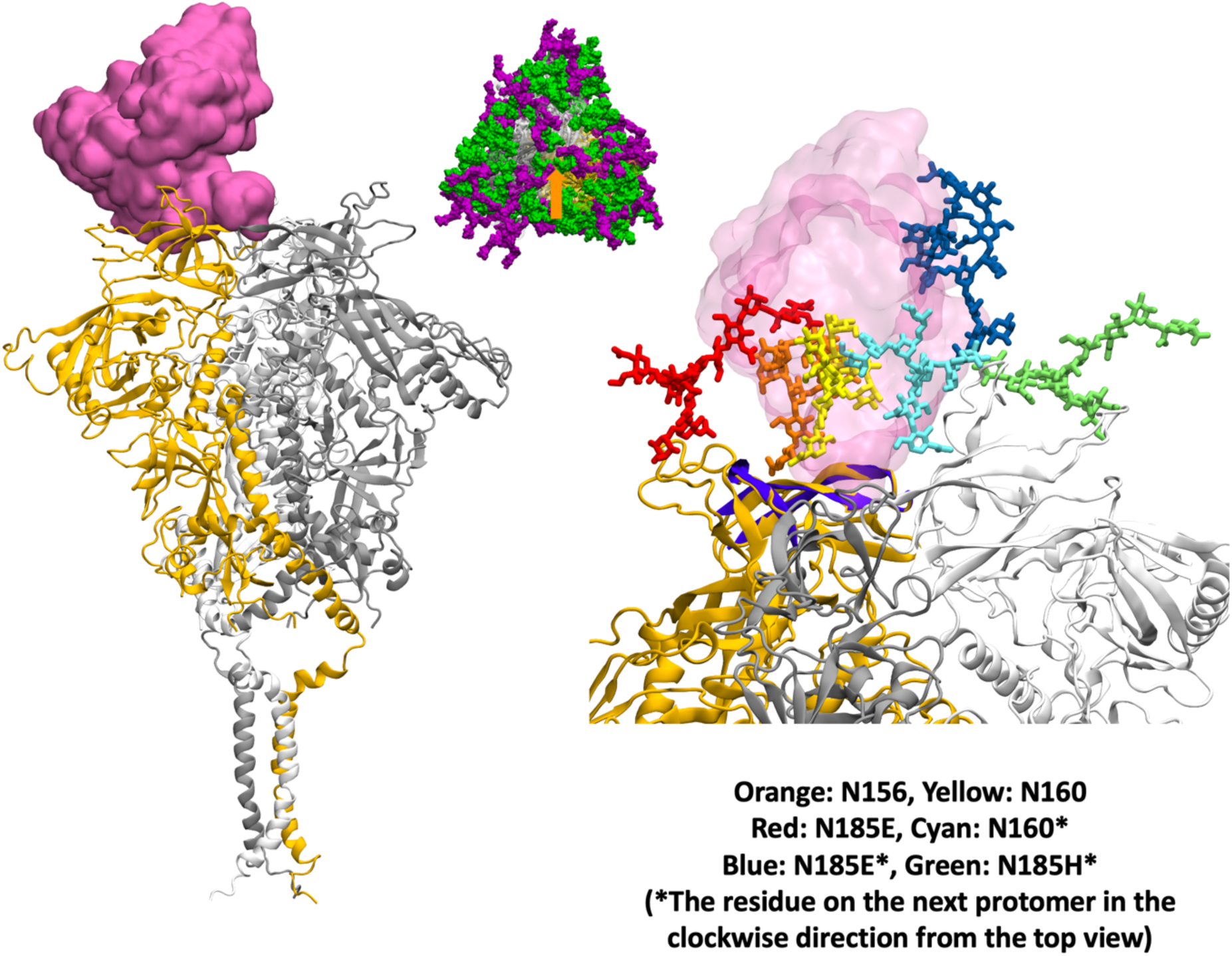
Shielding of antibody PG9 epitope. Labeling and color coding are the same as in Figure 6—figure supplement 1.

**Figure 6—figure supplement 3.**
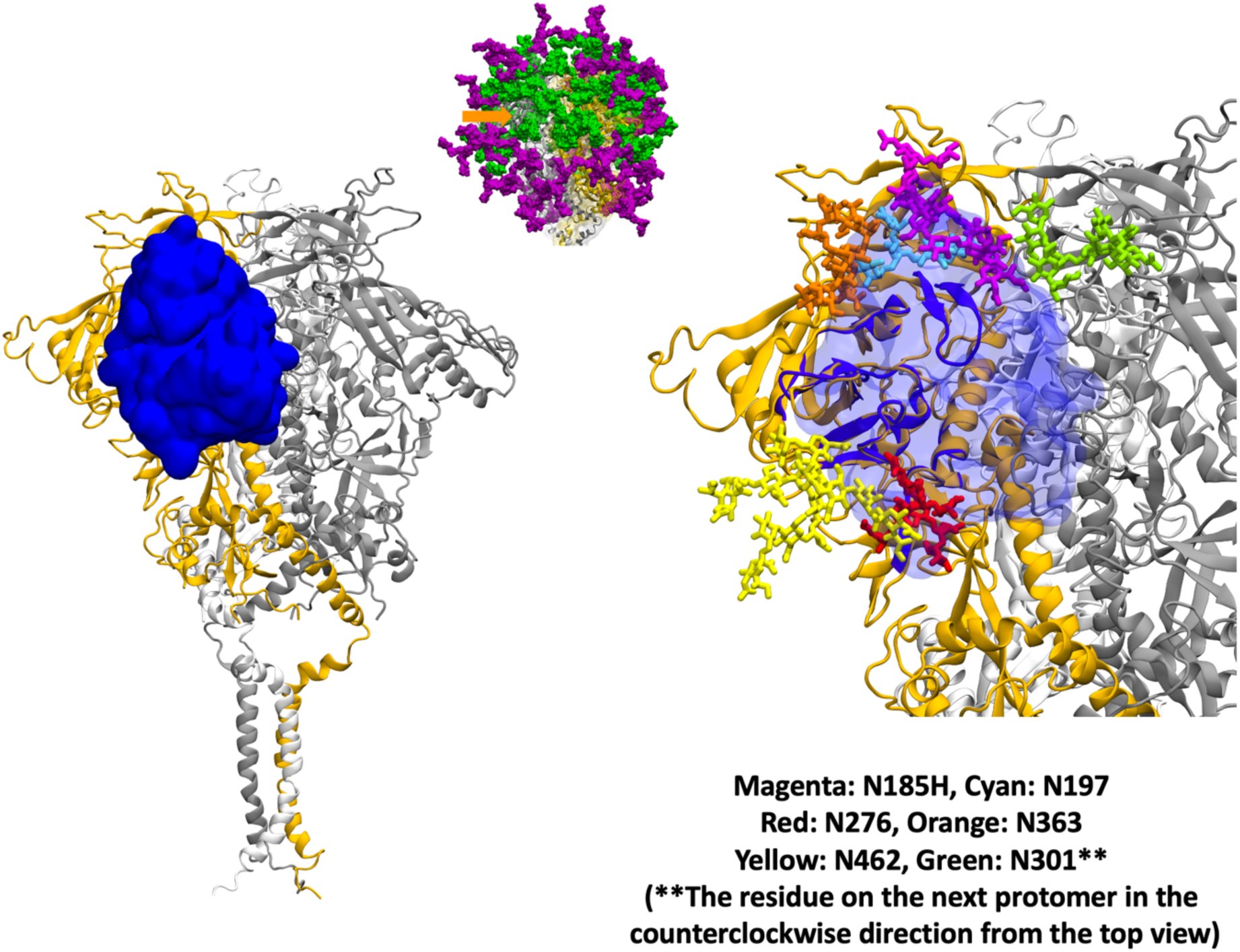
Shielding of antibody VRC01 epitope. Labeling and color coding are the same as in Figure 6—figure supplement 1. (**Middle**) The glycosylated trimeric protein is shown in a side view instead of the top view.

**Figure 6—figure supplement 4.**
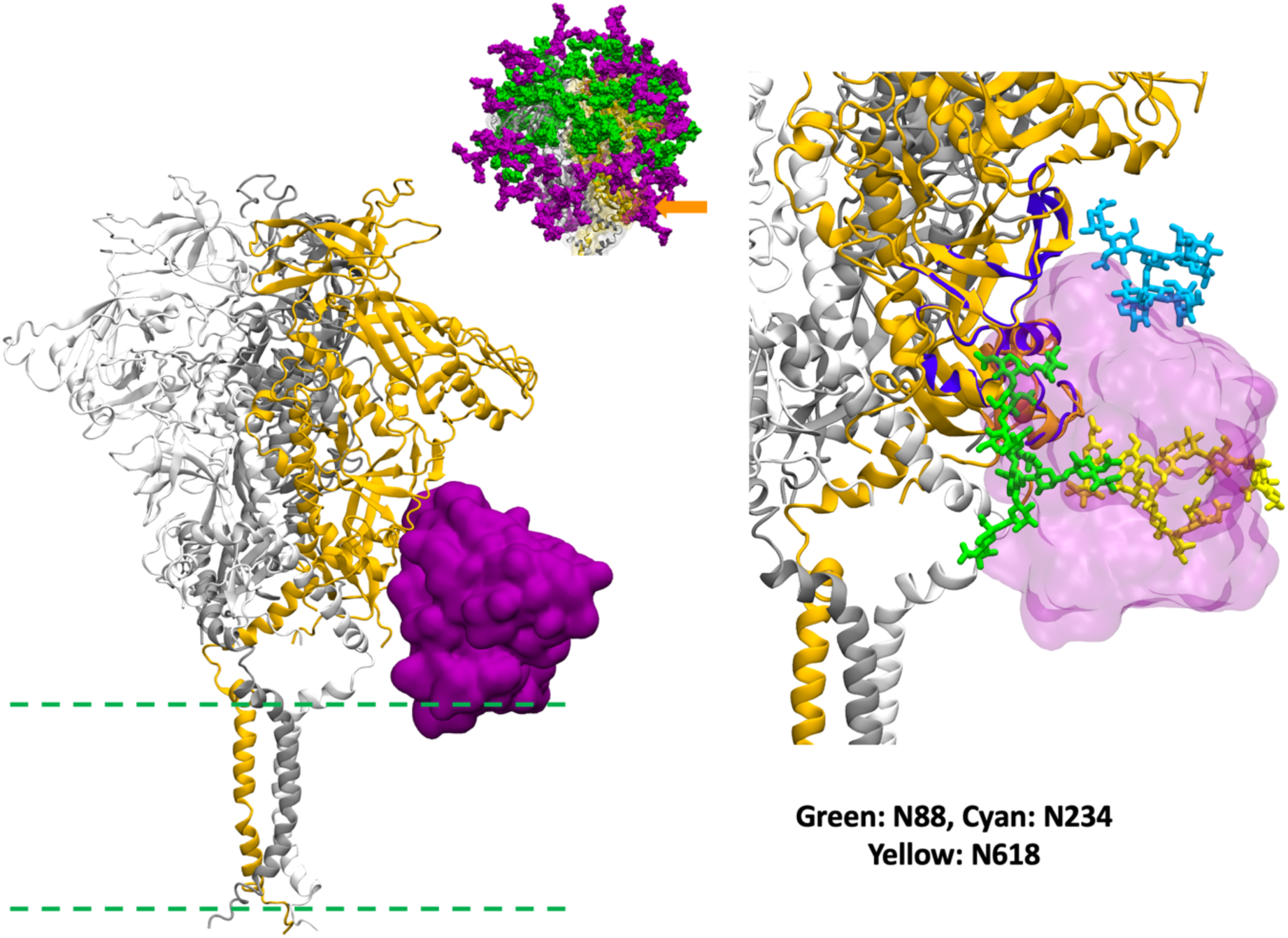
Shielding of antibody 35O22 epitope. Labeling and color coding are the same as in Figure 6—figure supplement 3. (**Left**) The dashed greens indicate the approximate location of the lipid headgroups.

**Figure 6—figure supplement 5.**
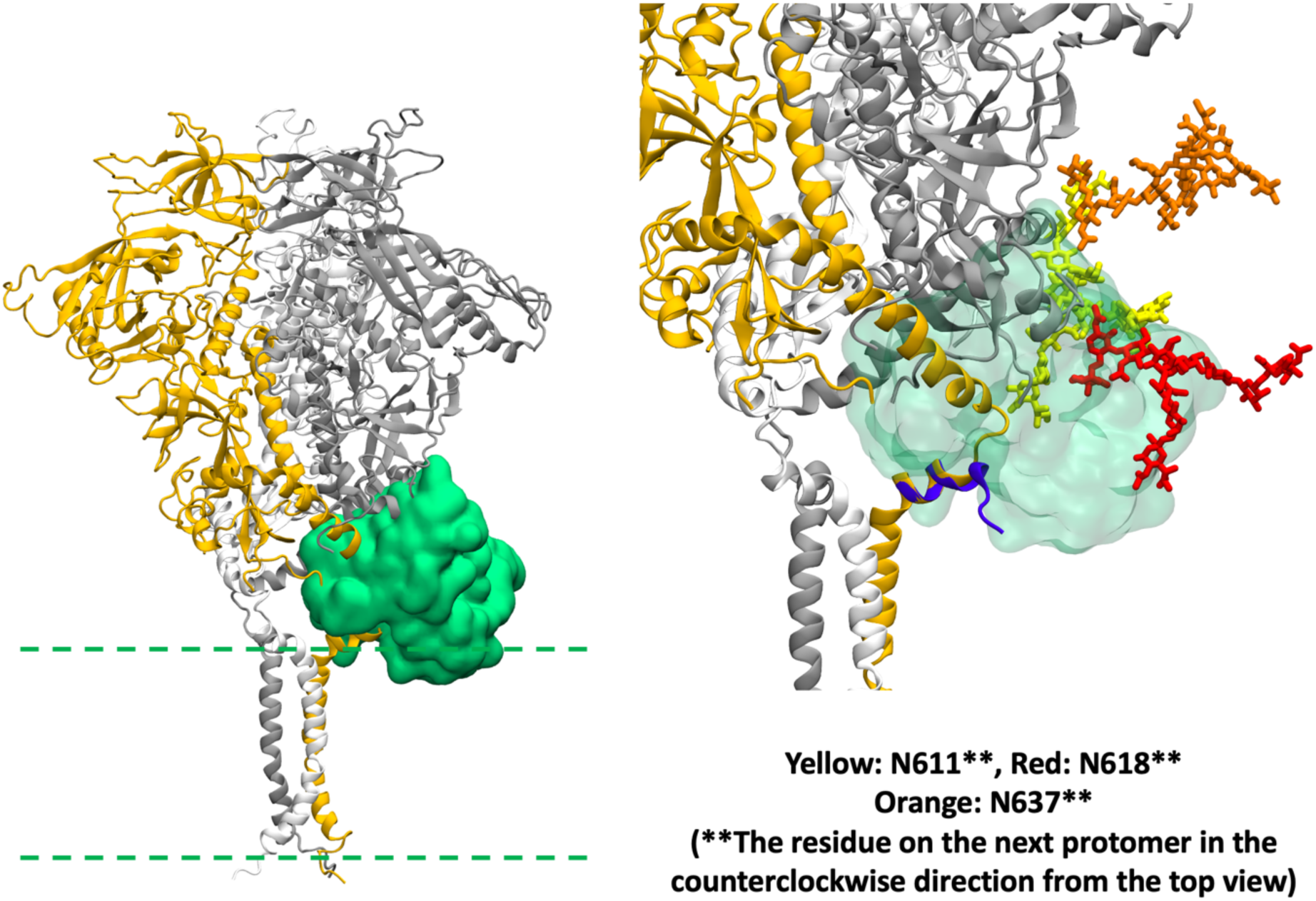
Shielding of antibody 4E10 epitope. Labeling and color coding are the same as in Figure 6—figure supplement 4.

**Figure 6—figure supplement 6.**
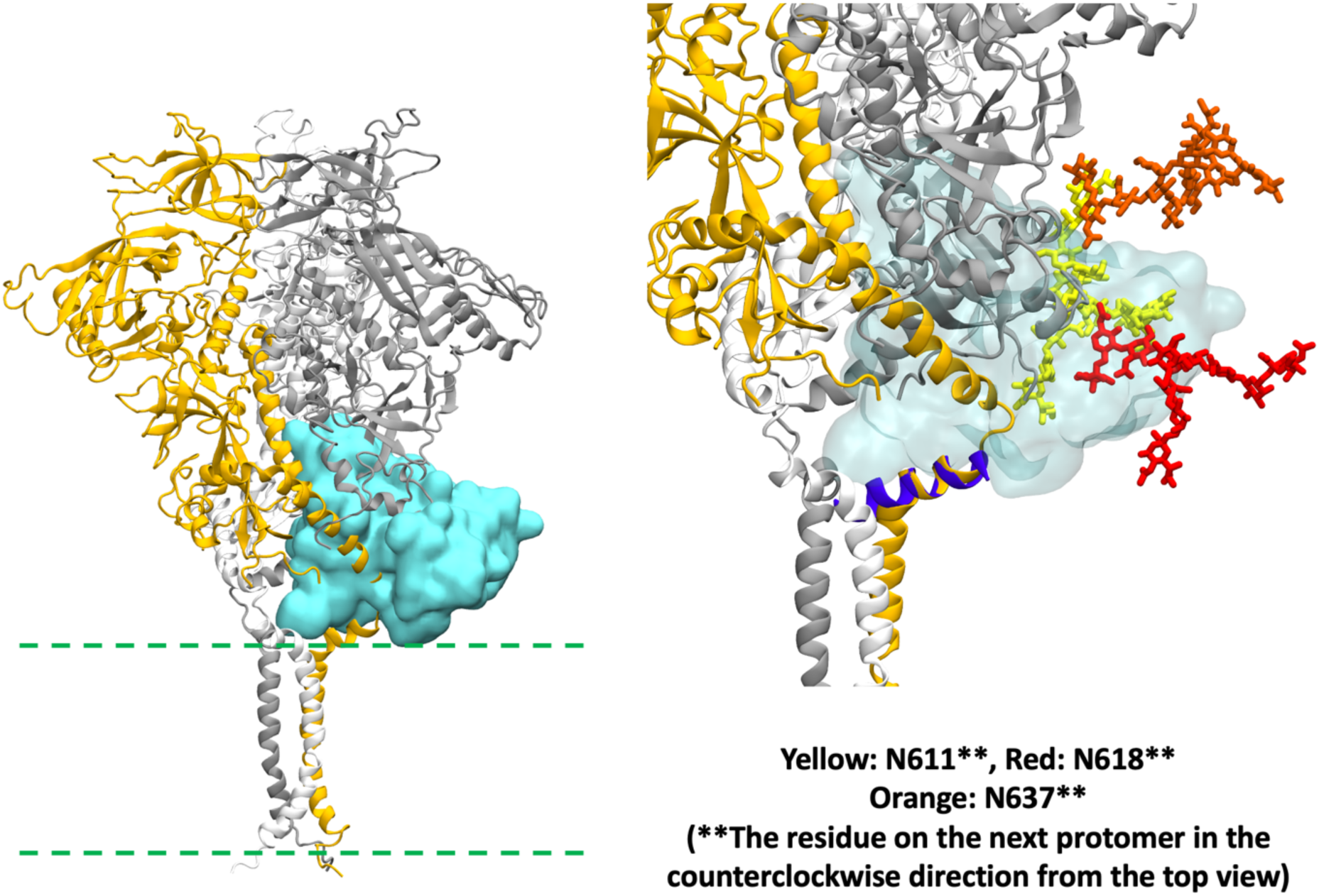
Shielding of antibody 10E8 epitope. Labeling and color coding are the same as in Figure 6—figure supplement 4.

**Figure 6—figure supplement 7.**
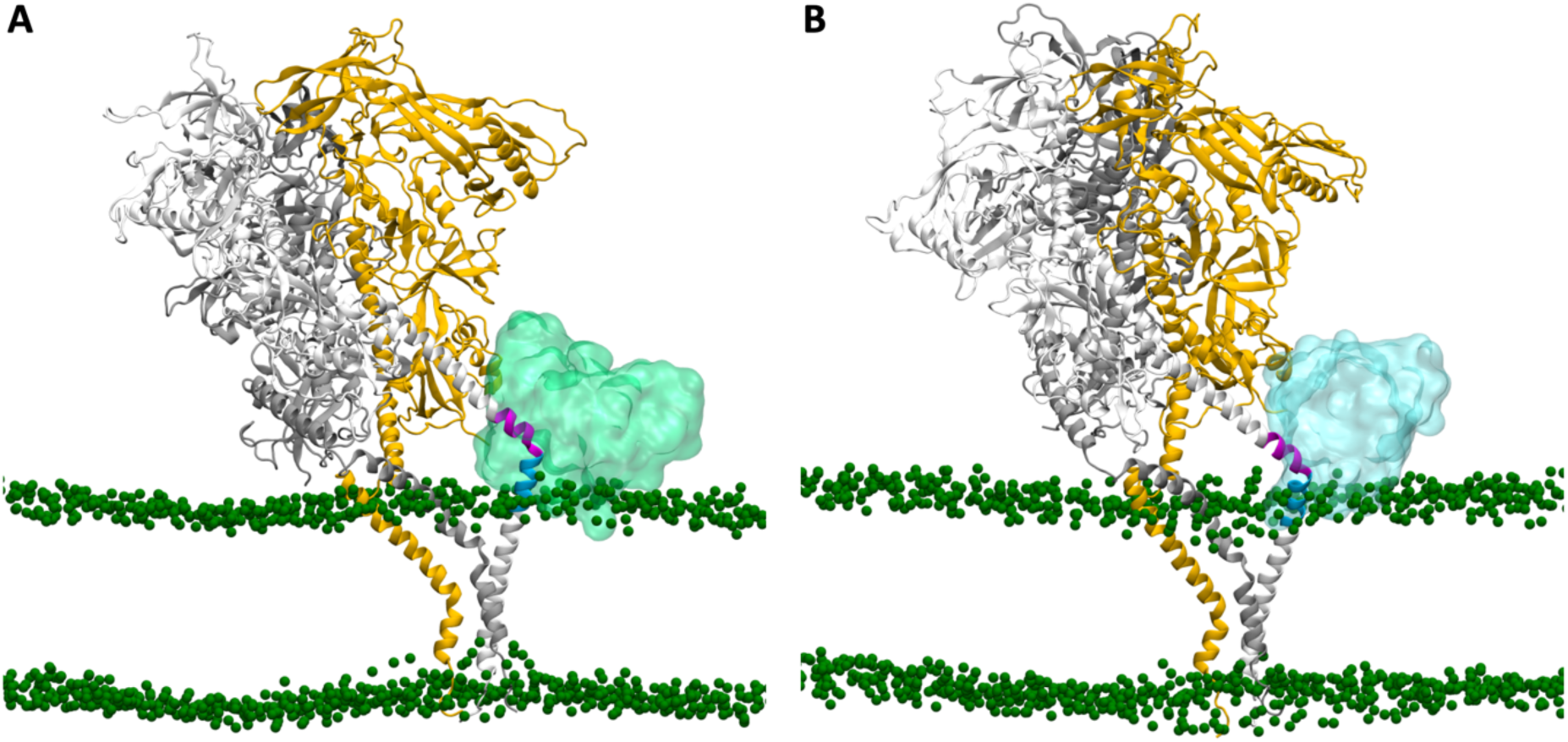
Snapshots showing the MPER of one protomer accessible to either 4E10 or 10E8. (**A**) Snapshot in which the MPER of the white protomer is accessible to 4E10 (cyan transparent surface) but not to 10E8. (**B**) Snapshot in which the MPER of the white protomer is accessible to 10E8 (green transparent surface) but not to 4E10.

**Figure 6—figure supplement 8.**
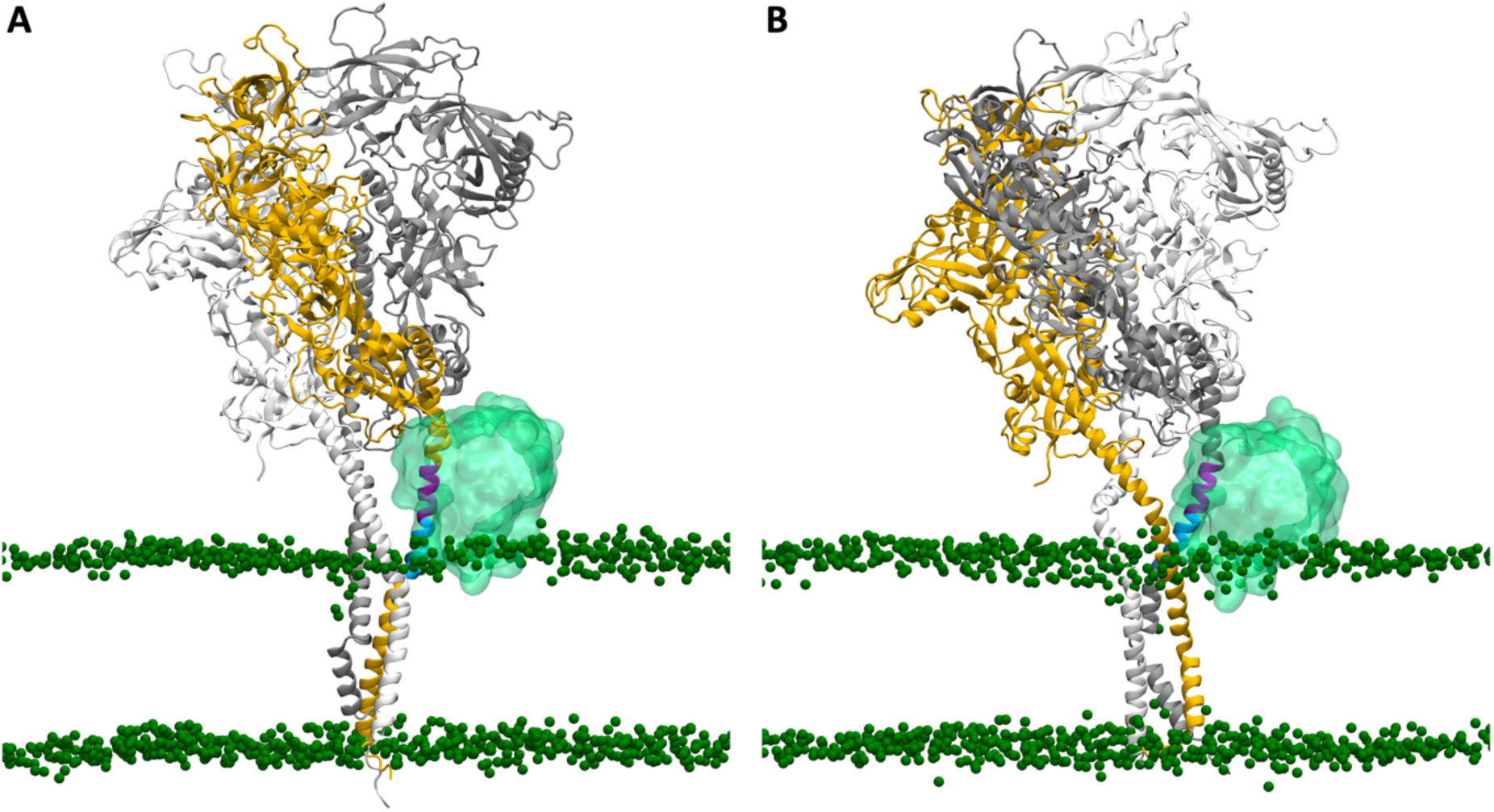
Snapshots showing the MPER of two protomers are accessible to 4E10. The MPER is accessible to 4E10, (**A**) exclusively in the yellow protomer, and (**B**) exclusively in the gray protomer.

**Figure 6—figure supplement 9.**
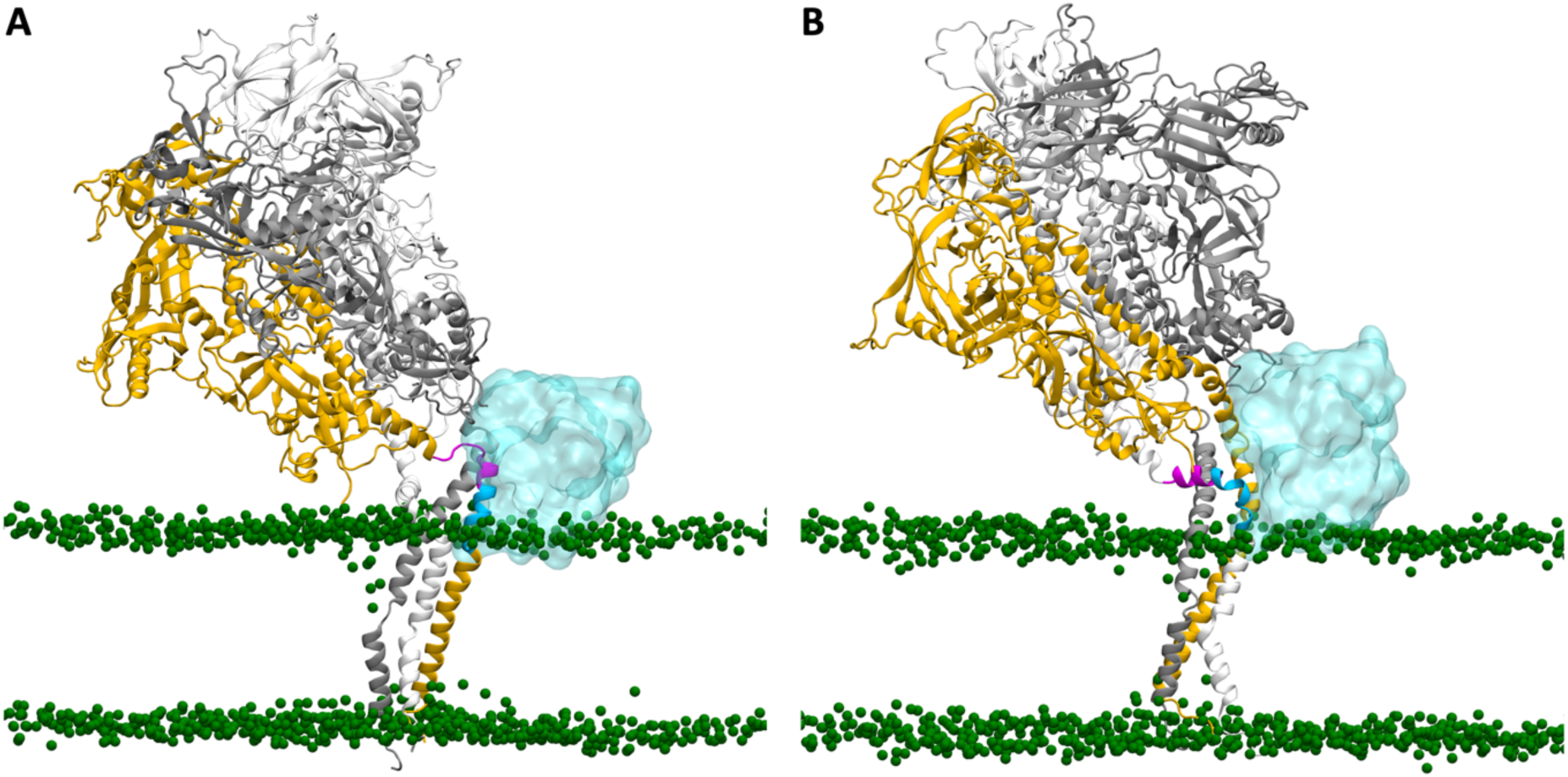
Snapshots showing the MPER of two protomers are accessible to 10E8. The MPER is accessible to 4E10, (**A**) exclusively in the yellow protomer, and (**B**) exclusively in the white protomer.

**Figure 6—figure supplement 10.**
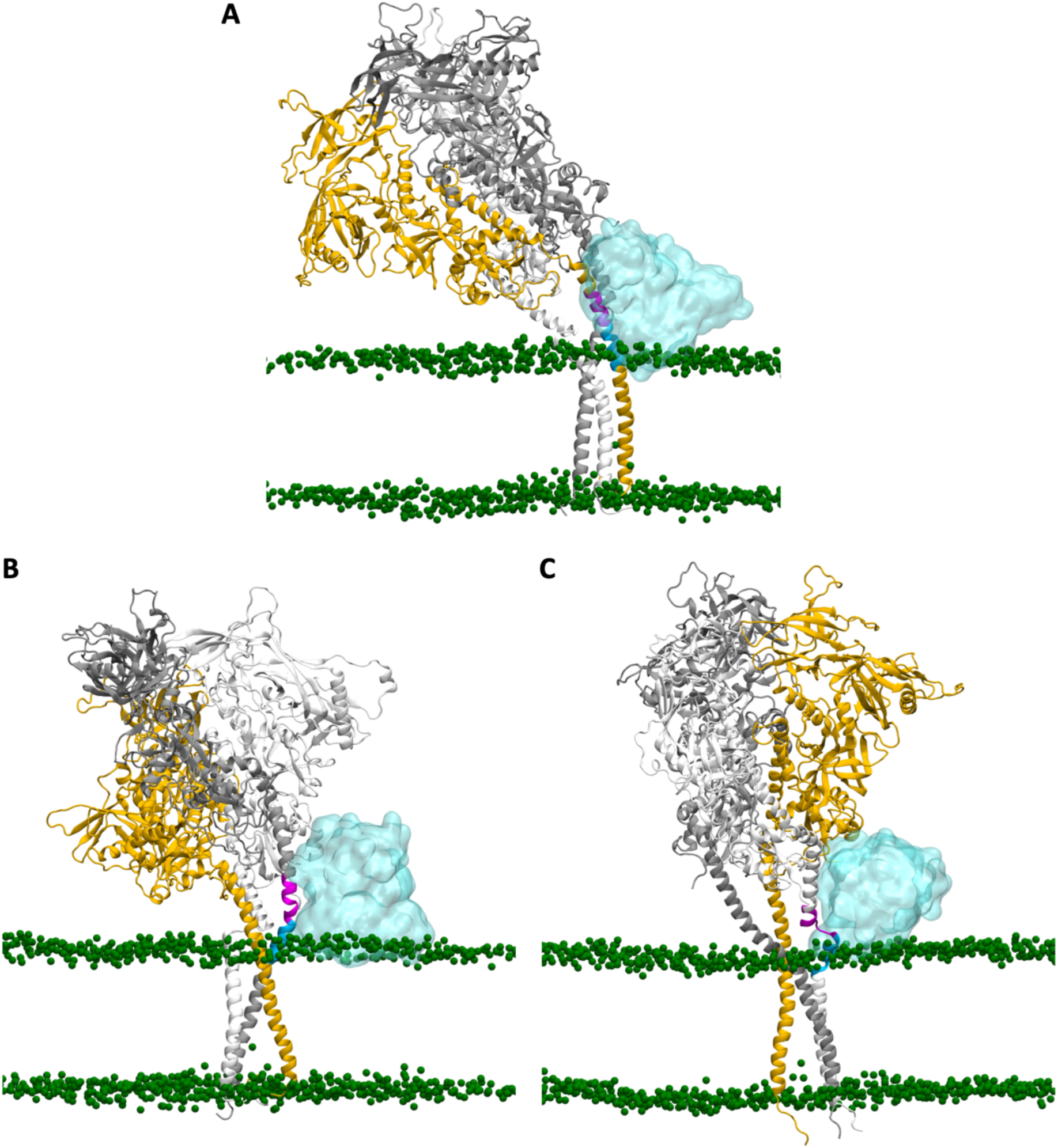
Snapshots showing the MPER of three protomers are accessible to 10E8. The MPER is accessible to 4E10, (**A**) exclusively in the yellow protomer, (**B**) exclusively in the gray protomer, and (**C**) exclusively in the white protomer.

