## Supplementary file for "Conformational Variability of HIV-1 Env Trimer and Viral Vulnerability"

| Name | Cleaved | TMD Position | CT Truncated | Number of Atoms | Simulation Time | Number of Simulation Runs |
|---|---|---|---|---|---|---|
| $CH^{\Delta CT}$ | Yes | High | Yes | 1,046,227 | 1 μs | 3 |
| $CL^{\Delta CT}$ | Yes | Low | Yes | 1,040,888 | 1 μs | 3 |
| $UH^{\Delta CT}$ | No | High | Yes | 1,080,915 | 1 μs | 3 |
| $UL^{\Delta CT}$ | No | Low | Yes | 1,080,164 | 1 μs | 3 |
| $CH^{CT}$ | Yes | High | No | 1,080,113 | 1 μs | 3 |
| $CL^{CT}$ | Yes | Low | No | 1,078,556 | 1 μs | 3 |
| $UH^{CT}$ | No | High | No | 1,118,568 | 1 μs | 3 |
| $UL^{CT}$ | No | Low | No | 1,118,375 | 1 μs | 3 |

**Supplementary Table 1. Simulation system information.**

| | $CH^{\Delta CT}1$ | $CH^{\Delta CT}2$ | $CH^{\Delta CT}3$ | $CL^{\Delta CT}1$ | $CL^{\Delta CT}2$ | $CL^{\Delta CT}3$ |
|---|---|---|---|---|---|---|
| A[1] | 19% | 7%[4] | 36% | 21% | 16% | 1% |
| B[1] | 76% | 1% | 15% | 19% | 3% | 46% |
| C[1] | 43% | 13% | 17% | 19% | 14% | 19% |
| A&B[2] | 17% | 0% | 8% | 8% | 1% | 0% |
| A&C[2] | 6% | 0% | 1% | 6% | 0% | 0% |
| B&C[2] | 37% | 0% | 3% | 2% | 0% | 14% |
| A&B&C[3] | 6% | 0% | 0% | 1% | 0% | 0% |

| | $UH^{\Delta CT}1$ | $UH^{\Delta CT}2$ | $UH^{\Delta CT}3$ | $UL^{\Delta CT}1$ | $UL^{\Delta CT}2$ | $UL^{\Delta CT}3$ |
|---|---|---|---|---|---|---|
| A | 5% | 32% | 2% | 3% | 9% | 5% |
| B | 35% | 38% | 41% | 2% | 28% | 2% |
| C | 12% | 16% | 53% | 45% | 3% | 33% |
| A&B | 1% | 15% | 1% | 0% | 5% | 0% |
| A&C | 1% | 9% | 1% | 1% | 1% | 2% |
| B&C | 4% | 10% | 20% | 1% | 2% | 1% |
| A&B&C | 1% | 5% | 1% | 0% | 1% | 0% |

| | $CH^{CT}1$ | $CH^{CT}2$ | $CH^{CT}3$ | $CL^{CT}1$ | $CL^{CT}2$ | $CL^{CT}3$ |
|---|---|---|---|---|---|---|
| A | 7% | 10% | 9% | 15% | 0% | 7% |
| B | 2% | 15% | 19% | 22% | 0% | 0% |
| C | 44% | 2% | 25% | 14% | 4% | 23% |
| A&B | 0% | 2% | 0% | 3% | 0% | 0% |
| A&C | 1% | 0% | 1% | 1% | 0% | 1% |
| B&C | 1% | 0% | 9% | 5% | 0% | 0% |
| A&B&C | 0% | 0% | 0% | 1% | 0% | 0% |

| | $UH^{CT}1$ | $UH^{CT}2$ | $UH^{CT}3$ | $UL^{CT}1$ | $UL^{CT}2$ | $UL^{CT}3$ |
|---|---|---|---|---|---|---|
| A | 1% | 1% | 26% | 33% | 1% | 2% |
| B | 2% | 23% | 26% | 45% | 67% | 48% |
| C | 4% | 34% | 13% | 3% | 38% | 17% |
| A&B | 0% | 1% | 5% | 14% | 1% | 0% |
| A&C | 0% | 1% | 1% | 2% | 1% | 1% |
| B&C | 0% | 9% | 3% | 2% | 29% | 1% |
| A&B&C | 0% | 1% | 0% | 1% | 1% | 0% |

**Supplementary Table 2. Frequency of epitope accessibility for antibody PGT128.** [1]Three protein chains are referred to as A, B, and C. [2]"A&B" denotes that the epitopes on both chains A and B are concurrently accessible. [3]"A&B&C" denotes that the epitopes on all three chains are concurrently accessible. [4]Red text highlights cases where the frequency of epitope accessibility on an individual chain is <10%. The same notations apply for Supplementary Table 3 and 4.

| | $CH^{\Delta CT}1$ | $CH^{\Delta CT}2$ | $CH^{\Delta CT}3$ | $CL^{\Delta CT}1$ | $CL^{\Delta CT}2$ | $CL^{\Delta CT}3$ |
|---|---|---|---|---|---|---|
| A | 0% | 0% | 0% | 0% | 0% | 0% |
| B | 20% | 34% | 1% | 11% | 0% | 15% |
| C | 0% | 0% | 1% | 0% | 0% | 0% |
| A&B | 0% | 0% | 0% | 0% | 0% | 0% |
| A&C | 0% | 0% | 0% | 0% | 0% | 0% |
| B&C | 0% | 0% | 0% | 0% | 0% | 0% |
| A&B&C | 0% | 0% | 0% | 0% | 0% | 0% |

| | $UH^{\Delta CT}1$ | $UH^{\Delta CT}2$ | $UH^{\Delta CT}3$ | $UL^{\Delta CT}1$ | $UL^{\Delta CT}2$ | $UL^{\Delta CT}3$ |
|---|---|---|---|---|---|---|
| A | 1% | 0% | 0% | 0% | 1% | 0% |
| B | 0% | 1% | 2% | 0% | 1% | 0% |
| C | 2% | 0% | 0% | 0% | 0% | 0% |
| A&B | 0% | 0% | 0% | 0% | 0% | 0% |
| A&C | 0% | 0% | 0% | 0% | 0% | 0% |
| B&C | 0% | 0% | 0% | 0% | 0% | 0% |
| A&B&C | 0% | 0% | 0% | 0% | 0% | 0% |

| | $CH^{CT}1$ | $CH^{CT}2$ | $CH^{CT}3$ | $CL^{CT}1$ | $CL^{CT}2$ | $CL^{CT}3$ |
|---|---|---|---|---|---|---|
| A | 0% | 0% | 0% | 0% | 0% | 0% |
| B | 1% | 0% | 2% | 3% | 0% | 31% |
| C | 0% | 0% | 0% | 0% | 5% | 0% |
| A&B | 0% | 0% | 0% | 0% | 0% | 0% |
| A&C | 0% | 0% | 0% | 0% | 0% | 0% |
| B&C | 0% | 0% | 0% | 0% | 0% | 0% |
| A&B&C | 0% | 0% | 0% | 0% | 0% | 0% |

| | $UH^{CT}1$ | $UH^{CT}2$ | $UH^{CT}3$ | $UL^{CT}1$ | $UL^{CT}2$ | $UL^{CT}3$ |
|---|---|---|---|---|---|---|
| A | 0% | 1% | 1% | 0% | 0% | 0% |
| B | 0% | 1% | 0% | 0% | 0% | 0% |
| C | 0% | 1% | 0% | 0% | 2% | 0% |
| A&B | 0% | 0% | 0% | 0% | 0% | 0% |
| A&C | 0% | 0% | 0% | 0% | 0% | 0% |
| B&C | 0% | 0% | 0% | 0% | 0% | 0% |
| A&B&C | 0% | 0% | 0% | 0% | 0% | 0% |

**Supplementary Table 3. Frequency of epitope accessibility for antibody PG9.**

| | $CH^{\Delta CT}1$ | $CH^{\Delta CT}2$ | $CH^{\Delta CT}3$ | $CL^{\Delta CT}1$ | $CL^{\Delta CT}2$ | $CL^{\Delta CT}3$ |
|---|---|---|---|---|---|---|
| A | 0% | 0% | 10% | 18% | 0% | 4% |
| B | 2% | 2% | 5% | 4% | 20% | 5% |
| C | 41% | 3% | 1% | 0% | 3% | 4% |
| A&B | 0% | 0% | 0% | 0% | 0% | 0% |
| A&C | 0% | 0% | 0% | 0% | 0% | 0% |
| B&C | 0% | 0% | 0% | 0% | 1% | 0% |
| A&B&C | 0% | 0% | 0% | 0% | 0% | 0% |

| | $UH^{\Delta CT}1$ | $UH^{\Delta CT}2$ | $UH^{\Delta CT}3$ | $UL^{\Delta CT}1$ | $UL^{\Delta CT}2$ | $UL^{\Delta CT}3$ |
|---|---|---|---|---|---|---|
| A | 15% | 0% | 1% | 7% | 15% | 4% |
| B | 4% | 0% | 1% | 0% | 7% | 13% |
| C | 24% | 5% | 32% | 33% | 6% | 45% |
| A&B | 0% | 0% | 0% | 0% | 2% | 0% |
| A&C | 7% | 0% | 1% | 1% | 1% | 1% |
| B&C | 0% | 0% | 0% | 0% | 0% | 10% |
| A&B&C | 0% | 0% | 0% | 0% | 0% | 0% |

| | $CH^{CT}1$ | $CH^{CT}2$ | $CH^{CT}3$ | $CL^{CT}1$ | $CL^{CT}2$ | $CL^{CT}3$ |
|---|---|---|---|---|---|---|
| A | 14% | 13% | 0% | 0% | 2% | 9% |
| B | 9% | 3% | 1% | 4% | 2% | 3% |
| C | 9% | 11% | 10% | 0% | 0% | 24% |
| A&B | 2% | 0% | 0% | 0% | 0% | 2% |
| A&C | 1% | 2% | 0% | 0% | 0% | 3% |
| B&C | 1% | 0% | 0% | 0% | 0% | 0% |
| A&B&C | 0% | 0% | 0% | 0% | 0% | 0% |

| | $UH^{CT}1$ | $UH^{CT}2$ | $UH^{CT}3$ | $UL^{CT}1$ | $UL^{CT}2$ | $UL^{CT}3$ |
|---|---|---|---|---|---|---|
| A | 17% | 0% | 16% | 4% | 22% | 5% |
| B | 6% | 0% | 1% | 0% | 0% | 0% |
| C | 57% | 19% | 57% | 35% | 14% | 44% |
| A&B | 2% | 0% | 0% | 0% | 0% | 0% |
| A&C | 6% | 0% | 11% | 2% | 2% | 0% |
| B&C | 2% | 0% | 1% | 0% | 0% | 0% |
| A&B&C | 0% | 0% | 0% | 0% | 0% | 0% |

**Supplementary Table 4. Frequency of epitope accessibility for antibody VRC01.**

| | $CH^{\Delta CT}1$ | $CH^{\Delta CT}2$ | $CH^{\Delta CT}3$ | $CL^{\Delta CT}1$ | $CL^{\Delta CT}2$ | $CL^{\Delta CT}3$ |
|---|---|---|---|---|---|---|
| A | 7%, 0%[1] | 4%, 0% | 9%, 8% | 3%, 0% | 24%, 0% | 7%, 4% |
| B | 29%, 2%[2] | 14%, 0% | 12%, 0% | 0%, 0% | 10%, 10% | 0%, 0% |
| C | 8%, 8% | 14%, 10% | 10%, 5% | 32%, 7% | 4%, 1% | 22%, 16% |
| A&B | 7%, 0% | 4%, 0% | 9%, 0% | 0%, 0% | 10%, 0% | 0%, 0% |
| A&C | 7%, 0% | 4%, 0% | 9%, 5% | 3%, 0% | 4%, 0% | 7%, 4% |
| B&C | 8%, 2% | 14%, 0% | 10%, 0% | 0%, 0% | 4%, 1% | 0%, 0% |
| A&B&C | 7%, 0% | 4%, 0% | 9%, 0% | 0%, 0% | 4%, 0% | 0%, 0% |

| | $UH^{\Delta CT}1$ | $UH^{\Delta CT}2$ | $UH^{\Delta CT}3$ | $UL^{\Delta CT}1$ | $UL^{\Delta CT}2$ | $UL^{\Delta CT}3$ |
|---|---|---|---|---|---|---|
| A | 16%, 12% | 49%, 1% | 9%, 7% | 14%, 5% | 23%, 19% | 43%, 10% |
| B | 2%, 2% | 5%, 5% | 10%, 9% | 7%, 7% | 3%, 1% | 5%, 0% |
| C | 22%, 13% | 25%, 25% | 23%, 23% | 26%, 3% | 6%, 5% | 5%, 5% |
| A&B | 2%, 2% | 5%, 1% | 9%, 7% | 7%, 5% | 3%, 1% | 5%, 0% |
| A&C | 16%, 12% | 25%, 1% | 9%, 7% | 14%, 3% | 6%, 5% | 5%, 5% |
| B&C | 2%, 2% | 5%, 5% | 10%, 9% | 7%, 3% | 3%, 1% | 5%, 0% |
| A&B&C | 2%, 2% | 5%, 1% | 9%, 7% | 7%, 3% | 3%, 1% | 5%, 0% |

| | $CH^{CT}1$ | $CH^{CT}2$ | $CH^{CT}3$ | $CL^{CT}1$ | $CL^{CT}2$ | $CL^{CT}3$ |
|---|---|---|---|---|---|---|
| A | 9%, 0% | 30%, 0% | 48%, 18% | 12%, 4% | 22%, 0% | 5%, 5% |
| B | 0%, 0% | 0%, 0% | 31%, 8% | 11%, 11% | 24%, 11% | 24%, 1% |
| C | 2%, 2% | 3%, 2% | 7%, 0% | 24%, 0% | 6%, 2% | 2%, 2% |
| A&B | 0%, 0% | 0%, 0% | 31%, 8% | 11%, 4% | 22%, 0% | 5%, 1% |
| A&C | 2%, 0% | 3%, 0% | 7%, 0% | 12%, 0% | 6%, 0% | 2%, 2% |
| B&C | 0%, 0% | 0%, 0% | 7%, 0% | 11%, 0% | 6%, 2% | 2%, 1% |
| A&B&C | 0%, 0% | 0%, 0% | 7%, 0% | 11%, 0% | 6%, 0% | 2%, 1% |

| | $UH^{CT}1$ | $UH^{CT}2$ | $UH^{CT}3$ | $UL^{CT}1$ | $UL^{CT}2$ | $UL^{CT}3$ |
|---|---|---|---|---|---|---|
| A | 5%, 4% | 18%, 16% | 33%, 0% | 31%, 0% | 31%, 0% | 41%, 0% |
| B | 26%, 16% | 8%, 5% | 11%, 7% | 2%, 1% | 2%, 0% | 18%, 1% |
| C | 25%, 0% | 7%, 0% | 20%, 9% | 3%, 2% | 29%, 15% | 7%, 7% |
| A&B | 5%, 4% | 8%, 5% | 11%, 0% | 2%, 0% | 2%, 0% | 18%, 0% |
| A&C | 5%, 0% | 7%, 0% | 20%, 0% | 3%, 0% | 29%, 0% | 7%, 0% |
| B&C | 25%, 0% | 7%, 0% | 11%, 7% | 2%, 1% | 2%, 0% | 7%, 1% |
| A&B&C | 5%, 0% | 7%, 0% | 11%, 0% | 2%, 0% | 2%, 0% | 7%, 0% |

**Supplementary Table 5. Frequency of epitope accessibility for antibody 35O22.** [1]The first value represents the frequency of epitope accessibility considering only glycan shielding, and the second value represents the frequency considering the shielding of both glycan and membrane. Red text highlights cases where the first value is <10%. [2]Magenta text highlights cases where the first value is ≥10% but the second value is <10%.

| Glycosylation Site | Composition and Type | Sequence |
|---|---|---|
| N197,<br>N355[1] | HexNAc(2)Hex(5)<br>High-mannose | 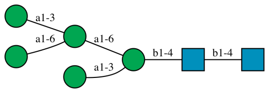 |
| N276[1] | HexNAc(2)Hex(7)<br>High-mannose | 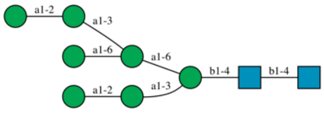 |
| N160[1] | HexNAc(2)Hex(8)<br>High-mannose | 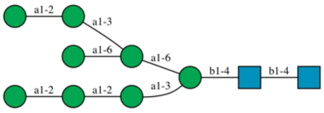 |
| N133, N156,<br>N234, N262,<br>N295, N301,<br>N332, N339,<br>N363, N386,<br>N392, N411,<br>N448 | HexNAc(2)Hex(9)<br>High-mannose | 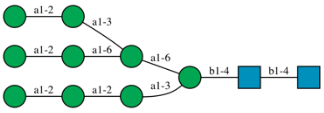 |
| N88, N137,<br>N185E, N185H,<br>N398, N406,<br>N618, N637[2] | HexNAc(4)Hex(5)<br>Fuc(1)Neu5Ac(2)<br>Complex | 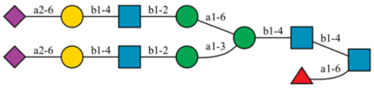 |
| N462,<br>N611[2,3] | HexNAc(5)Hex(6)<br>Fuc(1)Neu5Ac(3)<br>Complex | 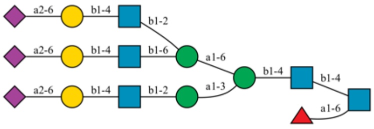 |

**Supplementary Table 6. Glycosylation sites, selected glycan compositions, and sequences used in this study.** [1]The same composition can correspond to multiple possible sequences, depending on which branch the mannose (Man) is attached to. In such cases, one representative sequence was selected for structural modeling. [2]N-Acetylneuraminic acid (Neu5Ac) can be either α2-3 or α2-6 linked to galactose (Gal). In this study, the α2-6 linkage was used for all non-reducing terminal Neu5Ac. [3]Both the α1-3 and α1-6 branches can contain one or two antennae. Here, we added one antenna to the α1-3 branch and two antennae to the α1-6 branch.

CT-truncated (ΔCT)

| Lipid Name | Lipid Head/Tail | Number of Lipids in Leaflets | |
|---|---|---|---|
| | | Exoplasmic | Cytoplasmic |
| POPC | PC (16:0/18:1(9Z)) | 120 | 56 |
| PLPC | PC (16:0/18:2(9Z,12Z)) | 165 | 88 |
| PAPE | PE (16:0/20:4(5Z,8Z,11Z,14Z)) | 23 | 96 |
| POPE | PE (16:0/18:1(9Z)) | 23 | 112 |
| POPI | PI (16:0/18:1(9Z)) | 0 | 40 |
| PAPS | PS (16:0/20:4(5Z,8Z,11Z,14Z)) | 0 | 88 |
| POPA | PA (16:0/18:1(9Z)) | 0 | 8 |
| SSM | SM (d18:1/18:0) | 83 | 40 |
| NSM | SM (d18:1/24:1) | 83 | 40 |
| CMH | GlcCer (d18:1/16:0) | 30 | 0 |
| CHOL | Cholesterol | 278 | 232 |
| TOTAL | | 805 | 800 |

Full-length (CT)

| Lipid Name | Lipid Head/Tail | Number of Lipids in Leaflets | |
|---|---|---|---|
| | | Exoplasmic | Cytoplasmic |
| POPC | PC (16:0/18:1(9Z)) | 120 | 49 |
| PLPC | PC (16:0/18:2(9Z,12Z)) | 165 | 77 |
| PAPE | PE (16:0/20:4(5Z,8Z,11Z,14Z)) | 23 | 84 |
| POPE | PE (16:0/18:1(9Z)) | 23 | 98 |
| POPI | PI (16:0/18:1(9Z)) | 0 | 35 |
| PAPS | PS (16:0/20:4(5Z,8Z,11Z,14Z)) | 0 | 77 |
| POPA | PA (16:0/18:1(9Z)) | 0 | 7 |
| SSM | SM (d18:1/18:0) | 83 | 35 |
| NSM | SM (d18:1/24:1) | 83 | 35 |
| CMH | GlcCer (d18:1/16:0) | 30 | 0 |
| CHOL | Cholesterol | 278 | 203 |
| TOTAL | | 805 | 700 |

**Supplementary Table 7. Membrane lipid composition.**

| Stage | | Ensemble | Time Step (fs) | Total Time (ns) | Position Restraint Force Constant (kJ/mol·nm$^2$) | | | Dihedral Restraint Force Constant (kJ/mol·nm$^2$) |
|---|---|---|---|---|---|---|---|---|
| | | | | | Protein Backbone | Protein Side Chain | Lipid Head Group | Glycan Ring |
| Equilibration | 1 | NVT | 1 | 1.25 | 4000 | 2000 | 1000 | 1000 |
| | 2 | NVT | 1 | 1.25 | 2000 | 1000 | 400 | 400 |
| | 3 | NPT | 1 | 1.25 | 1000 | 500 | 400 | 200 |
| | 4 | NPT | 2 | 5 | 500 | 200 | 200 | 200 |
| | 5 | NPT | 2 | 5 | 200 | 50 | 40 | 100 |
| | 6 | NPT | 2 | 5 | 50 | N/A | N/A | N/A |
| Production | | NPT | 4 | 1000 | N/A | N/A | N/A | N/A |

**Supplementary Table 8. Simulation settings for the equilibration and production stages.**

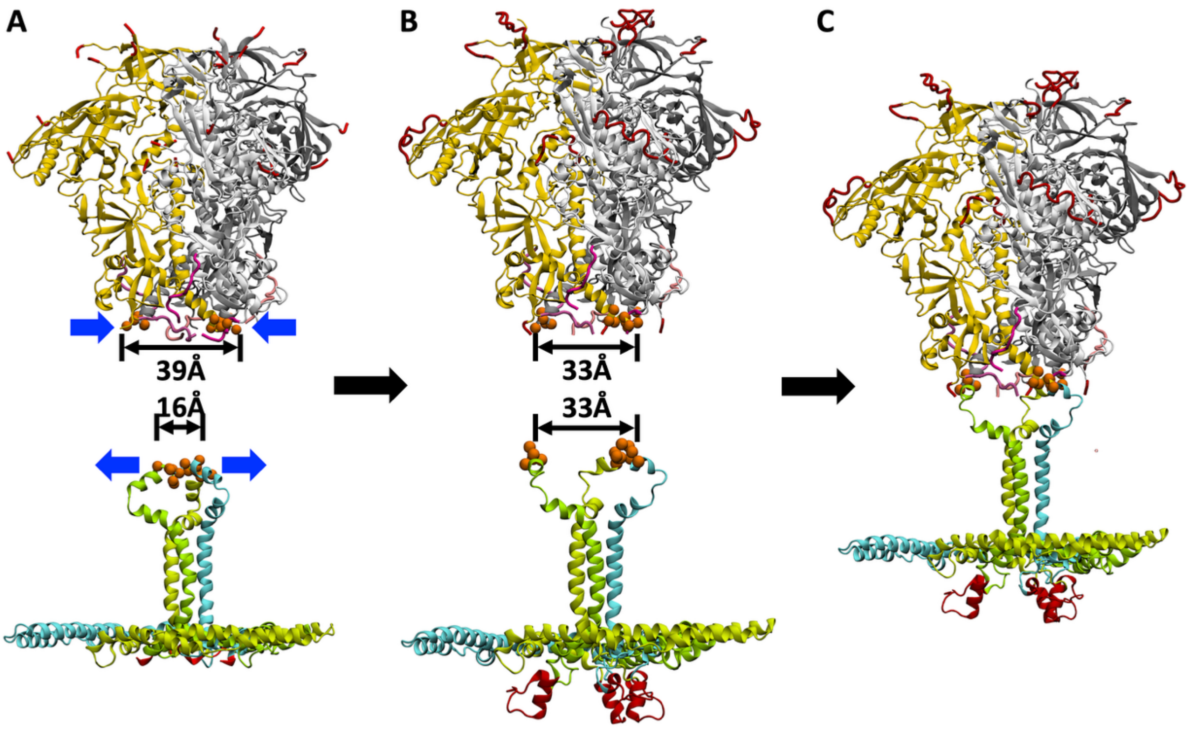


**Supplementary Figure 1. Steps in combining the structure of the ectodomain with the remaining domains.** (**A**) The original PDB structures (PDB IDs: 6B0N and 7LOI) with the common residues highlighted by orange spheres. The distance between the Cα of D664 is 39 Å in 6B0N and 16 Å in 7LOI. The broken ends of missing loops are shown in red, and the cleavage site in magenta. (**B**) Missing loops were grafted from a modeled structure generated by I-TASSER (red). The extra Gly and Ser residues from the 2x$G_4$S linker introduced at the cleavage site in 6B0N were removed (magenta), and MD simulations were performed to adjust distances between the common residues in two PDB structures. (**C**) The common residues were aligned, and two structures were merged.

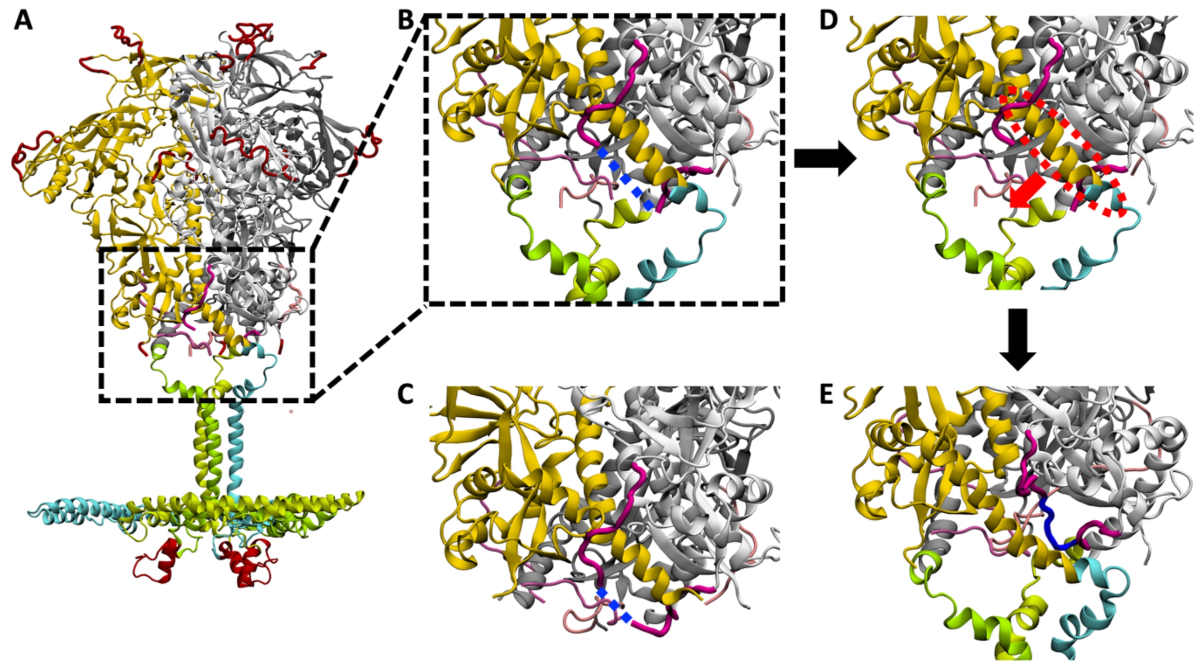


**Supplementary Figure 2. Steps in building the uncleaved model.** (**A**) The cleaved model. (**B**) The cleavage site with the flanking loops in one promoter highlighted in dark magenta. If the flanking loops are connected along the blue dashed line, the protomer shown in yellow and cyan passes through the loop formed by the neighboring protomer in white, resulting in an entangled, knot-like topology. (**C**) The gap between the flanking loops is shorter in PDB structure 6B0N due to the presence of a 10-residue linker $^{508}$GGGGSGGGGS$^{511}$, where the underlined residues are missing. (**D**) MD simulations were performed to adjust the length of the gap between the flanking loops to match that of the $^{508}$REKR$^{511}$ cleavage site, and to reposition the flanking loops and the HR2 helix (red circle) to prevent the connected loop from forming a knot-like structure with the neighboring protomer. (**E**) $^{508}$REKR$^{511}$ (blue) was modeled to connect the flanking loops.
